# Single-nuclei RNA-sequencing uncovers sexually divergent exercise signatures partially mimicked by TFEB overexpression in mouse skeletal muscle

**DOI:** 10.64898/2026.01.23.700915

**Authors:** Katrina Granger, Kailin Liu, Tiarra Joseph, Ari Adler, Ian Matthews, Jocelyne Leon, Clayton Baker, Minhoo Kim, Hu Wang, Xianlin Han, Bérénice A. Benayoun, Constanza J. Cortes

## Abstract

Exercise induces extensive, cell–type–specific transcriptional remodeling in skeletal muscle to support metabolic flexibility and adaptation. However, the regulatory mechanisms underlying these transcriptional programs, and the extent to which they differ between sexes, remain poorly defined. We previously reported that lifelong, muscle-specific overexpression of human Transcription Factor E-B (cTFEB;HSACre transgenic mice) recapitulates many adaptive features of endurance training in both sexes, leading to profound geroprotective effects during aging even in the absence of exercise. Here, we profile transcriptional adaptations to voluntary wheel running (VWR) and TFEB-overexpression at single-nucleus resolution in young male and female mouse tibialis anterior muscle. This represents, to our knowledge, the first integrated analysis of exercise and TFEB signaling using sex as a biological variable. Using robust bioinformatic and single-nuclei RNA-sequencing approaches, we profiled six muscle-resident cell populations and uncover previously unrecognized, sex-dependent signaling nodes governing exercise-associated metabolic plasticity. TFEB activation and endurance training by VWR elicit strongly correlated transcriptional programs enriched for lipid metabolism, mitochondrial remodeling, and immune modulation, establishing TFEB-overexpression as a partial exercise mimetic. In general, female muscle exhibited enhanced extracellular matrix and lipid-associated responses to endurance training and TFEB overexpression, whereas males preferentially engaged in angiogenic and oxidative networks, revealing distinct sex-specific, sex-dimorphic, or sex-agnostic regulatory routes to metabolic flexibility. Integration with independent multi-omics datasets from endurance-trained rats (MoTrPAC) confirms the conservation of TFEB–exercise transcriptional convergence in skeletal muscle across species and potentially muscle types. Together, these findings define TFEB as a regulator of exercise transcriptional programs and reveal sex-specific molecular frameworks that drive metabolic adaptation in skeletal muscle. Furthermore, the resulting sex-resolved, single-nucleus transcriptional atlas provides a unique resource for the field, enabling comparative, mechanistic, and hypothesis-driven exploration of exercise-responsive skeletal muscle regulatory networks across sexes.

## INTRODUCTION

Endurance exercise (running) is a potent physiological stimulus that induces significant structural and functional remodeling in skeletal muscle, including enhanced mitochondrial biogenesis, improved metabolic efficiency, and reinforced neuromuscular integrity ^1–4^. These exercise-induced adaptations contribute not only to improved physical performance but, if performed over the long term, also confer a myriad of health benefits, contributing to reduced susceptibility to age-associated chronic diseases, attenuation of age-associated functional decline, and preservation of muscle mass and function ^5^. Substantial progress has been achieved towards profiling the transcriptional diversity of skeletal muscle responses to aging, exercise, and disease in humans ^6,7^ and rodents ^4,8,9^. However, until very recently, these efforts have largely emphasized male physiology, thereby creating a significant gap in our understanding of skeletal muscle responses to endurance exercise training in females. Indeed, the complexity of the underlying skeletal muscle molecular responses to exercise is multifactorial and is often obscured by both cellular heterogeneity and sex-specific variations ^4,10,11^. The recent release of the Molecular Transducers of Physical Activity Consortium (MoTrPAC) multi-omic and multi-organ atlas of endurance-trained young rats provided a valuable resource to address this complexity and set the stage to begin unraveling the diverse molecular mechanisms underlying skeletal muscle adaptations to exercise across sexes and ages. Despite all of this progress, the cell-type-specific transcriptional responses to endurance training, and how biological sex may impact these profiles, remain unresolved. Thus, there is a critical need to build skeletal muscle cell-type specific transcriptional profiles associated with endurance exercise in a sex-informed manner, to ultimately accelerate the identification of molecular transducers of exercise that could be targeted for individualized therapeutic interventions ^4^.

Transcription Factor E-B (TFEB), a master regulator of mitochondrial function, the endolysosomal network ^12–14^ and lipid metabolism ^15–17^, is activated in skeletal muscle during exercise ^18^ and is required for the full manifestation of running-induced muscle metabolic remodeling ^12^. We have previously shown that muscle-specific human TFEB overexpression across the lifespan (cTFEB;HSACre transgenic mice) mimics key benefits of endurance training on skeletal muscle of both sexes, including fiber-type switching, enhanced oxidative capacity and mitochondrial function, increased exercise endurance, and preservation of muscle mass with aging ^19^. Proteomic profiling of TFEB-overexpressing muscle revealed enrichment of key networks driving enhanced metabolic signaling, including thermogenesis, oxidative phosphorylation, and fatty acid and amino acid metabolism ^19^. We also demonstrated that TFEB overexpression promotes secretion of exercise-responsive, muscle-derived factors into circulation (‘exerkines’) even in the absence of exercise interventions ^19,20^. Recently, we showed that TFEB overexpression also recapitulates exercise-associated cytokine signatures in skeletal muscle, with cTFEB;HSACre muscle displaying features of both early (pro-inflammatory) and late (anti-inflammatory) phases of the exercise-induced immune response ^21^, all profiles reminiscent of those seen after four weeks of voluntary wheel running ^5^. We were also the first to characterize all of these exercise-like effects across both sexes and have uncovered convergent sex- and age-dependent responses in skeletal muscle to both running and TFEB overexpression ^19,21^. Collectively, our previous work suggests that TFEB overexpression may prime muscle into an intermediate, exercise-like state ^21^, positioning the cTFEB;HSACre model as a powerful, orthogonal platform to identify and validate functionally relevant nodes of exercise-associated signaling across sexes. However, to date, there has been no comprehensive analysis of the effects of TFEB-overexpression across skeletal muscle cell types.

To address these multiple gaps in knowledge, we performed single-nuclei RNA sequencing of skeletal muscle from young male and female mice from (i) control, sedentary mice, (ii) mice subjected to voluntary wheel running for 4 weeks, and (ii) mice with skeletal muscle–specific TFEB overexpression. Leveraging our rich dataset, we defined cell-type– and sex-specific transcriptional programs underpinning endurance exercise adaptations, and uncovered extensive transcriptomic remodeling across myonuclear populations, revealing shared and sex-divergent gene expression signatures between endurance training and TFEB activation. These included mitochondrial biogenesis, lipid remodeling, and extracellular matrix reorganization. Furthermore, our work establishes TFEB overexpression as a sexually dimorphic mimic of exercise-induced skeletal muscle plasticity, with converging transcriptional activation of pathways also seen in endurance trained muscle. Integrative transcriptomic, proteomics and metabolomic cross-species validation via the MoTrPAC Datahub confirmed conserved modulation of these pathways by endurance training in the skeletal muscle of young rats. Overall, this work provides a valuable resource to the field and underscores the need for sex-informed exercise research to optimize exercise-mediated metabolic interventions.

## RESULTS

### A single-nuclei transcriptional atlas of young female and male mouse skeletal muscle from sedentary, runner, and cTFEB;HSACre mice

To evaluate sex- and TFEB-dependent transcriptional programs associated with exercise, we performed single-nuclei RNA-sequencing (snRNA-seq) on flash-frozen tibialis anterior (TA) muscle from 6-month-old male and female mice across three groups: (i) non-transgenic sedentary, (ii) non-transgenic voluntary wheel runners (VWR), and (iii) cTFEB;HSACre sedentary transgenics (n=4 animals/group, littermates) (**Figure 1A**). The TA was selected due to its robust response to running ^22,23^, age-related susceptibility to atrophy and dysfunction ^5,24–26^, high TFEB expression in the cTFEB;HSACre model ^19^, and its suitability for snRNA-seq given its high nuclei yield ^8,27–29^ and defined and homogenous fiber-type composition (∼10% type IIa, ∼60% type IIb, ∼30% type IIx with almost no detectable type I fibers in rodents) ^26^.

**Figure 1:**
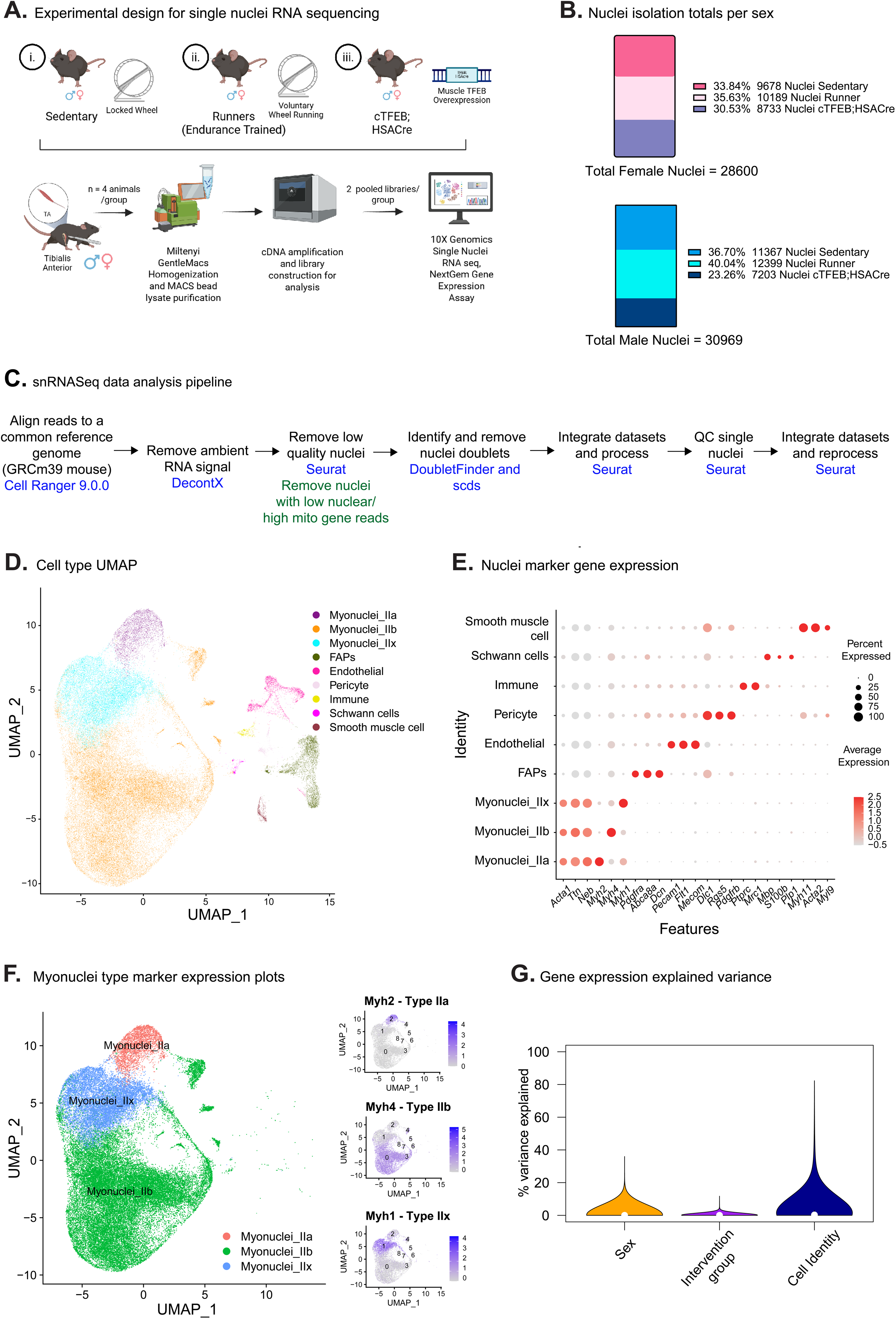
Sex-resolved single-nucleus transcriptional atlas of exercise and TFEB activation in mouse TA muscle. **(A)** Single nuclei RNA-Sequencing 3′ scRNA-seq (10× Chromium v2 and v3) was performed on dissociated TA muscles from sedentary (n=4 M/F), runner (n=4 M/F), and cTFEB;HSACre transgenic (n=4 M/F) mice. **(B)** Fraction of nuclei contributions from each sex group. **(C)** Single-nuclei RNA-Sequencing pipeline for analysis. **(D)** Uniform manifold approximation and projection (UMAP) representations of the final nuclei dataset. Cells colored by manually assigned cell-type IDs based on the expression of hallmark skeletal muscle genes shown in **(E)**. **(F)** UMAP representations of the final dataset for myonuclei only, highlighting nuclei clusters for IIa (Myh2+), IIb (Myh4+), and IIx (Myh1+) myonuclei on the right. **(G)** Variance in gene expression explained by intervention, sex, and cell identity across samples. Cell identity explains more of the variance in gene expression relative to sex or intervention.

Mice in the VWR group had individual 24-hour access to a running wheel for 4 weeks; sedentary mice were housed with nonfunctional, locked wheels (**Figure 1A**). This voluntary running paradigm promotes key muscle metabolic and functional adaptations, including mitochondrial activation ^30,31^, increased capillary density ^32^, and fiber-type switching ^32,33^, while also avoiding stress-related activation of inflammation sometimes seen in forced treadmill interventions ^34,35^. We confirmed continuous running behavior throughout the study for both sexes (**Supplementary Figure S1A**), with females running significantly longer total distances than male littermates (**Supplementary Figure S1B**). In our hands, four weeks of VWR did not reduce total body weight in young male or female runners compared to their sedentary littermates (**Supplementary Figure S1C**). As previously reported ^19^, male cTFEB;HSACre transgenic mice did not exhibit significant differences in total body weight relative to their sedentary littermate controls (**Supplementary Figure S1C**) at this age, although female cTFEB;HSACre mice included in this study showed a modest, but statistically significant, increase in total body weight compared to sedentary and runner females (**Supplementary Figure S1C**).

To comprehensively evaluate transcriptional responses associated with VWR or muscle-TFEB overexpression at a cell-type level and with sex-specific resolution, we generated snRNA-seq datasets of TA skeletal muscle from the groups described above (**Figure 1A**). Using established muscle-specific protocols ^8^, we isolated high-quality nuclei from each sample, and processed them as 2 independent libraries with nuclei pooled from 2 independent animals for each group, together yielding ∼70,000 nuclei before quality filtering (total of 4 animals per condition, **Figure 1B** and see **Methods**). We then applied stringent quality control pipelines (**Figure 1C**) to only retain high-quality singlet nuclei, resulting in a final dataset of 59,569 high-quality nuclei across our 12 libraries (**Figure 1B-C**; see **Methods**). The total number of nuclei contributions was relatively comparable between experimental cohorts (∼23-40% from each sex-matched sedentary, runners, and cTFEB;HSACre group) and between sexes (28,600 total female nuclei and 30,969 total male nuclei, **Figure 1B**). We confirmed that human *TFEB* transgene expression was detected exclusively in libraries derived from cTFEB;HSACre mice, and confirmed our previous reports of higher TFEB expression in male cTFEB;HSACre skeletal muscle relative to their female transgenic littermates ^19^ (**Supplementary Figure S1D**). We also confirmed the sex identity of all processed samples by expression of sex-specific genes *Xist* (X-inactive specific transcript; female-specific) in female samples and *Ddx3y* (a Y-linked gene; male-specific) in male samples only (**Supplementary Figure S1E**).

Importantly, we observed good mixing across UMAP space as a function of intervention within each sex (**Supplementary Figure S1F-H**). However, cells from females and males separate in UMAP space, suggestive of either strong sex-dimorphism in single cell transcriptomes or potentially due to separate sex batch processing of animals of each sex (**Supplementary Figure S1F**). To annotate single nuclei transcriptomes to cognate cell types, we leveraged a semi-supervised annotation approach combining (i) unsupervised clustering, (ii) marker-based annotation using markers from singleCellBase ^36^ and scMayoMap ^9^, and (iii) reference-based annotation using a mouse snRNA-seq muscle atlas ^37^ (**Supplementary Figure S1I,** see **Methods**). Using this approach and well-established markers for muscle cell types, we annotated nine nuclei populations, including expected myonuclei subtypes (IIa, IIb, IIx), fibrogenic adipose progenitors (FAPs), endothelial cells (ECs), smooth muscle cells (SMCs), pericytes, immune cells, and Schwann cells (**Figure 1D-E**). As expected, the most abundant nuclei type annotated was myonuclei IIb (*Myh4*+), followed by myonuclei IIx (*Myh1*+) and then myonuclei IIa (*Myh2*+) (**Figure 1F**), consistent with TA fibertype composition ^26^ and prior mouse TA snRNA-seq studies ^8,9^. As an important sanity check, cell identity represented the main source of gene expression variation between nuclei, followed by sex (consistent with UMAP clustering observations), then intervention (**Figure 1G**).

Next, we examined how endurance training and TFEB overexpression directly influenced the transcriptional responsiveness and nuclear composition of the TA muscle (**Figure 2A**). Augur analysis ^38,39^ identified myonuclei as the cell type most transcriptionally responsive to these interventions (**Figure 2B**). More specifically, myonuclei IIa displayed the highest responsiveness in females, whereas myonuclei IIx were the most responsive in males (**Figure 2C**). Endothelial cells also exhibited strong intervention-induced transcriptional changes in both sexes (**Figure 2C**).

**Figure 2:**
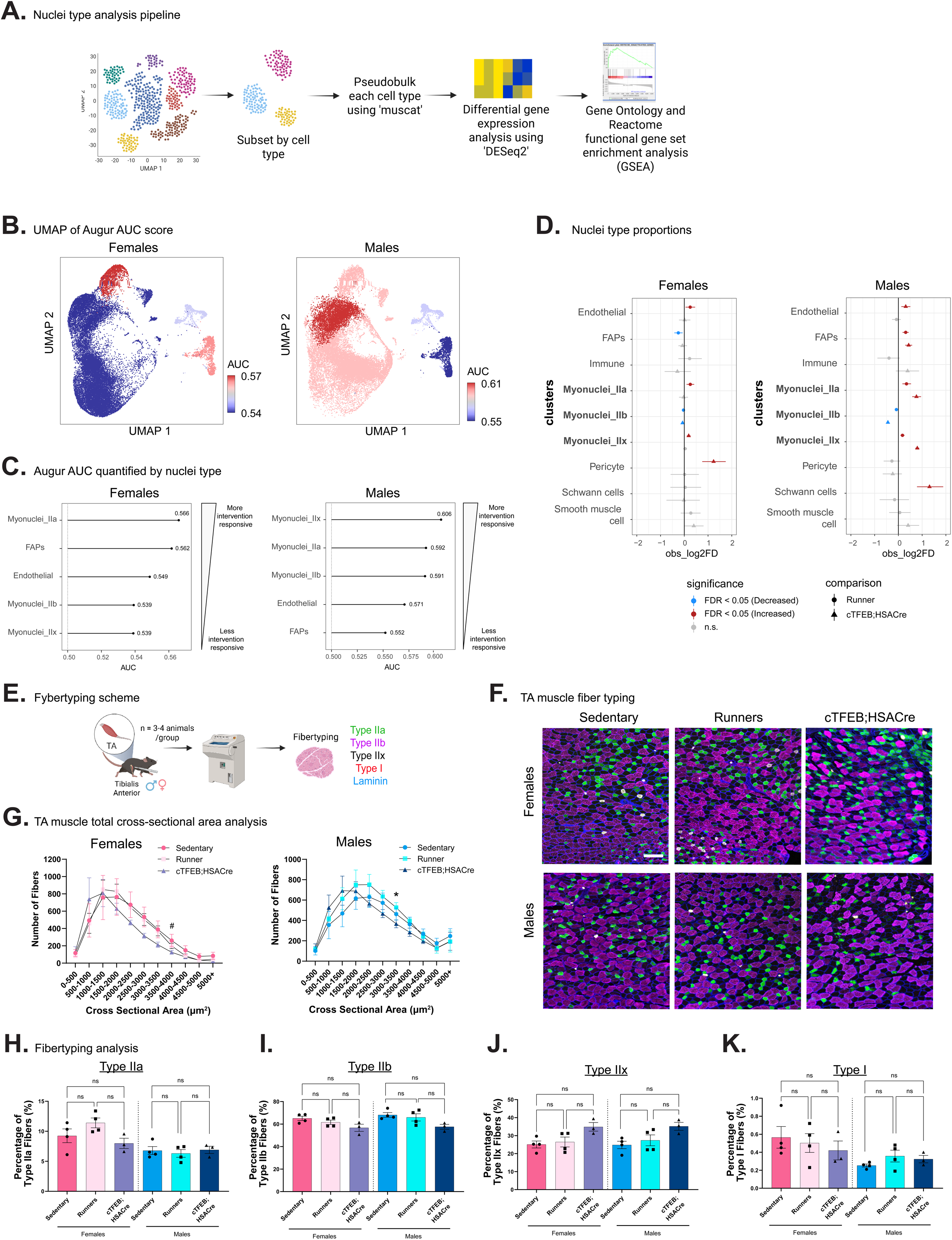
Nuclei type-transcriptional responses associated with running and TFEB-overexpression across sexes are validated by fiber typing analysis. **(A)** Nuclei type analysis scheme. The magnitude of gene expression differences between cell types were analyzed by Augur and cell type proportions were analyzed using scProportionTest. **(B)** UMAP representation of Augur AUC results. Values close to 0.5 denote smaller differences (blue shades), and values closer to 1 reflect larger differences (red shades). **(C)** Lollipop plot of Augur AUC results per nuclei type in either sex. **(D)** Skeletal muscle nuclei type proportion differences between sexes and groups. Blue-coded nuclei types are decreased, and red-coded cell types are more abundant in respective groups relative to sex-matched sedentary controls. Myonuclei are highlighted in bold. **(E)** Fiber typing immunostaining scheme in TA mouse muscle. **(F)** Representative images of fiber typing in TA muscle by sex x intervention. MyHC1 (fiber type I) in red, MyHC2A (fiber type IIa) in green, MyHC2B (fiber type IIb) in magenta, MyHC2X (fiber type IIx) unstained, and laminin in blue. Scale bar = 200 µm. **(G)** Distribution curves of cross-sectional area (CSA) results by sex x intervention. Data is represented as mean ± SD. **(H-K)** Quantification of individual fiber type abundance. Data is represented as mean ± SEM. One-way ANOVA within sex with Kruskal-Wallis post-hoc test. * Statistical difference between sedentary and runner. # Statistical difference between sedentary and cTFEB;HSACre. Lack of annotation indicates comparisons were not significant. Each data point represents the average of three separate images collected from three sections/individual. Statistical tests as listed. N.s: non-significant, * P ≤ 0.05, ** P ≤ 0.01.

We then examined the nuclei composition of each experimental group as a function of sex, using the snRNA-seq-specific scProportionTest tool ^40^, which uses permutation testing to quantify cell type proportion changes between groups (**Figure 2A, D**). We observed significant enrichment of type IIa myonuclei in runner samples from both sexes, as well as in male (but not female) TFEBs. Type IIx myonuclei were increased in TFEB samples from both sexes, as well as in male (but not female) runners. In contrast, type IIb-annotated myonuclei were decreased across both sexes and both interventions (**Figure 2D**). Fiber typing, the immunohistochemical classification of individual myofibers based on myosin heavy chain isoform expression, is a well-established method to determine fiber type size and abundance in intact skeletal muscle (**Figure 2E**). In general, voluntary wheel running exercise in mice generally does not produce major shifts in male TA fiber type composition, although it does induce functional and morphological adaptations in muscle fibers ^22,41^. Consistent with this, fiber typing analysis of whole TA muscle sections (**Figure 2F-K** and **Supplementary Figure S2,** n=3-4/group, independent cohort) revealed no major changes in female TA muscle overall fiber size (**Figure 2G**), and a slight shift toward an increased proportion of medium-sized fibers (cross-sectional area [CSA]: 1,000–3,000 µm²) in males following voluntary running (n=3-4 group, independent cohort, **Figure 2G**). This shift was even more pronounced in TFEB-overexpressing muscles, with both male and female cTFEB;HSACre mice exhibiting an increase in small- to medium-sized fibers (CSA: 500–2,000 µm²) and a concomitant decrease in medium- to large-sized fibers (CSA: 2,000–4,000 µm²) (**Figure 2G**). Quantification of individual fiber type abundance revealed no significant changes in the proportions of myofiber type IIa or type IIb in either runner or cTFEB;HSACre TA muscles of both sexes (**Figure 2H-I**), although there were trends towards a potential increase in type IIa (MyHC-IIa^+^) fibers. Notably, we observed a mild trend toward increased numbers of type IIx (MyHC-IIx^+^) and concomitant decrease in type I (MyHC-I^+^) fibers in TFEB-overexpressing TA muscles in both males and females (**Figure 2J-K**), although this did not reach statistical significance. Fiber type-specific CSA analyses further demonstrated that these changes were predominantly driven by a shift toward smaller fiber size in IIb and IIx myofibers in both male and female mice under both conditions (**Supplementary Figure S2A-B**). Male type IIa fibers were unaffected by either voluntary wheel running or TFEB overexpression, whereas female type IIa fibers appeared to have a mild increase in small-sized fibers (CSA: 500–1,500 µm²) in runners (**Supplementary Figure S2B**). These results are also consistent with our previous fiber typing and CSA analysis of cTFEB;HSACre gastrocnemius muscle of both sexes ^19^, suggesting that TFEB-overexpression increases the abundance of IIa and IIx fast-twitch fibers across multiple muscle types, promoting a runner-like fast-to-slow fiber typing profile ^26,42^. Importantly, these enrichment patterns in fiber typing are in line with those observed in our scProportionTest analysis (**Figure 2D**), suggesting that these changes in nuclei proportions are likely driven by our interventions (and/or sex status) rather than technical variations during nuclei isolation.

Notably, we also observed changes in the abundance of non-myonuclear populations across our samples as assessed by scProportionTest. Specifically, endothelial cell–annotated nuclei were increased in female runner samples (but not in female TFEB samples) and in male TFEB samples (but not in male runner samples) (**Figure 2D**), which we also confirmed via endothelial cell marker staining (see **Figure 7E**). In addition, there was a significant decrease in the number of FAP-annotated nuclei in female runners, whereas this population was increased in both male runners and male TFEB samples (**Figure 2D**). Pericyte-annotated nuclei were specifically increased in female TFEB samples only, while Schwann cell–annotated nuclei were increased exclusively in male TFEB samples (**Figure 2D**). Overall, this suggests intervention- and sex-specific effects on the relative abundance or representation of particular cell types in TA muscle, promoting shifts in myonuclei populations (fiber type switching) and non-myonuclei populations that may underlie some of the metabolic plasticity associated with running and TFEB-overexpression.

### Differential expression analysis reveals cell type-specific responses to exercise and *TFEB* overexpression

Next, we evaluated differential gene expression profiles as a response to our interventions in each sex across cell types. To avoid inherent statistical issues related to single-cell/nuclei level differential expression analysis ^43^, we decided to leverage a robust, stringent pseudobulk-level differential gene expression approach (see **Figure 3A** and **Methods**). To limit the influence of technical noise and improve interpretability, only cell types with at least 20 nuclei in every snRNA-seq library were analyzed further, yielding 6 cell types for analyses (i.e. Myonuclei IIa, Myonuclei IIb, Myonuclei IIx, Endothelial cell nuclei, smooth muscle cell nuclei, and FAP nuclei; **Figure 3**). First, we performed multi-dimensional scaling (MDS) to visualize the overall transcriptomic landscape similarity across nuclei types. Importantly, replicate libraries were clearly separated by treatment groups (sedentary/runner/transgenic status) within each sex across all cell types (**Supplementary Figure S3A-F**), consistent with the fact that our interventions have a larger effect on gene expression than isolation variability. Intriguingly, in most cases, one could visualize an axis of separation in MDS space between sedentary animals on the one hand vs. exercised and TFEB transgenic animals on the other hand, suggesting that there are common transcriptional shifts to muscle cell type components in response to exercise and TFEB overexpression (**Supplementary Figure S3A-F**).

**Figure 3:**
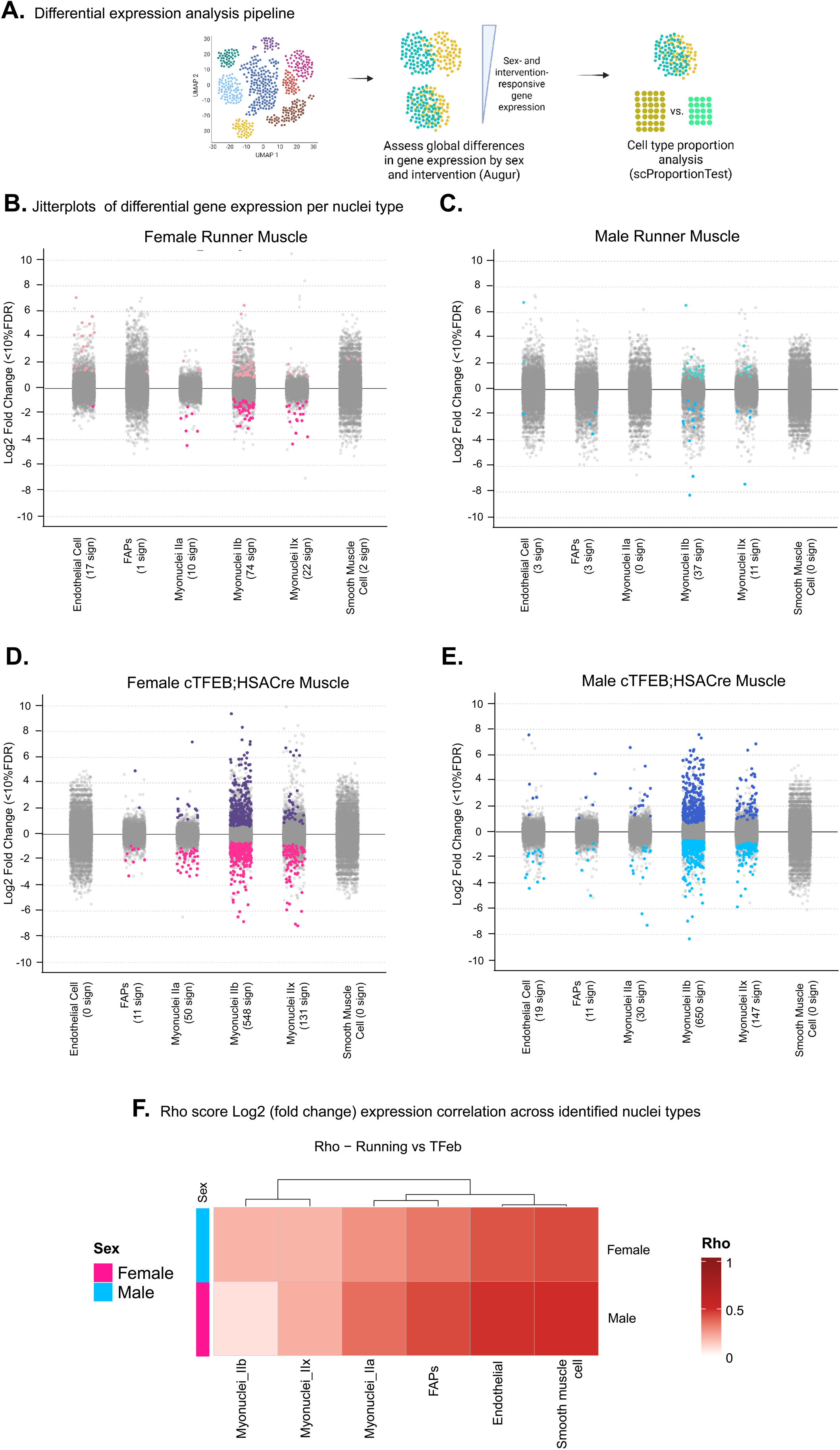
Differential gene expression analysis reveals shared transcriptional patterns between endurance-trained and TFEB-overexpressing skeletal muscle. **(A)** Differential expression analysis scheme. **(B)** Jitter plot of differentially expressed genes per nuclei type (FDR < 10%) per condition for each sex, with the number of significantly differential genes in parentheses. Differentially expressed transcripts are highlighted in their corresponding colors (pink hues for females, blue hues for males) while non-significant genes are in gray. Transcripts above the midline are up-regulated while transcripts below the midline are down-regulated. TFEB overexpression drives more transcriptional changes than endurance training. **(C)** Rho correlation heatmap of gene expression log2 fold change in runner vs. TFEB nuclei.

Interestingly, DESeq2 differential expression analysis revealed a wide range of responsiveness to exercise or TFEB overexpression across cell types, with myonuclei showing the highest numbers of differentially expressed genes and smooth muscle cells showing no significant genes, regardless of sex (**Figure 3B-E**). Endothelial cells also showed robust differential expression, although to a lesser extent than myonuclei, consistent with our Augur results (**Figure 2B-C**). In general, nuclei from cTFEB;HSACre muscle (**Figure 3D-E**) showed more transcriptional changes than those from sex-matched runners (**Figure 3B-C**) compared to their cognate sedentary wild-type controls, consistent with the role of TFEB as a transcription factor with high activity in myonuclei. The scope of these transcriptional changes is consistent with previous reports in both human and murine skeletal muscle snRNAseq studies ^6,8,28^.

We next tested the hypothesis that TFEB overexpression alters gene expression in the TA muscle by engaging genetic programs shared with endurance running across cell types and between sexes. For this purpose, we used Spearman’s rank correlation (which ranges from -1 to +1 and measures the strength and direction of the monotonic relationship between two gene sets) to compare global transcriptional profiles between runner/sedentary and cTFEB;HSACre/sedentary profiles across robustly detected cell types (**Figure 3F and Supplementary Figure S3G-L**). We found significant positive correlations in gene expression remodeling between runner and TFEB-muscle in all tested cell types (significance of correlation p < 9.18E-19), suggesting that TFEB overexpression indeed acts like an exercise mimetic. This was strongest in myonuclei subtypes, including IIa (Females Rho: 0.275, Male Rho: 0.359), IIb (F Rho: 0.19, M Rho: 0.0781), and IIx (F Rho: 0.186, M Rho: 0.202) (**Figure 3C**, and **Supplementary Figure S3G, I, K)**, consistent with our prior findings of exercise-like remodeling in TFEB-overexpressing muscle ^19^. To note, we also observed a strong transcriptional concordance in endothelial cell nuclei from runner and TFEB-overexpressing muscle (**Figure 3C**, and **Supplementary Figure S3H**).

### Functional enrichment analysis reveals that TFEB Overexpression and Exercise Induce Sex-Dimorphic, Myonuclear Subtype–Specific Transcriptional Programs in Skeletal Muscle

Next, we used our differential expression analysis results to identify functional processes regulated by running and/or cTFEB;HSACre expression across cell types. Importantly, for increased robustness, we focused our analyses hereafter on cell types with robust differentially expressed genes based on our analyses above (**Figure 3** and **Supplementary Figure S3**): myonuclei IIa, IIb, IIx, and endothelial cells. To investigate the potential functional consequences of running and/or cTFEB;HSACre expression across cell types, we utilized Gene Set Enrichment Analysis (GSEA) ^44^ together with gene sets defined by Gene Ontology (GO) and Reactome (**Figures 4-7**; see **Methods**). Using this approach, we identified hundreds of gene sets significantly regulated compared to sedentary controls in runner nuclei, cTFEB;HSACre nuclei, or both. GO GSEA analysis revealed many biological processes regulated by exercise were cell-type, sex- and intervention-specific, while others were shared across conditions. In addition to well-known responders to endurance training, our robust, sex-informed dataset served as a discovery tool, uncovering multiple previously unknown transcriptional nodes regulating skeletal muscle exercise plasticity. Furthermore, we also found that: a) running and TFEB overexpression shared significantly enriched GO terms, b) many of these terms exhibited sex-dimorphic, sex-specific, or sex-shared expression patterns, c) these sex-associated patterns were largely preserved across both interventions, and d) were validated in independent rat (MoTrPAC) transcriptional, proteomics, and metabolomics datasets, suggesting that these pathways are conserved across species. We describe our results below.

**Figure 4:**
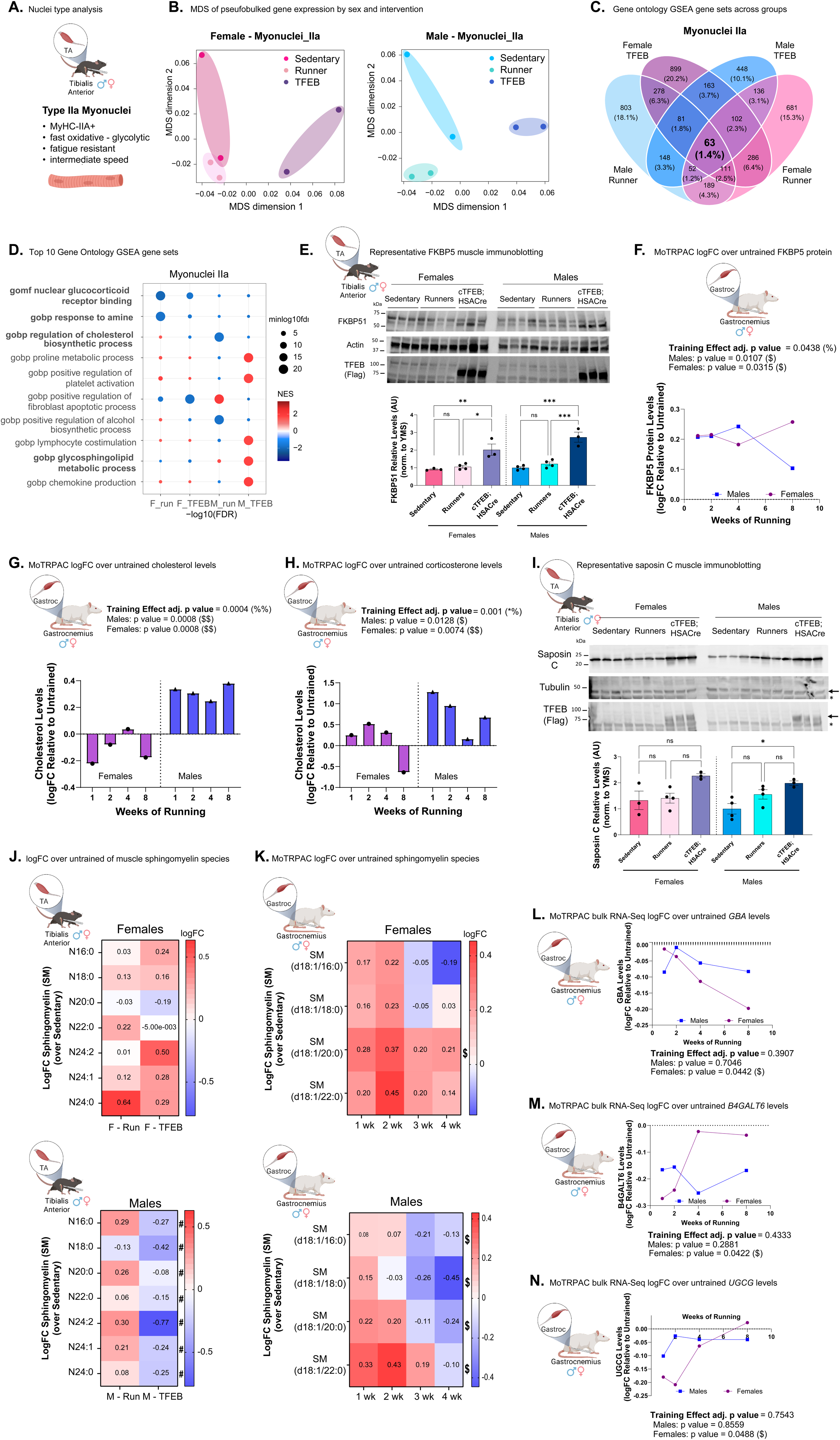
Conserved sex-dimorphic lipid-associated signatures in endurance-trained and TFEB-overexpressing IIa mouse myonuclei are also recapitulated in endurance-trained rats. **(A)** Type IIa myonuclei type functional characteristics. **(B)** MDS plots for myonuclei IIa in both sexes. See also **Supplementary Figure S3**. **(C)** Shared and unique GO terms (Biological Process, Cell Compartment, Molecular Function) enriched by GSEA in IIa myonuclei across all four conditions. The percentage of overlapping pathways is in parentheses. **(D)** Top ten GO terms enriched in at least one condition, highlighting shared and divergent transcriptional signatures across sex and intervention. Note the enrichment for lipid-associated pathways in bold. **(E)** Immunoblot analysis for glucocorticoid signaling-inhibitor FKBP51 from sedentary, runner, and cTFEB;HSACre tibialis anterior lysates from an additional cohort mice (n=3-4/group). Marker densitometry quantification relative to actin (loading control) below shows increased abundance of FKBP51 in TA from cTFEB;HSACre mice of both sexes. One-way ANOVA within sex with post-hoc test. **(F)** Increased abundance of FKBP5 protein in the gastrocnemius muscle of both male and female endurance-trained rats over sedentary controls (MoTrPAC DataHub). **(G-H)** Unbiased metabolomics analysis showing significant sex-dimorphic and temporal offsets effects for endurance training on levels of **(G)** cholesterol and cholesterol-metabolic product **(H)** corticosterone in rat skeletal muscle (MoTrPAC DataHub). **(I)** Immunoblot analysis for glycosphingolipid metabolism coactivator saposin C from same cohort from **(E)**. Marker densitometry quantification relative to tubulin (loading control) shows increased abundance of Saposin C in TA from male runner and cTFEB;HSACre mice of both sexes. One-way ANOVA within sex with post-hoc test. **(J)** Targeted lipidomics of mouse tibialis anterior muscle showing mild trends towards increased (both female groups, and male runners) and decreased (male TFEBs) levels of detected sphingomyelin (SM) species. **(K)** MoTrPAC DataHub targeted analyses of sphingomyelin (SM) levels confirms sex-dimorphic SM adaptations to endurance training in female (increased) and male (decreased) gastrocnemius muscle. All MoTrPAC results and lipid levels reported as logFC over sex-matched sedentary controls. **(L-N)** Targeted bulk RNA-Seq analysis confirms female-specific transcriptional downregulation of glycosphingolipid enzymes glucocerebrosidase (*GBA*) **(H)**, beta-1,4-galactosyltransferase 6 (*B4GATL6*) **(I)**, and UDP-glucose ceramide glucosyltransferase (*UGCG*) **(J)**, in gastrocnemius muscle from endurance-trained rats (MoTrPAC DataHub). * Statistical difference between sedentary and runner mouse muscle. # Statistical difference between runner and cTFEB;HSACre mouse samples, one-way ANOVA with post-hoc test. % Statistical difference of endurance training as reported by MoTrPAC. $ Statistical difference between trained and endurance groups within each sex as reported by MoTrPAC. N.s: non-significant, * P ≤ 0.05, ** P ≤ 0.01. *** P ≤ 0.001 Lack of annotation indicates comparisons were not significant. Data is represented as mean ± SEM unless otherwise noted.

#### IIa Myonuclei

Type IIa myonuclei are mostly considered to encode for fast-twitch and mixed (oxidative/glycolytic) myofibers, usually fatigue resistant and with intermediate contraction speed (**Figure 4A**). Multi-dimensional scaling (MDS) analysis of IIa nuclei transcriptional profiles confirmed that replicates clustered tightly by sex, with sedentary and runner samples clearly separating (**Figure 4B**). Interestingly, TFEB-expressing samples consistently separated between sedentary/runner clusters (**Figure 4B**), suggesting TFEB-overexpression may drive an intermediate transcriptional state. Consistent with this hypothesis, functional enrichment analysis using GSEA for Gene Ontology (GO) terms identified a myriad of significantly differentially expressed gene sets associated with Biological Process (BP), Cell Compartment (CC), and Molecular Function (MF) (F/R: 1,619 total; F/TFEB: 1,983 total; M/R: 1,724 total; M/TFEB: 1,192 total, see **Supplementary Table I**). The top five up- and down-regulated gene sets in each condition are presented in **Supplementary Figure S4A**. We identified sex-specific and sex-agnostic transcriptional responses to exercise and TFEB-overexpression, with 323 (∼12%) GO gene sets shared between male and female runners, and 320 (∼15%) shared between TFEB-expressing cohorts of both sexes (**Figure 4C**). Comparable overlaps were observed between experimental conditions, with 428 GO categories (∼19%) shared between female runner and female TFEB groups, and 250 (∼13%) shared between male runner and male TFEB IIa myonuclei (**Supplementary Figure S4B-C**). The four-way intersection across groups of significantly enriched GO categories highlighted 63 categories that were common to both sexes and both interventions (**Figure 4C**), suggesting the existence of core transcriptional signatures associated with exercise- and exercise-like induced remodeling of type IIa fibers. Among these, gene sets related to immune responses and inflammation (14/63), as well as lipid metabolism (9/63) were enriched. We highlight the top 10 of these 63 shared categories in **Figure 4D**, focusing on previously undescribed regulatory nodes of transcriptional regulation of exercise-associated plasticity.

For example, both runner and cTFEB;HSACre samples from both sexes exhibited a previously uncharacterized transcriptional downregulation of the GO Molecular Function for *nuclear glucocorticoid receptor binding* (**Figure 4D**). This effect was particularly strong in female samples, but was still significant in male runner and cTFEB;HSACre IIa mynuclei as well (**Figure 4D**). We confirmed trends towards reduced abundance of glucocorticoid receptor protein (**Supplementary Figure S4E**) and increased abundance of FK506-Binding Protein 51 (FKBP51, a negative regulator of glucocorticoid signaling) protein in skeletal muscle lysates from cTFEB;HSACre mice of both sexes (**Figure 4E**, n=3-4 group, independent cohort). Consistent with these findings in, targeted analysis of gastrocnemius muscle bulk RNA-Seq datasets from endurance trained young rats (MoTrPAC) ^4^ confirmed an endurance training effect for the downregulation of *NR3C1/GR* (the gene encoding the glucocorticoid receptor) in male (but not female) runner rat muscle (**Supplementary Figure S4F**), and a significant endurance training effect for the upregulation of glucocorticoid receptor-inhibitor *FKBP51* mRNA levels in both sexes (**Figure 4F**). This suggests that exercise (and/or TFEB-overexpression) may suppress canonical GR target gene signatures by promoting FKBP51-mediated GR sequestration.

Another down-regulated GO Biological Process gene set in IIa myonuclei shared across all samples was *response to amine biological processes,* as well as a conserved transcriptional upregulation of *proline metabolic process* in female (but not male) runners and cTFEB:HSACre IIa myonuclei of both sexes (**Figure 4D**). Supporting this observation, we have previously demonstrated via proteomics analysis that TFEB overexpression drives substantial remodeling of amino acid metabolism pathways of male cTFEB;HSACre quadriceps muscle, another fast-twitch muscle enriched for IIa myofibers ^19^. This included increased degradation of branched-chain amino acids (BCAAs; leucine, isoleucine, valine) and altered metabolism of several non-BCAAs, such as β-alanine, tryptophan, alanine, aspartate, glutamate, and arginine, as well as increased proline metabolism ^19^, consistent with our transcriptional results presented here.

Sex-dimorphic responses to running were also prevalent in these transcriptional signatures. Female-specific gene sets included upregulation of GO Biological Processes related to *platelet activation, lymphocyte stimulation,* and *chemokine production*. Notably, these gene sets were also upregulated in male and female cTFEB;HSACre (but not male runner) IIa myonuclei, despite the absence of exercise in this model, highlighting the convergence of TFEB overexpression and VWR (**Figure 4D**). Importantly, we have recently demonstrated that muscle-TFEB overexpressing muscle displays a similar, sex-specific, immune-signaling cytokine profile to that seen in endurance trained muscle, including multiple interleukins, chemokines, and other immune system-associated cytokines ^21^. By contrast, the GO Biological Process *regulation of cholesterol biosynthetic process* was significantly downregulated in males under both conditions (and female TFEB muscle) but upregulated in female runners (**Figure 4D**). Interrogation of existing MoTrPAC gastrocnemius muscle metabolomic profiles confirmed that 8 weeks of endurance training increases levels of cholesterol (**Figure 4G**) and decreases levels of corticosterone (a cholesterol precursor) (**Figure 4H**) in male skeletal muscle, with a much milder or no effect in female skeletal muscle, confirming a sex-dimorphic effect of endurance training on muscle cholesterol metabolism.

Many of the top categories found using this approach also indicated changes in lipid metabolism in response to endurance training or TFEB overexpression in IIa myonuclei (**Figure 4D**). For example, the GO Biological Process g*lycosphingolipid metabolic process* was clearly sex-dimorphic, being downregulated in female and upregulated in male IIa myonuclei (**Figure 4D**). This enrichment for lipid signaling terms was also found via Reactome analysis, which identified an upregulation of *phospholipids in phagocytosis*, *synthesis of prostaglandins and thromboxanes,* and *vitamin B12 transport and metabolism*, in runners and cTFEB;HSACre IIa myonuclei of both sexes (**Supplementary Figure S4D**), in agreement with the proposed lipid-associated anti-inflammatory effects of endurance training ^2,45^. We confirmed elevated saposin C levels (a small lysosomal activator protein that facilitates glycosphingolipid degradation) in muscle lysates from male groups and female cTFEB;HSACre runners (**Figure 4I**, n=3-4/group, independent cohort), consistent with our transcriptional signatures.

Guided by these results, and to directly interrogate lipid responses to endurance training and TFEB-overexpression we performed targeted lipidomics analysis of TA skeletal muscle from all groups (n=4 group, independent cohort). Ceramide is at the center of sphingolipid metabolism and acts both as a substrate and a precursor for the generation of other sphingolipids, including sphingomyelin ^46^. Although total skeletal muscle ceramide levels were not significantly changed across any of the groups (**Supplementary Figure S4G**), analysis of individual ceramide lipid species revealed trends towards general increases in female runners and female TFEB-muscles (**Supplementary Figure S4H**). In contrast, male runners and TFEB-muscle exhibited minimal ceramide changes, with a trend toward decreased levels across species N16:0 (**Supplementary Figure S4H**). These observations align with similar MoTrPAC metabolomics data showing significant training effects on ceramide levels, with minor ceramide alterations during the early weeks (1 and 2) of endurance training in both sexes, followed by increased ceramide levels from weeks 4 through 8 (**Supplementary Figures S4I**). In contrast, total sphingomyelin levels trended towards a decrease in male TFEB-overexpressing mice, whereas female runner and female TFEB-overexpressing muscle exhibited slight increases (**Supplementary Figure S4J**). Individual sphingomyelin species were increased in runners of both sexes and in female TFEB-muscle (**Figure 4J**), while male TFEB-overexpressing muscle displayed reductions across these same species instead (**Figure 4J**), consistent with the predicted transcriptional upregulation of glysosphingolipid metabolism in this group (**Figure 4D**). These findings align with data from the MoTrPAC consortium, which also found that endurance training has significant effects on sphingomyelin species in the gastrocnemius muscle of endurance-trained rats (**Figure 4K**). Female runner rat muscle progressively increased sphingomyelin levels during endurance training from weeks 1 through 8, especially of the longer chained isoforms (**Figure 4K**), whereas males exhibited a transient increase only during weeks 1 and 2, followed by a decrease from weeks 4 to 8 (**Figure 4K**). This is consistent with male-TFEB overexpressing muscle sharing features of both early and late exercise adaptations (see below). Targeted analysis of MoTrPAC gastrocnemius RNA-Seq datasets also confirmed endurance training effects in female (but not male) muscle in enzymes involved in glycosphingolipid degradation (glucocerebrosidase, *GBA*) and synthesis (beta-1,4-galactosyltransferase 6, *B4GALT6*, and UDP-glucose ceramide glucosyltransferase, *UGCG*) (**Figure 4L-N**), suggesting that the transcriptional downregulation of glycosphingolipid metabolism identified via mouse snRNASeq may be a consistent response across murine muscle. Overall, this integrated transcriptional and lipidomic profile highlights sex-specific differences in sphingolipid dynamics during both running endurance adaptations and TFEB modulation, although the physiological consequences of these lipid shifts remain unknown.

#### IIb Myonuclei

Next, we examined transcriptional responses to voluntary running and TFEB overexpression in Type IIb myonuclei, overall considered to encode for very-fast twitch, highly fatigable glycolytic fibers (**Figure 5A**). As with Type IIa myonuclei, Type IIb myonuclei also strongly clustered by intervention in our MDS analysis, with TFEB-expressing samples clustering between sedentary and runner groups for both sexes (**Figure 5B**). We also identified high numbers of significantly differentially expressed GO Phenotest GSEA gene sets (F/R: 2055 total; F/TFEB: 1804 total; M/R: 1651 total; M/TFEB: 1425 total, see **Supplementary Table I**); the top five up-and down-regulated pathways are presented in **Supplementary Figure S5A**. Of these, there were 819 (∼36%) female and 306 (22%) male shared categories between running and TFEB-overexpression GO gene sets (**Supplementary Figure S5B**). Sex-specific and sex-agnostic transcriptional responses to exercise and TFEB were also detected, with 579 (∼24%) GO gene sets shared between male and female runners, and 470 (∼24%) shared between TFEB-expressing cohorts of both sexes (**Figure 5C**). Overlaps were also observed between experimental conditions, with 819 GO Biological Process categories (∼36%) shared between female runner and female TFEB groups, and 306 (∼15%) shared between male runner and male TFEB IIb myonuclei (**Figure 5C**). One hundred and twenty one (121) GO gene set categories were common to both sexes and both interventions (**Figure 5C**). Among these, there was an enrichment for categories related to GO gene sets for *cell adhesion* (13/121), *neuronal signaling and axons* (35/121), and *extracellular matrix* (14/121). We highlight the top 10 of these shared categories in **Figure 5D**. Female runner and cTFEB;HSACre IIb myonuclei samples demonstrated complete concordance in the transcriptional upregulation of the top-enriched gene sets (10 out of 10 top GO terms). In contrast, male cTFEB;HSACre IIb myonuclei exhibited only partial overlap, sharing differentially expressed features of both the female and male runner transcriptional profiles (**Figure 5D**).

**Figure 5:**
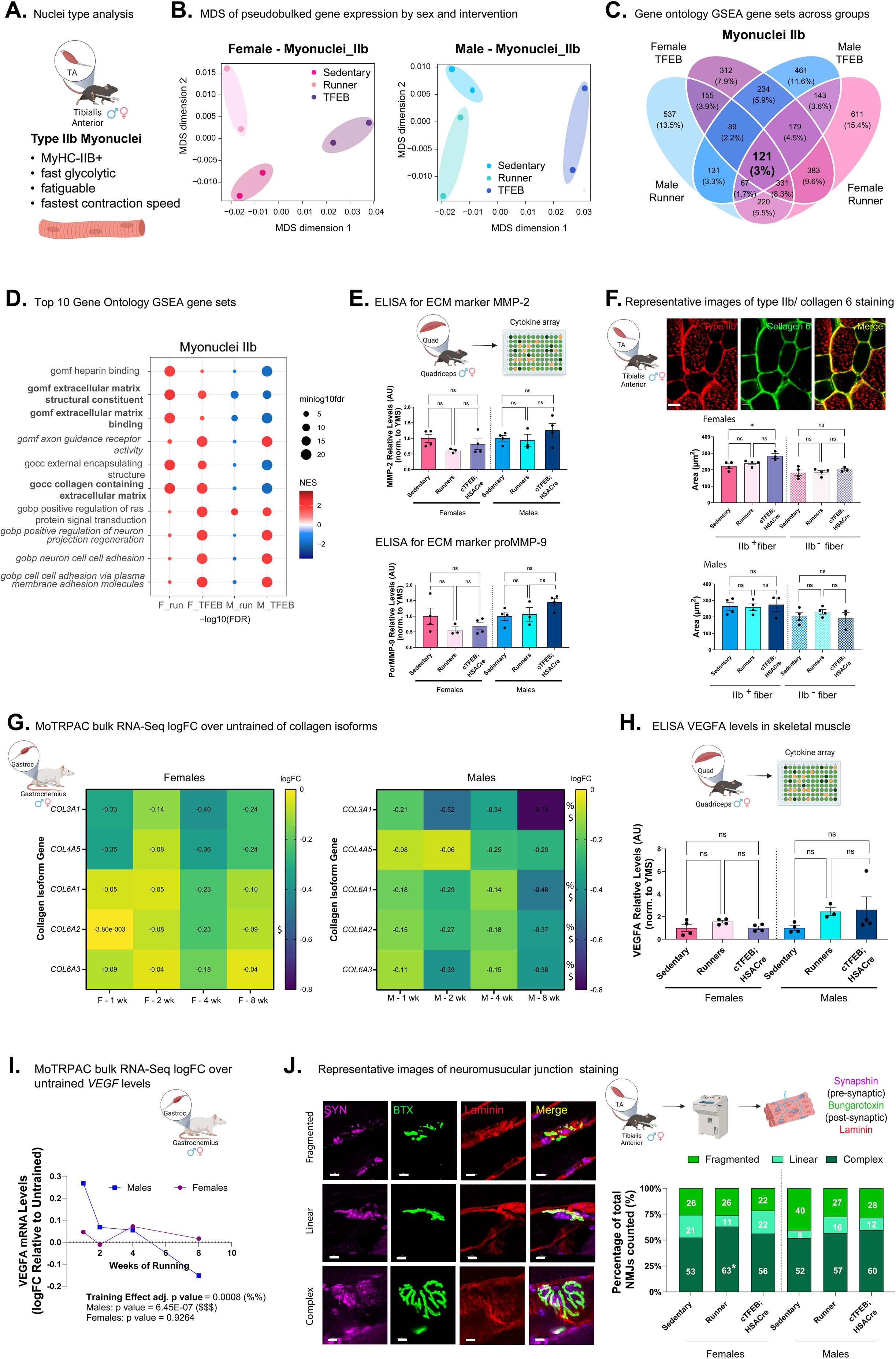
Sex-dimorphic extracellular matrix and neuromuscular junction remodeling in type IIb myonuclei and fibers. **(A)** Type IIb myonuclei type functional characteristics. **(B)** MDS plots for myonuclei IIb in both sexes. See also **Supplementary Figure S3**. **(C)** Shared and unique GO terms (BP, CC, MF) enriched by GSEA in IIb myonuclei across all four conditions. The percentage of overlapping pathways is in parentheses. **(D)** Top ten GO terms enriched in at least one condition, highlighting shared and divergent transcriptional signatures across sex and intervention. Note the enrichment for extracellular matrix-associated pathways (in bold) and neuronal signaling (in italics). **(E)** Cytokine array confirms trends towards sex-specific (MMP-2, decreased in females only) and sex-dimorphic (pro-MMP9, decreasing in females, increasing in males) effects of endurance training and TFEB-overexpression in quadriceps muscle lysates. One-way ANOVA within sex with post-hoc test. **(F)** TA cross-sections from sedentary, runner and cTFEB;HSACre mice stained for type IIb fibers (red) and collagen 6 (green) confirms increased levels of collagen 6 in IIb fibers from female (but not male) TFEB-overexpressing muscle. One-way ANOVA within sex/fiber type with post-hoc test. Scale bars = 30 µm. **(G)** MoTrPAC DataHub targeted bulk RNA-Seq results confirms transcriptional downregulation of multiple collagen isoforms (*COL3A1, COL6A1, COL6A2, COL6A3*) in male (but not female) gastrocnemius muscle of endurance trained rats. **(H)** Cytokine array shows clear trends towards sex-specific increases in VEGF levels in male (but not female) endurance-trained and TFEB-overexpressing quadriceps muscle lysates. One-way ANOVA within sex with post-hoc test. **(I)** Transcriptional regulation of skeletal muscle VEGF levels also appears to be sex-specific (males only) in endurance-trained rats (MotrPAC DataHub). **(J)** Quantification of neuromuscular junction (NMJ, α-Bungarotoxin/nicotinic acetylcholine receptors in green, synaptophysin in magenta, laminin in red) shape in longitudinal cryosections reveals significant increases in complex NMJs in female (but not male) runner TA muscle. One-way ANOVA within NMJ type/sex with post-hoc test. Scale bars = 10 µm. All MoTrPAC results are reported as logFC over sex-matched sedentary controls. * Statistical difference between sedentary and runner mouse samples, # Statistical difference between runner and cTFEB;HSACre mouse samples, one-way ANOVA with post-hoc test. % Statistical difference of endurance training as reported by MoTrPAC. $ Statistical difference between trained and endurance groups within each sex as reported by MoTrPAC. * P ≤ 0.05, ** P ≤ 0.01. Lack of annotation indicates comparisons were not significant. Data is represented as mean ± SEM unless otherwise noted.

In females, runner and cTFEB;HSACre IIb myonuclei both showed upregulation of extracellular matrix (ECM)-related terms, including *ECM structural constituents, ECM binding, collagen-containing ECM, external encapsulating structure,* and *heparin binding* (**Figure 5D**). These same gene sets were markedly downregulated in males from both intervention groups, underscoring a robust sex-dependent divergence in ECM remodeling under both conditions (**Figure 5D**). Importantly, Reactome analysis of the same DEGs revealed similar sex-dimorphic patterns, with opposite transcriptional regulation of ECM-associated categories, including *non-integrin* and *integrin ECM interactions*, *ECM matrix organization*, *ECM degradation,* and *collagen degradation*, in males vs. females across both groups (**Supplementary Figure S5C-D**). Remodeling of the extracellular matrix (ECM) is essential for skeletal muscle regeneration, adaptation, and repair ^47,48^, particularly through collagen turnover. This process is facilitated by matrix metalloproteinases (MMPs), which directly degrade ECM components ^49,50^, and tissue inhibitors of metalloproteinases (TIMPs), which are the primary inhibitors of MMP activity ^51^. Consistent with the sex-dimorphic transcriptional responses observed in our snRNA-seq (**Figure 5D**), re-analysis of our previously published cytokine array analysis of endurance-trained and TFEB-overexpressing quadriceps muscle ^21^ reveals trends toward sex-specific shifts in MMP2, proMMP9 (**Figure 5E**), and TIMP1 (**Supplementary Figure S5E**) abundance, with males trending towards increases and females trending towards decreases relative to their sedentary cognate groups. To examine whether these changes in MMP abundance translate into functional ECM remodeling, we examined staining patterns of collagen 6, a major component of the skeletal muscle ECM, via immunofluorescence. Consistent with our snRNA-Seq results, we detected increased levels of collagen 6 staining in female IIb^+^ fibers in cTFEB;HSACre muscle, with trends towards increased stains in female runner muscle as well (**Figure 5F**, n=3-4/group, independent cohort), which was not observed in IIb-fibers of the same groups. No changes in Collagen 6 staining were detected in male fibers of any group (**Figure 5F**). Targeted interrogation of MoTrPAC bulk RNASeq datasets reveals significant training effects in male (but not female) skeletal muscle from endurance-trained rats for transcriptional downregulation of multiple collagen isoforms (*COL3A1*, *COL6A1*, *COL6A2*, and *COL6A3*) (**Figure 5G**), similar to what we observed in our snRNA-seq of mouse IIb myonuclei (**Figure 5B**, MoTrPAC Datahub).

MMP-mediated cleavage of ECM components contributes to the release of sequestered growth factor VEGFA, ultimately contributing to VEGF-associated angiogenesis in skeletal muscle ^52,53^. Using the same cytokine array mentioned above ^21^, we also detected trending increases in VegfA abundance in both male runner and cTFEB:HSACre muscle lysates, with little change in female cognate muscle lysates (**Figure 5H**), indicating its potential release from ECM-bound stores ^54,55^. Consistent with these findings, RNA-Seq analysis confirms an early (week 1 of training) and robust *VEGF* transcriptional response in male (but not female) endurance-trained rat muscle (**Figure 5I**, MoTrPAC Datahub), a response that eventually is transcriptionally silenced as endurance training progresses (week 8). Altogether, this data suggests that endurance training and TFEB-overexpression promote ECM remodeling in IIb mouse myonuclei, potentially engaging opposing transcriptional profiles in either sex and leading to sex-specific alterations in free VEGF levels.

Additional GO Biological Processes and Molecular Function gene sets that were upregulated in female runner and TFEB-overexpressing IIb myonuclei included *axon guidance, Ras protein signal transduction, neuron projection regeneration, neuron cell–cell adhesion*, and *plasma membrane adhesion molecules* (**Figure 5D**). Notably, these neuronal signaling terms were downregulated in male runners but upregulated in male cTFEB;HSACre IIb myonuclei (**Figure 5D**), supporting the idea that TFEB overexpression in males induces an intermediate, female exercise–like transcriptional state. These GO signatures also suggest neuromuscular remodeling may be a strong feature of transcriptional responses in IIb myonuclei. Indeed, endurance training increases neuromuscular junction (NMJ) size and structural complexity and generally improves transmission reliability across this specialized motor neuron-muscle synapse ^56^. Consistent with our transcriptional results, we observed an enrichment in complex ‘pretzel-like’ neuromuscular junction structures in TA muscle from female (but not male) runners, with some similar trends in cTFEB;HSACre muscle of both sexes (**Figure 5J**). MoTrPAC confirmed dynamic and sex-dimorphic responses to endurance training by multiple integrin isoforms (ie. *ITGA4*, *ITGA6*, *ITGA7*) involved in NMJ architecture, specifically in male rat muscle (**Supplementary Figure S5F-H,** MoTrPAC Datahub). Overall, this data suggests that endurance training may have preferential effects on growth factor responses (males) and neuromuscular junction remodeling (females), likely mediated through sex-dimorphic responses to extracellular matrix dynamics and processing.

#### IIx Myonuclei

We next analyzed transcriptional responses in IIx myonuclei, which encode for an intermediate (glycolytic/oxidative) fiber with intermediate fatigue resistance and contraction speed (**Figure 6A**). As before, Type IIx myonuclei strongly clustered by intervention in our MDS analysis, with TFEB-expressing samples clustering between sedentary and runner groups for both sexes (**Figure 6B**). We found 1680 GO gene sets significantly differently expressed in female runners, 863 in female TFEBs, of which 300 (∼13%) overlapping between conditions. We found 1308 differentially expressed GO gene set categories in male runners, 1531 in male TFEBs, with 521 (∼22%) overlapping (**Figure 6C**, and see **Supplementary Table I**). The top 5 up- and down-regulated gene sets for each group are shown in **Supplementary Figure S6A**. Overall, the top ten strongest differentially expressed categories mapped to mitochondrial-associated terms (**Figure 6D**). Notably, in females, runner and cTFEB;HSACre IIx myonuclei exhibited near-complete overlap in upregulated gene sets (10/10 top GO categories), with strong enrichment for mitochondrial functions (**Figure 6D**). These included GO Cell Compartment gene sets of the *respirasome, NADH dehydrogenase complex, inner mitochondrial membrane, cytochrome complex,* and GO Biological Process for *oxidative phosphorylation* (**Figure 6D**). Male runner muscle displayed a similar mitochondrial transcriptional signature (10/10 top GO categories), indicating a conserved exercise-induced mitochondrial program activated by running across sexes. In contrast, IIx myonuclei from male cTFEB;HSACre muscle showed a marked downregulation of these same top 10 GO mitochondrial gene sets, representing a striking sex- and TFEB-specific divergence (**Figure 6D**). Reactome analysis further confirmed this pattern: while female runner and cTFEB;HSACre samples and male runners showed consistent enrichment in Reactome gene sets such as *respiratory electron transport* and *complex I biogenesis*, these same gene sets were transcriptionally suppressed in male TFEB-overexpressing muscle (**Supplementary Figure S6C-D**). We have previously reported that TFEB overexpression promotes maintenance of mitochondrial function and bioenergetic reserves in skeletal muscle, preventing the age-associated decline of mitochondrial respiration ^19^. This suggests that this transcriptional down-regulation of mitochondrial-associated gene sets in male TFEB type IIx myonuclei may be uncoupled from true mitochondrial function and/or may be de-repressed during the aging process.

**Figure 6:**
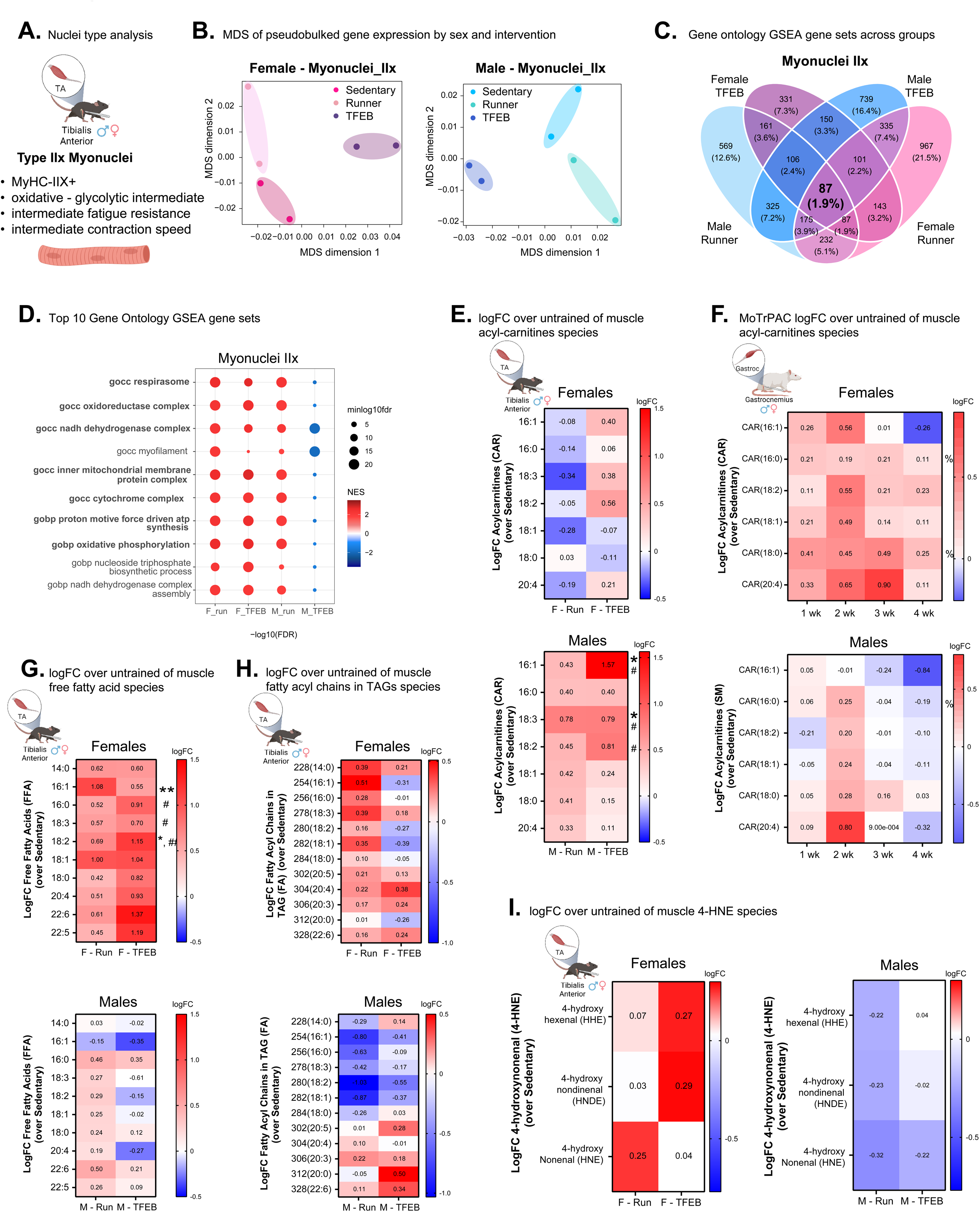
Mitochondrial and lipidomic signatures converge in IIx myonuclei with sex-specific metabolic divergence. **(A)** Type IIx myonuclei type functional characteristics. **(B)** MDS plots for myonuclei IIx in both sexes. See also **Supplementary Figure S3**. **(C)** Shared and unique GO terms (BP, CC, MF) enriched by GSEA in IIx myonuclei across all four conditions. The percentage of overlapping pathways is in parentheses. **(D)** Top ten GO terms enriched in at least one condition in IIx myonuclei, highlighting shared and divergent transcriptional signatures across sex and intervention. Note the robust enrichment for mitochondrial function pathways. **(E)** Targeted lipidomics for acyl-carnitine (CAR) lipid species showing sex-dimorphic trends (runners: decreased in females, increased in males) and intervention-dimorphic trends (TFEB: increased in females, increased in males) in the mouse tibialis anterior muscle. **(F)** Lipidomics analyses shows opposite sex-responses in acyl-carnitine abundance in young rat gastrocnemius muscle across the training period, with females trending towards higher levels and males trending towards lower levels (MoTrPAC Datahub). **(G)** Significantly increased levels of free fatty acids (FFA) lipid species in female (but not male) runner and TFEB-overexpressing TA muscle. (**H**) Sex-dimorphic effects of endurance training on fatty acyl chains in triglcyerides. **(I)** Sex-dimorphic effect of running and TFEB expression for 4-hydronynonenal (4-HNE) species in female (trending increase) and male (trending decrease) TA muscle of both groups. All MoTrPAC results and lipid levels reported as logFC over sex-matched sedentary controls. * Statistical difference between sedentary and runner samples, two-way ANOVA with post-hoc test. # Statistical difference between sedentary and cTFEB;HSACre samples, two-way ANOVA with post-hoc test. % statistical difference of endurance training as reported by MoTrPAC. Lack of annotation indicates comparisons were not significant.

Mitochondria serve as the principal sites of fatty acid β-oxidation, utilizing acylcarnitines, circulating free fatty acids (FFAs), and triacylglycerols (TAGs) as key fuels. This oxidative capacity is augmented by aerobic exercise training, which promotes enhanced utilization of lipid stores and dynamic triacylglycerol remodeling, although some of these changes have recently been found to be sex-dependent ^57^. In agreement with this, targeted lipidomics analysis revealed minor or no changes in total levels of free acylcarnitine in female skeletal muscle under either condition, while both endurance training and TFEB-overexpression trended towards increasing total levels of free acylcarnitine in male TA muscle (**Supplementary Figure S6E**). Examination of individual long-chained acylcarnitine species confirmed that while female runner muscle had no significant changes in carnitine abundance and even trended towards reductions in most species detected (**Figure 6E**), male runner muscle had significant increases in carnitine 16:1 and 18:2, with all other detected species also trending in the same direction (**Figure 6E**). Male cTFEB;HSACre muscle strongly followed the same trends, with significant increases in 18:3 and 18:2 saturated carnitines (**Figure 6E**). Interestingly, MoTrPAC gastrocnemius muscle lipidomics of the same long-chained acylcarnitines profile suggested an opposite sex effect, with female endurance-trained rat muscle increasing acylcarnitine species to a higher level than males (**Figure 6F**). Individual long-chain free fatty acid (FFAs) species (another oxidation source associated with endurance training) were increased in female runner and cTFEB; HSACre transgenic muscle (**Figure 6G**), with trending elevations of total free fatty acid levels (**Supplementary Figure S6F**). By contrast, free fatty acids remained mostly unchanged in male samples (**Figure 6G** and **Supplementary Figure S6F**). We also detected a female-specific increase in fatty acid chains esterified within triacylglycerols (FAs), which was partially recapitulated by TFEB overexpression (**Figure 6H** and **Supplementary Figure S6G**). Conversely, both male runner and male TFEB-muscle had a selective decrease in small (14:0 – 18:0) esterified fatty acid chains within triacylglycerols. Furthermore, male TFEB muscle increased the abundance of long (20:5 – 22:6) esterified fatty acid chains within triacylglycerols (**Figure 6H** and **Supplementary Figure S6G**), once again mirroring the female responses to exercise. Finally, analysis of 4-hydroxynonenal (4-HNE), a by-product of β-oxidation–driven oxidative stress, showed little change in total levels in female runners and even a mild increase in female cTFEB;HSACre muscle (**Figure 6I and Supplementary Figure S6H**), with opposite mild decreases in 4-HNE species in male runner and TFEB-overexpressing muscle (**Figure 6I and Supplementary Figure S6H**). These changes appeared to be driven mostly by 4-hydroxynonenal (HNE) in runners, and by 4-hydroxyhexenal (HHE) and 4-hydroxynondinenal (HNDE) species in TFEB-muscle instead, potentially suggesting differential fatty acid processing capabilities in runners vs. TFEB-overexpressing muscle. Overall, this data indicates potential sex-specific effects in mitochondrial β-oxidation (the breakdown of fatty acids in the mitochondrial matrix) and lipolysis (either intramuscular or in adipose tissue) activated by endurance exercise. Furthermore, it suggests that TFEB-overexpression promotes mitochondrial remodeling in both male and female skeletal muscle IIx myonuclei in patterns reminiscent of those seen in female runner muscle, with potential sex-discrepancies in the transcriptional regulation and functional outcomes of this process.

#### Endothelial Cell Nuclei

Endothelial cells play a critical role in regulating vascular tone, angiogenesis, and nutrient exchange in skeletal muscle (**Figure 7A**). In the context of exercise, they contribute to skeletal muscle adaptations by promoting capillary remodeling ^58^ and enhancing perfusion ^59^. Consistent with patterns observed in myonuclei (**Figures 4-6**), snRNA-seq analysis of endothelial cell nuclei revealed that they also undergo sex- and condition-specific transcriptional remodeling in response to VWR and muscle-TFEB overexpression, with some overlap between conditions (**Figure 7B-D** and **Supplementary Figure S7A**). We found 419 significantly differentially expressed GO gene sets in female runners, and 634 GO significantly differentially expressed categories in male runners, of which only 59 (∼6%) overlapped (**Figure 7C** and **Supplementary Figure S7B,** and see **Supplementary Table I**). Within-sex comparisons revealed 701 significantly differentially expressed GO categories in female TFEBs, of which 122 (∼12%) overlapped with runners of the same sex (**Figure 7C** and **Supplementary Figure S7B**). We found 1080 significantly differentially expressed GO categories in male TFEBs, of which 184 (12%) overlapped with male runners. Of these, only 14 GO categories were shared across all 4 conditions (**Figure 7C-D**), which we discuss below.

**Figure 7:**
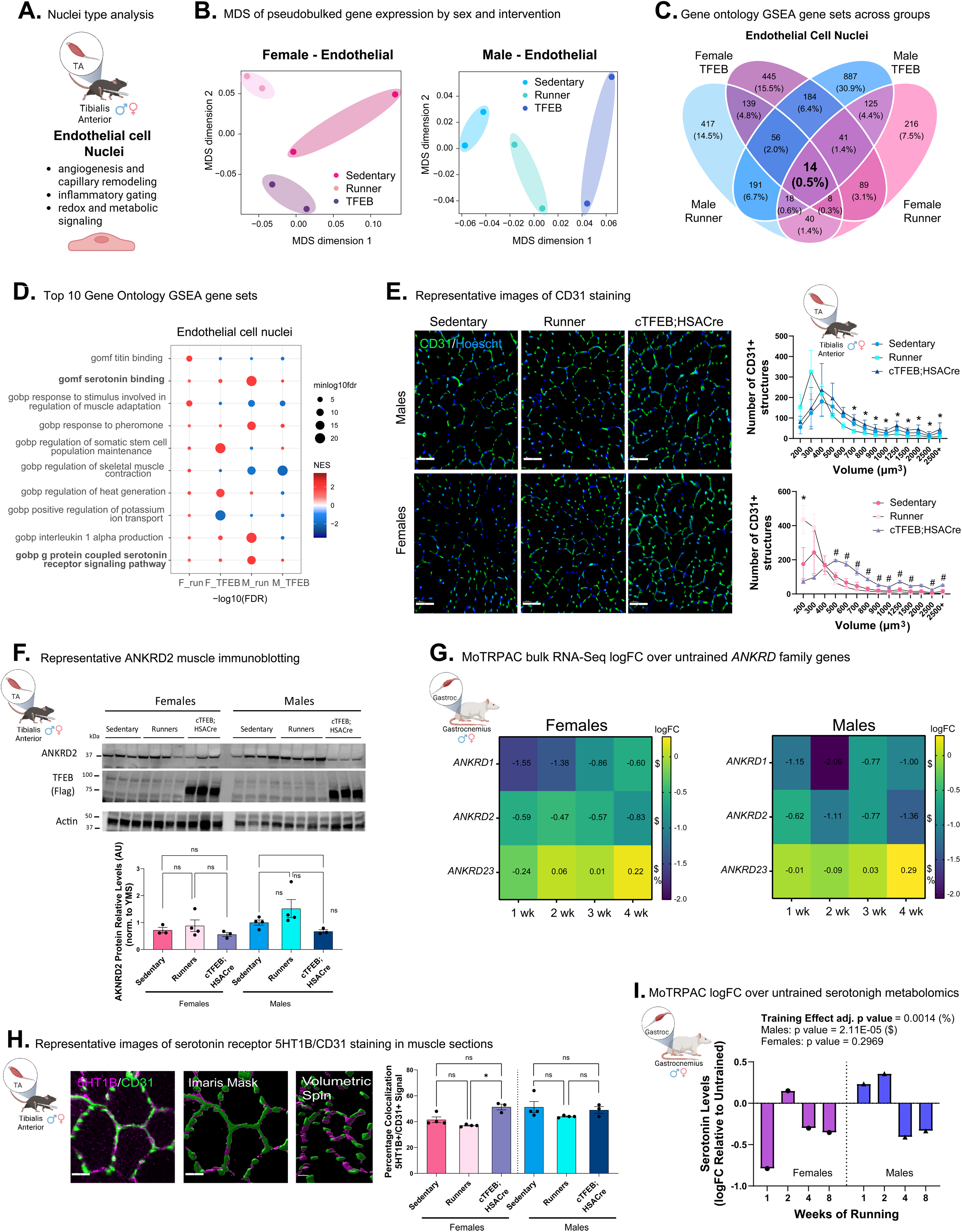
Sex-specific endothelial mechanosensory and serotonin pathway activation in TFEB/exercise responses. **(A)** Endothelial cell (EC) nuclei functional characteristics. **(B)** MDS plots for EC nuclei in both sexes. See also **Supplementary Figure S3**. **(C)** Shared and unique GO terms enriched by GSEA in EC nuclei across all four conditions. The percentage of overlapping pathways is in parentheses. **(D)** Top ten GO terms enriched in at least one condition in EC nuclei, highlighting shared and divergent transcriptional signatures across sex and intervention. Setononin pathway is highlighted in bold. **(E)** Immunostaining and distribution curves of CD31+ structures sizes within each group by sex show increased angiogenesis in both runner and TFEB-overexpressing TA muscle (CD31 in green). One-way ANOVA within sex with Kruskal-Wallis post-hoc test. * Statistical difference between sedentary and runner. # statistical difference between sedentary and TFEB. Scale bar = 60 µm **(F)** Increased levels of mechanosensor Ankyrin Repeat Domain 2 (ANKRD2/ARPP) protein in male runner muscle lysates. Marker densitometry quantification relative to actin (loading control) below. Data is represented as mean ± SEM, with one-way ANOVA within sex with post-hoc test. **(G)** MoTrPAC DataHub targeted bulk RNA-Seq analysis confirms downregulation of multiple mechanosensory family members in gastrocnemius muscle of endurance-trained rats of both sexes, including *ANKRD1*, *ANKRD2*, *AKNDR23*. **(H)** Serotonin receptor 5HT1BR colocalizes with endothelial cell marker CD31 and this colocalization is increased in female (but not male) TFEB expressing muscle. (CD31 in green, 5HT1B in magenta). One-way ANOVA within sex with Kruskal-Wallis post-hoc test. Scale bar = 20 µm. **(I)** MoTrPAC metabolomics confirmed endurance training effect on serotonin levels in male (but not female) gastrocnemius muscle. For immunostainings: each data point represents the average of three separate images collected from three sections/individual. MoTrPAC results reported as logFC over sex-matched sedentary controls. Statistical tests as listed. N.s: non-significant, * P ≤ 0.05, ** P ≤ 0.01. % Statistical difference of endurance training as reported by MoTrPAC. $ Statistical difference between trained and endurance groups within each sex as reported by MoTrPAC. Lack of annotation indicates comparisons were not significant. Data is represented as mean ± SEM unless otherwise noted.

First, and as predicted by our snRNA-seq nuclei proportion analysis (**Figure 2D**), we confirmed a significant increase in the number of endothelial cell marker CD31^+^-small structures (sizes 200-300 µm³) in TA muscle of runners of both sexes (**Figure 7E**). Interestingly, TFEB-overexpression also increased the total number of CD31+ structures in male (and trending in female) (**Supplementary Figure S7E**), but this effect was driven by increases in the number of medium-to-large size range (>500 µm³) CD31^+^-structures instead (**Figure 7E**). This increase was especially pronounced in the TA muscle of female cTFEB;HSACre mice (**Figure 7E**). These findings suggest that TFEB overexpression promotes endothelial cell activation and potentially enhances angiogenesis in skeletal muscle, similar to endurance training.

Functional enrichment analysis by GSEA of Gene Ontology terms identified previously unreported female runner-specific enrichment of terms related to *titin binding* (Molecular Function)*, regulation of skeletal muscle contraction*, and *responses to stimuli involved in muscle adaptation* (Biological Process), gene sets which were conversely downregulated in male runners and in cTFEB;HSACre endothelial cell nuclei from both sexes (**Figure 7D**). Ankyrin Repeat Domain-Containing Protein 2 (ANKRD2) is a titin-binding protein primarily expressed in skeletal muscle, where it acts as a mechanosensor responding to stretch and muscle stress during exercise ^60^, but also as a potential regulator of exercise-activated inflammatory signals ^61^. Consistent with our transcriptional analysis, immunoblotting confirmed trends towards reduced abundance of ANKRD2 protein in TFEB-overexpressing muscle lysates, and an opposite trend towards increased levels in runner-lysates of both sexes (**Figure 7F**). Furthermore, we confirmed multiple members of the muscle ankyrin repeat protein family (ie. *ANKRD1*, *ANKRD2*, *ANKRD23*) are also transcriptionally responsive to endurance training in young trained rats (**Figure 7G**, MoTrPAC Datahub), and that they also display sex-specific and timing-specific responses. This may suggest that female runner muscle preferentially promotes endothelial cell activation to support skeletal muscle myofiber contraction during the exercise bouts.

Notably, a previously unreported transcriptional node that was consistently upregulated across all four experimental groups (with particularly strong effects in male runners) relative to sedentary controls was GO Molecular Function for *serotonin binding* as well as GO Biological Process for *G protein-coupled serotonin receptor signaling* (**Figure 7D**). GSEA Reactome analysis also identified *serotonin receptors* as a shared upregulated signaling gene set across all four groups (**Supplementary Figure S7C-D**). We confirmed that serotonin receptor 5HT1B is expressed in skeletal muscle (**Figure 7H** and **Supplementary Figure S7E**) and that it strongly colocalizes with endothelial cell marker CD31. We did not observe changes in the amount of this colocalization in male samples (as predicted in our snRNA-seq results), although we did detect a significant increase in 5HT1B/CD31 colocations in female TFEB-overexpressing muscle (**Figure 7H**). However, MoTrPAC DataHub metabolomics revealed that endurance training dynamically modifies serotonin levels across the training process in the gastrocnemius muscle of male (but not female) rats (**Figure 7I**), consistent with our mouse transcriptional signatures (**Figure 7**).

## DISCUSSION

Here we report that 4 weeks of voluntary wheel running, a time sufficient to promote endurance training-associated metabolic plasticity, elicits profound, sex-biased transcriptional remodeling across myonuclear and endothelial cell nuclei within tibialis anterior mouse skeletal muscle. Furthermore, we identify that key pathways activated by VWR in myonuclei, such as lipid reprogramming, immune signaling, ECM remodeling, and mitochondrial function, are partially recapitulated by TFEB overexpression, particularly in female muscle. In male muscle, TFEB induces an intermediate transcriptional state, blending facets of exercise-induced and unique TFEB-specific programs. This underscores TFEB’s capacity to mimic aspects of exercise-like adaptations, and highlights a previously underappreciated layer of sex-dependent specificity in muscle plasticity associated with endurance exercise training. Importantly, to our knowledge, this study provides a unique resource for the field as the first comprehensive, sex-informed single-nucleus transcriptional atlas of endurance exercise adaptation in mouse skeletal muscle.

Exercise activates a coordinated network of paracrine signaling among myonuclei and non-myonuclei populations within skeletal muscle, facilitating intercellular communication that supports the complex metabolic and functional remodeling during and after exercise ^1,2,4,62^. Within this space, critical gaps persist in our understanding of sex-specific muscle adaptations to exercise ^4,11,57^. To increase our understanding of shared and unique transcriptional signatures activated by endurance training across different nuclei in skeletal muscle, we performed snRNA-seq in endurance-trained and TFEB-overexpressing skeletal muscle of both sexes. Through this approach, we identified known core responders to endurance exercise, such as mitochondrial activation and anti-inflammatory signatures, some of which were detectable across multiple cell types but exhibited significant overrepresentation or enrichment in specific nuclei clusters. Furthermore, this study identifies previously uncharacterized pathways associated with endurance training, TFEB-overexpression, or both, expanding the known regulatory landscape of exercise-associated muscle metabolic signaling.

For example, we observed a prominence of sex-dimorphic lipid signaling and phospholipid-related processes in IIa myonuclei, with opposite transcriptional regulation in female vs. male runner muscle. Endurance exercise triggers extensive lipidomic remodeling in skeletal muscle, involving coordinated changes in gene expression, substrate utilization, and intracellular lipid handling ^63–68^. These adaptations enhance fatty acid uptake ^69^, mitochondrial lipid transport ^70^, and lipid oxidation to meet energy demands as well as to promote training-associated muscle metabolic remodeling ^71–73^. Notably, women exhibit higher intramyocellular lipid content ^68,74,75^, higher fatty acid transporter expression ^68,74^, and preferential lipid utilization ^76,77^ during submaximal endurance efforts compared to men ^76,78,79^. Indeed, recent studies have found higher fatty acid flux into skeletal muscle during exercise in trained women than in trained men under the same exercise intervention ^80,81^, but the functional impact of these sex differences remains unclear. Training-induced lipid metabolic adaptations in skeletal muscle translate into an increased ability to dispose of lipids from the circulation ^82^, potentially also acting as central regulators of exercise-activated metabolic health. Our findings suggest that cTFEB:HSACre skeletal muscle shares these endurance-trained transcriptional profiles that promote broad reprogramming of lipid-related pathways in skeletal muscle, with potential sex-dimorphic effects. Although the biological functions of these sex-dimorphic responses are still unknown, one possibility is that endurance-trained female muscle preferentially diverts ceramides towards sphingomyelin synthesis rather than glycosphingolipid production, consistent with a metabolic shift favoring sphingomyelin generation and transcriptional glycosphingolipid metabolic silencing. In contrast, endurance-trained male muscle exhibits reductions in both ceramide and sphingomyelin levels, implicating a possible suppression of the sphingomyelinase ceramide-processing pathway and diversion of ceramide towards glycosphingolipid metabolism instead. TFEB overexpression in male muscle also appears to mimic a late endurance training phase characterized by lowered sphingomyelin and (potentially) ceramide levels, whereas female TFEB-overexpressing muscle shows increases consistent with an early training phase instead. Interestingly, a recent study found that very-long-chain ceramide species in skeletal muscle may be linked to the protective effects of high-intensity interval training in men with type 2 diabetes ^83^, although this hypothesis remains untested to date. On the other hand, the higher levels of free fatty acids (FFAs) and fatty acyl chains in TAGs (FAs) in female runner and TFEB-overexpressing muscle, versus the higher levels of acylcarnitines in male runner and TFEB-overexpressing muscle, may represent a novel sex-dimorphic mitochondrial fuel utilization node in response to endurance training. Collectively, this may reflect a higher capacity to handle and oxidize fatty acids during exercise, and the ability to adapt fatty acid utilization to fatty acid availability in endurance training in female muscle. Additionally, given that FAs can act as signaling molecules themselves via G-protein–coupled receptors and nuclear receptor co-activators (PPARs), the increases in FFAs we see in female muscle might also reflect downstream signaling processes contributing to lipid metabolism changes themselves.

Another newly identified transcriptional gene set was associated with glucocorticoid signaling. Chronic glucocorticoid treatment induces atrophy in fast-twitch glycolytic muscle fibers, likely through its catabolic effects on protein synthesis and turnover ^84,85^. These catabolic effects of glucocorticoids on skeletal muscle are believed to contribute to age-related muscle loss and sarcopenia. In contrast, endurance training and TFEB overexpression exert anabolic effects in aging skeletal muscle, promoting geroprotection and preservation of muscle mass ^5,19^. However, the role of glucocorticoids and GR signaling during exercise adaptations remains currently unknown. Interestingly, corticosterone can also bind directly to GR (NR3C1), suggesting a complementary relationship between these two signaling nodes to ultimately promote metabolic plasticity during exercise-mediated reprogramming. Altogether, our findings suggest that endurance training and TFEB-overexpression may antagonize glucocorticoid receptor signaling in fast-twitch fibers of mice and rats, a novel signaling node of particular relevance to age-associated muscle atrophy and disease.

Serotonin regulation was another newly discovered signaling node activated in endothelial cells by VWR and TFEB overexpression. Serotonin is a monoamine neurotransmitter, and accumulating evidence suggests that serotonin and serotonin receptors play important roles in the regulation of energy balance, glucose homeostasis, and lipid metabolism in peripheral tissues ^86^. Although studies have extensively investigated peripheral serotonin in metabolic dysfunction, its exact role in skeletal muscle remains largely underexplored. Interestingly, evidence shows that serotonin deficiency impairs skeletal muscle adaptations to endurance training ^87^ and improves glucose uptake, insulin sensitivity, and overall metabolic health under high-fat-diet-induced obesity ^88^. Furthermore, serotonin enhances muscle stem cell function and regeneration ^89^, suggesting that it may represent a new neurotransmitter modulator of exercise-associated plasticity in skeletal muscle. In agreement with this, MoTrPAC metabolomics studies found an early peak (weeks 1-2) of serotonin abundance in male (but not female) endurance-trained muscle. While the consequences of these sex-specific effects remain unknown, our data suggest that the endothelial cell microenvironment is dynamically remodeled alongside myonuclei under both VWR and TFEB-overexpression conditions, promoting sex-specific niche remodeling.

In agreement with this, muscle endothelial cells were recently identified among the strongest responders to exercise in a landmark single-cell RNA-sequencing analysis of multiple stem cell compartments ^45^, suggesting they may be previously underappreciated regulators of exercise-associated skeletal muscle plasticity. In agreement with this, we found that muscle endothelial cell nuclei are quite responsive to endurance training and myofiber TFEB-overexpression. In general, endothelial cell nuclei activated programs implicated in paracrine signaling coordination (such as serotonin receptor signaling and IL-1α production), suggesting potential mechanical communication between myonuclei and vasculature. In agreement with this, we saw increases in CD31-staining patterns in both runner and TFEB-overexpressing TA muscle, although each intervention appeared to target different CD31+-vessel sizes. We observed that while running increased capillary density (increased abundance of smaller CD31+ structures, indicative of sprouting angiogenesis, i.e. new vessel formation), TFEB appeared to induce capillary complexity instead (increased abundance of larger CD31+ structures, more indicative of intussusceptive angiogenesis or vessel splitting), particularly in female muscle. Interestingly, we also determined that running and TFEB-overexpression increases levels of VEGF in male but not female muscle. Since intussusceptive angiogenesis is less reliant on VEGF and can occur independently or with minimal VEGF input ^90^, this may reflect parallel angiogenic signaling processes activated by running and TFEB-overexpression. Alternatively, the timing of angiogenic responses and the degree of response may be correlated with running distance and/or TFEB-expression levels, which we also discuss below.

It is interesting that despite the use of a myofiber-specific HSA-Cre driver ^91^ (the specificity of which was also validated by recent single-nuclei RNA-seq studies ^27^), we detected significant transcriptional responses in non-myofiber (ie. Endothelial cell) nuclei in our cTFEB;HSACre transgenic model. This suggests that myofiber-specific TFEB overexpression induces cell non-autonomous transcriptional and metabolic remodeling across multiple muscle cell types, potentially mediated by local myokine secretion. In agreement with this, we have previously demonstrated that muscle-TFEB overexpression increases the secretion of the known muscle-originating, exercise-responsive circulating factor cathepsin B ^19^. The precise mechanisms driving the cell-non-autonomous transcriptional signatures in cTFEB;HSACre muscle we report here remain currently unknown. TFEB is a central regulator of lysosomal network dynamics, including lysosomal exocytosis ^92^, suggesting TFEB activation (either via overexpression and/or running) may promote secretion of intracellular cargo. Given the well-known remodeling effects of exercise on both muscle and organismal ‘secretome’ ^93,94^, this may indicate TFEB as a potential regulator of the exercised muscle-secretome. Indeed, the overlap between exercise and TFEB-induced transcriptional changes reported here, as well as our previously reported functional outcomes in cTFEB;HSACre transgenic mice, indicates endurance-like remodeling and geroprotective effects of muscle-TFEB overexpression ^19,21^, and suggests that TFEB may function as a pan-cell regulator capable of activating exercise-like transcriptional and metabolic programs in skeletal muscle, particularly in females. This is supported by previous work demonstrating that skeletal muscle TFEB expression is required for the full metabolic remodeling observed after endurance training ^12^. These proof-of-concept findings confirm the utility of this approach, highlighting the shared pathways mediating skeletal muscle remodeling across both interventions and positioning TFEB as a promising node for modulating muscle aging and disease. The identification of TFEB-specific single-nuclei transcriptional signatures absent from exercised muscle and with known geroprotective effects during aging further highlights its potential to both mimic and extend the benefits of physical activity.

The sex-specific expression patterns we observed across multiple cell types and interventions are particularly striking. Of note, transcriptional programs involved in extracellular matrix remodeling showed some of the most robust sex-dimorphic responses, with marked upregulation in female and downregulation in male type IIb myonuclei. Existing literature suggests that multiple ECM-related gene sets show widespread female-bias in expression in primary mouse neutrophils ^95^, as well as robust sex differences in ECM remodeling in valvular interstitial cells ^96^ and smooth muscle progenitor cells ^97^. Similar to what we report here, previous work has shown that resistance exercise induces MMP2, MMP9, and TIMP1 expression and activity in human muscle ^98^, although sex was not included as a biological variable in that study. Our findings of elevated MMP2 and proMMP9 in males (but not females) after running and TFEB-overexpression may reflect enhanced ECM degradation capacity (given their roles in collagen degradation), which aligns with TFEB-mediated lysosomal-autophagy upregulation, potentially promoting matrix turnover. The lack of change in these factors in females suggests restrained or parallel ECM remodeling despite increased running burden, potentially preserving ECM integrity amid training adaptation and/or reflecting earlier dynamic responses not captured due to our cross-sectional approach.

Although substantial strides have been made in the past few years, the field of exercise biology remains predominantly male-centric ^4,57^, limiting our ability to contextualize and interpret many of the sex-dependent molecular responses found in this study. Building on existing paradigms in the field of sex-differences research, several non-mutually exclusive factors may underlie the sex-specific, sex-dimorphic, or sex-agnostic differences we report here. First, there were substantial differences in total voluntary wheel running distance for our two runner groups, with female mice running approximately four times more than their male counterparts. This is a well-known and extensively reported sex-dependent variable in mice ^99–101^, and although the mechanisms underlying sex-divergent voluntary running performance remain contested, differences in ovarian hormones, energy metabolism, and potential evolutionary/behavioral strategies have been proposed ^102–105^. This disparity raises the possibility that some of the observed transcriptional profiles simply reflect differences in voluntary exercise burden and its associated metabolic and mechanical demands on skeletal muscle. However, we also report here roughly threefold higher TFEB expression in male compared to female cTFEB;HSACre transgenic muscle, which we have previously confirmed to also exist at the protein level ^19^. Thus, the strong concordance between key transcriptional profiles and functional outcomes in runner and cTFEB;HSACre muscle within each sex, despite significant and opposing differences in both running performance and TFEB expression, suggests the existence of convergent, sex-dependent transcriptional programs activated by both endurance exercise and TFEB overexpression. Second, it is important to note that while female cTFEB;HSACre muscle consistently mimicked exercise-associated upregulation of mitochondrial, ECM, and immune-related programs, for example, male TFEB-overexpressing muscle often diverged, either showing transcriptional repression (as in IIx mitochondrial programs) or an intermediate activation state (as seen in IIa and IIb myonuclei). These data suggest that sex-specific differences in TFEB activity, Cre driver expression, and/or epigenetic landscape may further influence the transcriptional response to TFEB overexpression. To date, our understanding of how these biological mechanisms may differ by sex remains woefully incomplete. However, the rapidly evolving field of sex differences has highlighted multiple physiological responses that diverge between males and females, including differential susceptibility and adaptation to aging ^106^, disease progression, pharmacological interventions ^107^, and environmental/lifestyle stimuli such as exercise ^4,10^. Our work adds to this critical gap in knowledge, showcasing differential exercise responsiveness in male and female muscle nuclei and raising important considerations for personalized exercise medicine strategies. Additionally, temporal evaluation of exercise burdens (ie. examining acute versus chronic exercise, as done in MoTrPAC ^4^) and/or TFEB induction (with inducible Cre-systems, for example), could illuminate dynamic transcriptional trajectories and transient compensatory mechanisms. Finally, exploring hormonal regulation and/or chromosomal contributions will be needed to shed light on the mechanistic drivers of the sex-dimorphic profiles observed here. Importantly, the consistent overlap between female endurance-trained and TFEB-induced gene programs supports the utility of TFEB as a discovery platform for conserved exercise-responsive programs in a severely underserved population, while also underscoring the necessity of incorporating sex as a critical variable in exercise mimetic development.

## METHODS

### Animals

We have previously described the generation of fxSTOP-TFEB transgenic and cTFEB;HSACre double transgenic mice. For this work, we utilized cTFEB;HSACre double transgenic male and female littermates and young wild-type mice of both sexes from the National Institute of Aging (NIA) colony (∼4 months of age). NIA mice were acclimated to our vivarium for one month. At 5 months of age, all mice enrolled in these experiments were single-housed with access to in-cage locked (sedentary) or free (running) wheels for 4 weeks (see below). Animals were allocated at random for snRNA-seq studies (N=4/group), immunofluorescence and western blotting (n=3-4/group, left/right TA each), or lipidomics analysis (n=4/group).

Body weight was measured before euthanasia and tissue collection. All animals are in the C57BL/6J genetic background and maintained in standard shoebox-style cages with filter tops with a 12:12 light-dark cycle. Mice were not fasted before tissue collection, and they were allowed free access to standard NIH-07 mouse chow throughout the experiment. All animal experimentation adhered to NIH guidelines and was performed by blinded investigators (when possible) and was approved by and performed in accordance with the University of Southern California (USC) Institutional Animal Care and Use.

### Exercise Protocol

To equilibrate the influence of single housing on animal behavior, each mouse from sedentary/runner cohorts, as well as an age-matched additional cTFEB;HSACre individuals, was single-caged in standard vivarium polypropylene cages (290 × 180 × 160 mm) throughout the experiment. Bedding for all cages was changed to sawdust and included two nestlets and manzanita sticks for enrichment. Each running mouse had free access to a running wheel (10.16 cm in diameter) connected to a counter to record the running distance (Columbus Instruments monitoring software). This in-cage wheel running system for mice collects the number of revolutions and the timing of the revolutions for each cage every 1 minute. Sedentary controls were also given free access to a locked wheel to control for enrichment. Investigators monitored animal welfare and physical activity daily.

### Tissue collection

Tissue collections were all undertaken between 9-11AM on the last day of the fourth-week of the trial. Animals were anesthetized with 3.8% Avertin solution (2,2,2-tribromoethanol (Sigma, T48402) and 2-methyl-2-butanol (Sigma, 152463) in miliQ water) prior to tissue collection. Euthanasia was completed via diaphragm transection followed by transcardial perfusion with 60 mL of ice-cold 1x PBS. Tibialis anterior skeletal muscle tissue was flash-frozen in liquid nitrogen and/or frozen in OCT following dissection and stored at -80 C until nuclei extraction and downstream analyses.

### Immunofluorescence Staining

Skeletal muscle tissue was embedded in OCT (TissueTek, 4583), frozen in 2-methylbutane (VWR, 103525-278) vapors before submersion in liquid nitrogen and stored at -80 C until used. All samples were sectioned on a standing cryostat (Leica) and sections were 20 mm thick.

### Immunofluorescence staining and Quantification

Muscle sections from a second independent cohort of mice were permeabilized with 0.1% Triton (Thermo Scientific, TritonX-100, A16046AE) in 1X PBS for 15min and blocked with 5% BSA for 30min. Sections were incubated in primary antibodies: anti-laminin (R&D Systems, MAB2549, 1:100), anti-5HT1B (Millipore Sigma, SAB4501470, 1:200), anti-CD31 (BD Pharmingen, 550274, 1:200), anti-COL6A1 (Proteintech, 17023-1-AP, 1:200), anti-synaptophysin (Cell Signaling Technology, D8F6H, 1:200), and anti-MHC Type 2B (DSHB, BF-F3, 1:100) overnight at 4°C. The next day, slides were washed with 1X PBS 3 times, 5min each and incubated in secondary antibodies for 1hr at RT. All antibodies were diluted in 5% BSA. Next, slides were incubated in Hoechst (ThermoFisher Scientific, 62249, 1:5000) for 10min, and fixed with 4% PFA (Electron Microscopy Sciences, 32% PFA, 15714-S) in 1X PBS for 5min. For Alpha-Bungarotoxin, CF(R)640R (Biotium, 00004, 1:100), slides were incubated for 15min following PFA. Slides were washed 2-3 times in between steps with 1X PBS for 5min each. Finally, slides were dried and mounted with Fluoromount-G Anti Fade (Southern Biotech, 0100-35). Muscle sections were imaged with a Stellaris confocal microscope (Leica) at 20x or 63x magnification, and Z-stacks of 10 microns were taken. When appropriate, sections were imaged with an ECHO Revolution epifluorescent microscope at 10x magnification, and tile scans of entire sections were acquired with no Z-stack, yielding 2D images which were used for analysis. Immunofluorescence images were analyzed using Imaris software (v. 10.2). For quantitation of signal, the Surface function was employed to generate 2D/3D reconstructions based on signal intensity. Segmentation of muscle fibers was performed when appropriate for normalization purposes, and desired metrics were exported as Excel files for further analysis. To assess neuromuscular junction morphology, k-means clustering was performed following Imaris 3D reconstruction. Complex, Linear, and Fragmented categories were determined based on the volume, elongation, and number of components of the 3D reconstructions. An Excel VBA macro was created and used to standardize features and run k-means clustering (k = 3) with 50 random restarts which run until convergence and chooses the best clustering with the lowest total within-cluster sum of squared distances.

### Fiber typing Staining and Analysis

For fiber typing, muscle sections were permeabilized with 0.1% Triton (Thermo Scientific, TritonX-100, A16046AE) in 1X PBS for 15min and blocked with 5% BSA for 30min-1hr. Sections were incubated in anti-laminin (Abcam, ab11575, 1:200), anti-MHC Type 1 (DSHB, BD-D5, 1:100), anti-MHC Type 2A (DSHB, SC-71, 1:100), and anti-MHC Type 2B (DSHB, BF-F3, 1:100) overnight at RT. The next day, slides were washed with 1X PBS 3 times, 5min each and incubated in secondary antibodies for 1hr at RT. All antibodies were diluted in 5% BSA. Next, slides were washed with 1X PBS 3 times, 5min each and fixed with 4% PFA (Electron Microscopy Sciences, 32% PFA, 15714-S) in 1X PBS for 5min, then washed in 1X PBS again for 3 times, 5min each. Finally, slides were dried and mounted with Fluoromount-G Anti Fade (Southern Biotech, 0100-35). Muscle sections were imaged with an ECHO Revolution epifluorescent microscope at 4x magnification. Tile scans of entire sections were acquired with no Z-stack, yielding 2D images which were used for analysis. Immunofluorescence images were analyzed using Imaris software (v. 10.2), and initial semiautomated segmentation of fibers was performed according to Gilda et al ^108^. Cell function was utilized to identify muscle fibers. Segmentation was based on cell membrane detection of the laminin signal, and the following parameters were specified: Cell Smallest Diameter - 32.0, Membrane Detail - 0.7, Filter Type - Local Contrast. The Cell Membrane Threshold was not altered. Finally, filtering of cells was performed based on cell area to remove unwanted cells. Next, segmented fibers were filtered based on Cell Mean Intensity for each immunofluorescence channel to identify individual fiber types. Following completion, Cell Area values for all segmented fibers and fiber types were exported as Excel files.

### Protein isolation and Immunoblotting

Protein lysates were prepared following previously published protocols with minor modifications ^19^. In short, flash frozen perfused tibialis anterior muscles were placed in plastic tubes with silica beads (MP Biomedical, 116913100) and homogenized in the FastPrep-24 5G bead grinder and lysis system (MP Biomedical, 116005500) in RIPA Lysis and Extraction Buffer (Invitrogen, 89900), 1X Halt Protease and Phosphatase Inhibitor Cocktail (Invitrogen, 78442). Protein concentration was quantified using a Pierce BCA Protein Assay (23227). Proteins were separated on 4–15% TGXTM gels (Bio-Rad, 5671085), blotted onto 0.45 µm porePVDF, and blocked in 5% non-fat dry milk in TBST (1X TBS with 0.05% Tween 20) for 1 h at RT. Membranes were incubated with anti-FLAG antibody (Sigma, M2, F1804, 1:1000), anti-ANKRD2 antibody (Protein Tech, 11821-AP 1:1200), anti-FKPB5 antibody (Stressmarq, SMC-138 1:1000), in PBS-T with 5% BSA at 4C overnight. Primary antibodies were visualized with horseradish peroxidase conjugated anti-rabbit (Santa Cruz, sc-2004, 1:5,000) or anti-mouse (Santa Cruz, sc-2005, 1:5,000) and enhanced chemiluminescence, or anti-mouse IgG 680 (Invitrogen, A21058, 1:10,000), anti-rabbit IgG 488 (Invitrogen, A32790TR, 1:10,000) or anti-goat IgG (Invitrogen, A21432, 1:10,000) and fluorescence detection. Densitometry analysis was performed using ImageJ software application.

### Cytokine Array

Cytokine arrays (Abcam, ab193659) and muscle lysate generation were originally described in Patterson et al ^21^.

### Skeletal Muscle Nuclei Isolation

Nuclei were isolated via homogenizing frozen TA muscle (n=4/condition) at 4°C using Miltenyi GentleMACs Octo dissociator (Cat # 130-096-427) using a pre-programmed Nuclei setting and Miltenyi Nuclei Extraction Buffer (Cat # 130-128-024). Lysate was purified using Miltenyi Anti-Nucleus Microbeads (Cat #130-132-997) and LS Columns (Cat # 130-042-401). Nuclei were counted using MACSQuant Analyzer 10 Flow Cytometer and diluted to 700-1,200 nuclei per uL in PBS with 2% BSA (Miltenyi Cat # 130-128-024). Samples were treated with 0.2X Riboguard RNAse Inhibitor (Biosearch Technologies Cat # RG90910K) during all steps. Samples from 2 independent animals were combined for each replicate to increase biological sampling.

### Single nuclei RNA-seq library construction and sequencing

Single-nucleus libraries were generated using the Chromium Next GEM Single Cell 3ʹ GEM, Library & Gel Bead Kit v3.1 (10X Genomics, PN-1000121) following the manufacturer’s guidelines. Based on flow cytometry-based nuclei counts, suspensions were loaded to target a recovery of approximately 6,000 nuclei per sample. Libraries were prepared using the Chromium Next GEM Chip G (10X Genomics, 2000177) according to the manufacturer’s instructions.

Library quality was evaluated using the 4200 TapeStation system (Agilent Technologies, G2991A) with High Sensitivity D1000 DNA ScreenTape (Agilent Technologies, 50675584). Sequencing was performed on an Illumina NovaSeq X Plus platform to generate 150 bp paired-end reads at Novogene Corporation (USA). Raw sequencing data have been deposited in GitHub (https://github.com/BenayounLaboratory/Muscle_snRNAseq_CortesLab).

### Transgene expression analysis

To determine whether the human *TFEB* transgene was appropriately expressed in the expected samples only from our single cell transcriptomics, we used a simple pseudobulking strategy. Briefly, a custom transcriptome reference was built using kallisto v0.43.0 ^109^, using the mouse gencode vM36 transcript sequences and the human TFEB open reading frame transgene sequence. Read 2 from the scRNAseq libraries (which contains transcript sequences immediately upstream of the 3’ UTR) was then pseudoaligned with kallisto using options “--single -l 150 -s 50”. Normalized transcript per million (tpm) values computed by kallisto were then used to evaluate sample-wise overall expression of human TFEB. As expected, only samples from transgenic mice showed detectable expression of the transgene.

### snRNAseq data analysis

The following pipeline was run for all libraries. CellRanger 9.0.0 (10x Genomics) was used to preprocess our libraries, together with the GRCm39 mouse reference genome. FASTQ read files were aligned to the reference genome and quantified using “cellranger count”. The ambient RNA ‘soup’ was estimated and removed using DecontX from ‘celda’ v1.16.1 in R v4.3.1 ^110^ using the raw feature barcode matrix folder. Quality control was performed in R v4.3.1 using Seurat v4.3.0.1 [XXXX] by removing dead and low-quality cells by selecting for cells that contained between 250-5000 UMIs, <15% mitochondrial-mapping reads, < 10000 UMI counts and < 25% estimated background contamination according to DecontX. A list of mouse cell cycle genes was obtained from the Seurat Vignettes (https://www.dropbox.com/s/3dby3bjsaf5arrw/cell_cycle_vignette_files.zip?dl=1), derived from a mouse study of cell cycle ^111^, and cell cycle phase was predicted using the Seurat CellCycleSorting() function. Multiplets were annotated using a combination of Doubletfinder v2.0.3 ^112^ and scds v 1.16.0 ^113^ for each individual library. Only cells called as singlets by both methods were considered to be singlets with high confidence, yielding 59569 high confidence, high quality single nuclei transcriptomes.

Gene expression was then normalized on a global scale using SCTransform v0.3.5 [PMID: 31870423], regressing to the variables “nFeature_RNA”, “nCount_RNA”, “percent.mito”, and “Phase” with the parameter variable.features.n = 5000. Dimensionality analysis revealed an optimal number of PCs of 18 (the minimum of the PC rank (i) where change of % of variation is more than 0.1% and (ii) which exhibits cumulative percent greater than 90% and % variation associated with the PC as <5), which was used for all downstream analyses (including UMAP construction).

### snRNA-seq cell type annotation

We used Seurat v4.3.0.1 to perform unsupervised cell clustering with a single nearest neighbor resolutions and a resolution of 0.6, which was chosen to provide granularity pre-annotation with 25 clusters, which served as the basis for downstream annotation efforts. Two different R packages, scSorter v0.0.2 ^114^ (marker-based) and scMAGIC v0.1.0 ^115^ (reference-based) were used. Specifically, scSorter was used using markers from (i) singleCellBase ^36^ and (ii) scMayoMap ^9^. In parallel, for use with scMAGIC, we leveraged a mouse snRNA-seq muscle reference atlas ^37^, using parameters: “atlas = “MCA”, method_HVGene = ‘SciBet_R’, method_findmarker = ‘Seurat’, cluster_num_pc = 18, min_cell = 10, method1 = ‘spearman’, percent_high_exp = 0.8 “. First pass predictions of cell type identity from both scSorter and scMAGIC runs were gated back to the original Seurat object for further processing. We then annotated each of the 0.6 SNN Seurat clusters using combined information from (i) scSorter and scMAGIC automated annotations, (ii) top predicted cell cycle phase, (iii) top 5 enriched markers in the cluster according to Seurat FindAllMarkers with parameters only.pos = TRUE, min.pct = 0.25, logfc.threshold = 0.5.

### Cell type prioritization analysis with Augur

For single nuclei RNA-seq and ATAC-seq datasets, we split objects by sex and intervention to investigate their impact on each homogenous biological group. We applied the Augur v1.0.3 ^38,39^ algorithm to determine which nuclei types were most transcriptionally impacted by intervention in each sex.

### Analysis of nuclei type proportion changes in snRNA-seq libraries

Nuclei type proportions were calculated from the final Seurat object using the R package scProportionTest v0.0.0.9000^40^ in R v4.3.1, to compare interventions to their sex-matched sedentary control in a pairwise fashion, after splitting the objects for each sex. This algorithm runs a permutation test on pairs of conditions for each nuclei type and returns the relative change in cell type abundance between groups with a confidence interval for each comparison.

### Analysis of variance explained for gene expression

The variance explained by experimental covariates was computed using the scater v1.28.0 package ^116^.

### Pseudobulking and differential gene expression analysis

We used muscat v1.14.0 ^43^ to pseudobulk the single-nuclei gene expression data by sample and cell type. Only nuclei types with ≥20 nuclei across all samples were used for downstream analyses. Analyses were conducted separately for samples from females and males due to batch processing. The package sva v3.48.0 ^117^ was used on the pseudobulked objects to estimate technical noise, and limma v3.56.2 was used to regress these effects from the counts, and DESeq2 v1.40.2 ^118^ for differential gene expression analysis. For maximum sensitivity, we considered genes with False Discovery Rate [FDR] <10% as significantly impacted by biological condition. This DEseq2-based analysis was used for all downstream analyses (*i.e.* functional enrichment).

### Functional enrichment analysis for differential gene expression and chromatin accessibility

For gene set enrichment analysis, we obtained Gene Ontology [GO] and Reactome gene sets from the Molecular Signature Database (mouse v2024.1) ^119^. DEseq2 results were used as input for phenoTest v1.48.0 to run gene set enrichment analysis (GSEA). The DESeq2-derived log_2_(FoldChange) was used to rank genes. We used 5000 permutations, minimum gene set size of 10 and maximum gene set size of 2500 to compute enrichment. Terms with a FDR < 5% were considered significantly regulated.

### Shotgun Lipidomics

Lipid species were analyzed using multidimensional mass spectrometry-based shotgun lipidomics ^120^ in a separate cohorts of individuals (n=4 group, independent cohort). In brief, each sample homogenate containing 0.5 mg of protein which was determined with Pierce BCA assay was accurately transferred to a disposable glass culture test tube. A premixture of lipid internal standards (IS) was added prior to conducting lipid extraction for quantification of the targeted lipid species. Lipid extraction was performed using a modified Bligh and Dyer procedure ^120^, and each lipid extract was reconstituted in chloroform:methanol (1:1, v:v) at a volume of 400 µL/mg protein. Phosphatidylethanolamine (PE), free fatty acid (FFA) and 4-hydroxynonenal (4-HNE) were derivatized as described previously ^121–124^ before lipidomic analysis.

For shotgun lipidomics, individual lipid extract was further diluted to a final concentration of ∼500 fmol total lipids per µL. Mass spectrometric analysis was performed on a triple quadrupole mass spectrometer (TSQ Altis, Thermo Fisher Scientific, San Jose, CA) and a Q Exactive mass spectrometer (Thermo Scientific, San Jose, CA), both of which were equipped with an automated nanospray device (TriVersa NanoMate, Advion Bioscience Ltd., Ithaca, NY) as described ^125^. Identification and quantification of lipid species were performed using an automated software program ^126,127^. Data processing (e.g., ion peak selection, baseline correction, data transfer, peak intensity comparison, and quantitation) was performed as described ^127^. The results were normalized to the protein content (nmol lipid/mg protein).

### Molecular Transducer of Physical Activity Consortium Data

Data used in the preparation of this article were obtained from the Molecular Transducers of Physical Activity Consortium (MoTrPAC) ^128^ database (MoTrPAC Datahub) ^4^, which is available for public access at https://motrpac-data.org.

### Biochemistry Statistical Analysis

All immunoblotting, immunostaining, cytokine array, and biochemical data were analyzed as described in figure legends. Statistical significance was defined as p < 0.05. Data are represented as means with standard error of the mean using GraphPad Prism 10.4.2 (La Jolla, CA). When possible, data analyses were conducted in a blinded fashion. All data were prepared for analysis with standard spreadsheet software (Microsoft Excel).

## ACKNOWLEDGEMENTS

We thank members of the lab, past and present, as well as former and current colleagues and collaborators (especially our friends in the Hill lab), for their helpful contributions.

## AUTHOR CONTRIBUTIONS

K.G., K.L., T.J., A.A., I.M., J.L., C.B., M.K., H.W., X.H., B.A. B. and C.J.C. performed the experimental work and/or data analysis, and reviewed the manuscript. K.G., B.A.B., and C.J.C. wrote and edited the manuscript. C.J.C. conceptualized the experiments outlined here and obtained (or helped obtain) most associated funding.

## REFERENCES

1 Chow, L. S. et al. Exerkines in health, resilience and disease. Nat Rev Endocrinol 18, 273–289, doi:10.1038/s41574-022-00641-2 (2022). PMC9554896

2 Gleeson, M. et al. The anti-inflammatory effects of exercise: mechanisms and implications for the prevention and treatment of disease. Nature reviews. Immunology 11, 607–615, doi:10.1038/nri3041 (2011)

3 Sousa, N. S. et al. The immune landscape of murine skeletal muscle regeneration and aging. Cell Rep 43, 114975, doi:10.1016/j.celrep.2024.114975 (2024)

4 MoTr, P. A. C. S. G., Lead, A. & MoTr, P. A. C. S. G. Temporal dynamics of the multi-omic response to endurance exercise training. Nature 629, 174–183, doi:10.1038/s41586-023-06877-w (2024). PMC11062907

5 Cartee, G. D., Hepple, R. T., Bamman, M. M. & Zierath, J. R. Exercise Promotes Healthy Aging of Skeletal Muscle. Cell Metab 23, 1034–1047, doi:10.1016/j.cmet.2016.05.007 (2016).PMC5045036

6 Lovric, A. et al. Single-cell sequencing deconvolutes cellular responses to exercise in human skeletal muscle. Commun Biol 5, 1121, doi:10.1038/s42003-022-04088-z (2022).PMC9588010

7 de Jong, J. et al. Sex differences in skeletal muscle-aging trajectory: same processes, but with a different ranking. Geroscience 45, 2367–2386, doi:10.1007/s11357-023-00750-4 (2023).PMC10651666

8 Walter, L. D. et al. Transcriptomic analysis of skeletal muscle regeneration across mouse lifespan identifies altered stem cell states. Nat Aging 4, 1862–1881, doi:10.1038/s43587-024-00756-3 (2024).PMC11645289 Biotechnology and Aegeria Soft Tissue and is an advisor for Tessera Therapeutics, HapInScience, Regenity and Font Bio. The other authors declare no competing interests.

9 Yang, L. et al. Single-cell Mayo Map (scMayoMap): an easy-to-use tool for cell type annotation in single-cell RNA-sequencing data analysis. BMC Biol 21, 223, doi:10.1186/s12915-023-01728-6 (2023).PMC10588107

10 Cortes, C. J. & De Miguel, Z. Precision Exercise Medicine: Sex Specific Differences in Immune and CNS Responses to Physical Activity. Brain Plasticity, pp. 1–13, doi:10.3233/BPL-220139 (2022)

11 Hanks, S. C. et al. Extensive differential gene expression and regulation by sex in human skeletal muscle. Cell Genom 5, 100915, doi:10.1016/j.xgen.2025.100915 (2025).PMC12366654

12 Mansueto, G. et al. Transcription Factor EB Controls Metabolic Flexibility during Exercise. Cell Metab 25, 182–196, doi:10.1016/j.cmet.2016.11.003 (2017).PMC5241227

13 Sardiello, M. et al. A gene network regulating lysosomal biogenesis and function. Science 325, 473–477, doi:10.1126/science.1174447 (2009)

14 Settembre, C. et al. TFEB links autophagy to lysosomal biogenesis. Science 332, 1429–1433, doi:10.1126/science.1204592 (2011). 3638014

15 Settembre, C. et al. TFEB controls cellular lipid metabolism through a starvation-induced autoregulatory loop. Nature cell biology 15, 647–658, doi:10.1038/ncb2718 (2013).3699877

16 Settembre, C. & Ballabio, A. Lysosome: regulator of lipid degradation pathways. Trends Cell Biol 24, 743–750, doi:10.1016/j.tcb.2014.06.006 (2014).PMC4247383

17 Evans, T. D. et al. TFEB drives PGC-1alpha expression in adipocytes to protect against diet-induced metabolic dysfunction. Sci Signal 12, doi:10.1126/scisignal.aau2281 (2019).PMC6882500

18 Erlich, A. T., Brownlee, D. M., Beyfuss, K. & Hood, D. A. Exercise induces TFEB expression and activity in skeletal muscle in a PGC-1α-dependent manner. American journal of physiology. Cell physiology 314, C62–c72, doi:10.1152/ajpcell.00162.2017 (2018).PMC5866381

19 Matthews, I. et al. Skeletal muscle TFEB signaling promotes central nervous system function and reduces neuroinflammation during aging and neurodegenerative disease. Cell Rep 42, 113436, doi:10.1016/j.celrep.2023.113436 (2023).PMC10841857

20 Taha, H. B. et al. Activation of the muscle-to-brain axis ameliorates neurocognitive deficits in an Alzheimer’s disease mouse model via enhancing neurotrophic and synaptic signaling. GeroScience, doi:10.1007/s11357-024-01345-3 (2024)

21 Patterson, D. C., Birnbaum, A., Matthews, I. & Cortes, C. J. Convergent skeletal muscle cytokine responses to TFEB overexpression and voluntary wheel running reflect sex-based variability to exercise adaptations in mice. Physiol Rep 13, e70519, doi:10.14814/phy2.70519 (2025).PMC12367098

22 Ishihara, A. et al. Effects of running exercise with increasing loads on tibialis anterior muscle fibres in mice. Exp Physiol 87, 113–116, doi:10.1113/eph8702340 (2002)

23 Valdez, G. et al. Attenuation of age-related changes in mouse neuromuscular synapses by caloric restriction and exercise. Proc Natl Acad Sci U S A 107, 14863–14868, doi:10.1073/pnas.1002220107 (2010).PMC2930485

24 Kelly, N. A. et al. Quantification and characterization of grouped type I myofibers in human aging. Muscle Nerve 57, E52–E59, doi:10.1002/mus.25711 (2018).PMC5711619

25 Roberts, B. M. et al. Human neuromuscular aging: Sex differences revealed at the myocellular level. Exp Gerontol 106, 116–124, doi:10.1016/j.exger.2018.02.023 (2018).PMC6031257

26 Giacomello, E. et al. Age Dependent Modification of the Metabolic Profile of the Tibialis Anterior Muscle Fibers in C57BL/6J Mice. Int J Mol Sci 21, doi:10.3390/ijms21113923 (2020).PMC7312486

27 Kim, M. et al. Single-nucleus transcriptomics reveals functional compartmentalization in syncytial skeletal muscle cells. Nature communications 11, 6375, doi:10.1038/s41467-020-20064-9 (2020).PMC7732842

28 Petrany, M. J. et al. Single-nucleus RNA-seq identifies transcriptional heterogeneity in multinucleated skeletal myofibers. Nature communications 11, 6374, doi:10.1038/s41467-020-20063-w (2020).PMC7733460

29 Santos, M. D. et al. Extraction and sequencing of single nuclei from murine skeletal muscles. STAR Protoc 2, 100694, doi:10.1016/j.xpro.2021.100694 (2021).PMC8339382

30 LaBarge, S. A. et al. p300 is not required for metabolic adaptation to endurance exercise training. FASEB J 30, 1623–1633, doi:10.1096/fj.15-281741 (2016).PMC4799503

31 Ikeda, S. et al. Muscle type-specific response of PGC-1 alpha and oxidative enzymes during voluntary wheel running in mouse skeletal muscle. Acta Physiol (Oxf) 188, 217–223, doi:10.1111/j.1748-1716.2006.01623.x (2006)

32 Waters, R. E., Rotevatn, S., Li, P., Annex, B. H. & Yan, Z. Voluntary running induces fiber type-specific angiogenesis in mouse skeletal muscle. American journal of physiology. Cell physiology 287, C1342–1348, doi:10.1152/ajpcell.00247.2004 (2004)

33 Allen, D. L. et al. Cardiac and skeletal muscle adaptations to voluntary wheel running in the mouse. J Appl Physiol (1985) 90, 1900–1908, doi:10.1152/jappl.2001.90.5.1900 (2001)

34 Svensson, M. et al. Forced treadmill exercise can induce stress and increase neuronal damage in a mouse model of global cerebral ischemia. Neurobiol Stress 5, 8–18, doi:10.1016/j.ynstr.2016.09.002 (2016).PMC5145912

35 Cook, M. D. et al. Forced treadmill exercise training exacerbates inflammation and causes mortality while voluntary wheel training is protective in a mouse model of colitis. Brain Behav Immun 33, 46–56, doi:10.1016/j.bbi.2013.05.005 (2013).PMC3775960

36 Meng, F. L. et al. singleCellBase: a high-quality manually curated database of cell markers for single cell annotation across multiple species. Biomark Res 11, 83, doi:10.1186/s40364-023-00523-3 (2023).PMC10510128

37 McKellar, D. W. et al. Large-scale integration of single-cell transcriptomic data captures transitional progenitor states in mouse skeletal muscle regeneration. Commun Biol 4, 1280, doi:10.1038/s42003-021-02810-x (2021).PMC8589952

38 Squair, J. W., Skinnider, M. A., Gautier, M., Foster, L. J. & Courtine, G. Prioritization of cell types responsive to biological perturbations in single-cell data with Augur. Nat Protoc 16, 3836–3873, doi:10.1038/s41596-021-00561-x (2021)

39 Skinnider, M. A. et al. Cell type prioritization in single-cell data. Nat Biotechnol 39, 30–34, doi:10.1038/s41587-020-0605-1 (2021).PMC7610525

40 Miller, S. A. et al. LSD1 and Aberrant DNA Methylation Mediate Persistence of Enteroendocrine Progenitors That Support BRAF-Mutant Colorectal Cancer. Cancer Res 81, 3791–3805, doi:10.1158/0008-5472.CAN-20-3562 (2021).PMC8513805

41 Jesus, I. et al. Effects of aerobic exercise training on muscle plasticity in a mouse model of cervical spinal cord injury. Sci Rep 11, 112, doi:10.1038/s41598-020-80478-9 (2021).PMC7794462

42 Olesen, A. T. et al. Age-related myofiber atrophy in old mice is reversed by ten weeks voluntary high-resistance wheel running. Exp Gerontol 143, 111150, doi:10.1016/j.exger.2020.111150 (2021)

43 Crowell, H. L. et al. muscat detects subpopulation-specific state transitions from multi-sample multi-condition single-cell transcriptomics data. Nature communications 11, 6077, doi:10.1038/s41467-020-19894-4 (2020).PMC7705760 D.C., and M.D.R. declare no competing interests.

44 Subramanian, A. et al. Gene set enrichment analysis: a knowledge-based approach for interpreting genome-wide expression profiles. Proc Natl Acad Sci U S A 102, 15545–15550, doi:10.1073/pnas.0506580102 (2005).PMC1239896

45 Liu, L. et al. Exercise reprograms the inflammatory landscape of multiple stem cell compartments during mammalian aging. Cell Stem Cell 30, 689–705 e684, doi:10.1016/j.stem.2023.03.016 (2023).PMC10216894

46 Tan-Chen, S., Guitton, J., Bourron, O., Le Stunff, H. & Hajduch, E. Sphingolipid Metabolism and Signaling in Skeletal Muscle: From Physiology to Physiopathology. Front Endocrinol (Lausanne) 11, 491, doi:10.3389/fendo.2020.00491 (2020).PMC7426366

47 Ahmad, K. et al. Extracellular matrix: the critical contributor to skeletal muscle regeneration-a comprehensive review. Inflamm Regen 43, 58, doi:10.1186/s41232-023-00308-z (2023).PMC10680355

48 Loreti, M. & Sacco, A. The jam session between muscle stem cells and the extracellular matrix in the tissue microenvironment. NPJ Regen Med 7, 16, doi:10.1038/s41536-022-00204-z (2022).PMC8854427

49 Rullman, E. et al. Endurance exercise activates matrix metalloproteinases in human skeletal muscle. J Appl Physiol (1985) 106, 804–812, doi:10.1152/japplphysiol.90872.2008 (2009)

50 Gumpenberger, M. et al. Remodeling the Skeletal Muscle Extracellular Matrix in Older Age-Effects of Acute Exercise Stimuli on Gene Expression. Int J Mol Sci 21, doi:10.3390/ijms21197089 (2020).PMC7583913

51 Sternlicht, M. D. & Werb, Z. How matrix metalloproteinases regulate cell behavior. Annu Rev Cell Dev Biol 17, 463–516, doi:10.1146/annurev.cellbio.17.1.463 (2001).PMC2792593

52 Birot, O. J., Koulmann, N., Peinnequin, A. & Bigard, X. A. Exercise-induced expression of vascular endothelial growth factor mRNA in rat skeletal muscle is dependent on fibre type. J Physiol 552, 213–221, doi:10.1113/jphysiol.2003.043026 (2003).PMC2343332

53 Delavar, H. et al. Skeletal myofiber VEGF is essential for the exercise training response in adult mice. Am J Physiol Regul Integr Comp Physiol 306, R586–595, doi:10.1152/ajpregu.00522.2013 (2014).PMC4043130

54 Lee, S., Jilani, S. M., Nikolova, G. V., Carpizo, D. & Iruela-Arispe, M. L. Processing of VEGF-A by matrix metalloproteinases regulates bioavailability and vascular patterning in tumors. J Cell Biol 169, 681–691, doi:10.1083/jcb.200409115 (2005).PMC2171712

55 Stamenkovic, I. Extracellular matrix remodelling: the role of matrix metalloproteinases. The Journal of pathology 200, 448–464, doi:10.1002/path.1400 (2003)

56 Arnold, A. S. et al. Morphological and functional remodelling of the neuromuscular junction by skeletal muscle PGC-1alpha. Nature communications 5, 3569, doi:10.1038/ncomms4569 (2014).PMC4846352

57 Ortlund, E. et al. Endurance Exercise Training Alters Lipidomic Profiles of Plasma and Eight Tissues in Rats: a MoTrPAC study. Res Sq, doi:10.21203/rs.3.rs-5263273/v1 (2024).PMC11601870

58 Fan, Z. et al. Exercise-induced angiogenesis is dependent on metabolically primed ATF3/4(+) endothelial cells. Cell Metab 33, 1793–1807 e1799, doi:10.1016/j.cmet.2021.07.015 (2021).PMC8432967

59 Green, D. J., Maiorana, A., O’Driscoll, G. & Taylor, R. Effect of exercise training on endothelium-derived nitric oxide function in humans. J Physiol 561, 1–25, doi:10.1113/jphysiol.2004.068197 (2004).PMC1665322

60 McKoy, G. et al. Expression of Ankrd2 in fast and slow muscles and its response to stretch are consistent with a role in slow muscle function. J Appl Physiol (1985) 98, 2337–2343; discussion 2320, doi:10.1152/japplphysiol.01046.2004 (2005)

61 Bean, C. et al. Ankrd2 is a modulator of NF-kappaB-mediated inflammatory responses during muscle differentiation. Cell Death Dis 5, e1002, doi:10.1038/cddis.2013.525 (2014).PMC4040671

62 Oprescu, S. N., Yue, F., Qiu, J., Brito, L. F. & Kuang, S. Temporal Dynamics and Heterogeneity of Cell Populations during Skeletal Muscle Regeneration. iScience 23, 100993, doi:10.1016/j.isci.2020.100993 (2020).PMC7125354

63 Sexton, W. L. Vascular adaptations in rat hindlimb skeletal muscle after voluntary running-wheel exercise. J Appl Physiol (1985) 79, 287–296, doi:10.1152/jappl.1995.79.1.287 (1995)

64 Triolo, M., Oliveira, A. N., Kumari, R. & Hood, D. A. The influence of age, sex, and exercise on autophagy, mitophagy, and lysosome biogenesis in skeletal muscle. Skelet Muscle 12, 13, doi:10.1186/s13395-022-00296-7 (2022).PMC9188089

65 Turkel, I. et al. Exercise and Metabolic Health: The Emerging Roles of Novel Exerkines. Curr Protein Pept Sci 23, 437–455, doi:10.2174/1389203723666220629163524 (2022)

66 Distefano, G. & Goodpaster, B. H. Effects of Exercise and Aging on Skeletal Muscle. Cold Spring Harbor perspectives in medicine 8, doi:10.1101/cshperspect.a029785 (2018).PMC5830901

67 Hargreaves, M. & Spriet, L. L. Skeletal muscle energy metabolism during exercise. Nat Metab 2, 817–828, doi:10.1038/s42255-020-0251-4 (2020)

68 Kiens, B. Skeletal muscle lipid metabolism in exercise and insulin resistance. Physiol Rev 86, 205–243, doi:10.1152/physrev.00023.2004 (2006)

69 Tunstall, R. J. et al. Exercise training increases lipid metabolism gene expression in human skeletal muscle. American journal of physiology. Endocrinology and metabolism 283, E66–72, doi:10.1152/ajpendo.00475.2001 (2002)

70 Meex, R. C. et al. Restoration of muscle mitochondrial function and metabolic flexibility in type 2 diabetes by exercise training is paralleled by increased myocellular fat storage and improved insulin sensitivity. Diabetes 59, 572–579, doi:10.2337/db09-1322 (2010).PMC2828651

71 Goodpaster, B. H., He, J., Watkins, S. & Kelley, D. E. Skeletal muscle lipid content and insulin resistance: evidence for a paradox in endurance-trained athletes. J Clin Endocrinol Metab 86, 5755–5761, doi:10.1210/jcem.86.12.8075 (2001)

72 Schenk, S. & Horowitz, J. F. Acute exercise increases triglyceride synthesis in skeletal muscle and prevents fatty acid-induced insulin resistance. J Clin Invest 117, 1690–1698, doi:10.1172/JCI30566 (2007).PMC1866251

73 Bosma, M. Lipid droplet dynamics in skeletal muscle. Exp Cell Res 340, 180–186, doi:10.1016/j.yexcr.2015.10.023 (2016)

74 Kiens, B. et al. Lipid-binding proteins and lipoprotein lipase activity in human skeletal muscle: influence of physical activity and gender. J Appl Physiol (1985) 97, 1209–1218, doi:10.1152/japplphysiol.01278.2003 (2004)

75 Tarnopolsky, M. A. et al. Influence of endurance exercise training and sex on intramyocellular lipid and mitochondrial ultrastructure, substrate use, and mitochondrial enzyme activity. Am J Physiol Regul Integr Comp Physiol 292, R1271–1278, doi:10.1152/ajpregu.00472.2006 (2007)

76 Carter, S. L., Rennie, C. & Tarnopolsky, M. A. Substrate utilization during endurance exercise in men and women after endurance training. American journal of physiology. Endocrinology and metabolism 280, E898–907, doi:10.1152/ajpendo.2001.280.6.E898 (2001)

77 Steffensen, C. H., Roepstorff, C., Madsen, M. & Kiens, B. Myocellular triacylglycerol breakdown in females but not in males during exercise. Am J Physiol Endocrinol Metab 282, E634–642, doi:10.1152/ajpendo.00078.2001 (2002)

78 Horton, T. J., Pagliassotti, M. J., Hobbs, K. & Hill, J. O. Fuel metabolism in men and women during and after long-duration exercise. J Appl Physiol (1985) 85, 1823–1832, doi:10.1152/jappl.1998.85.5.1823 (1998)

79 Venables, M. C., Achten, J. & Jeukendrup, A. E. Determinants of fat oxidation during exercise in healthy men and women: a cross-sectional study. J Appl Physiol (1985) 98, 160–167, doi:10.1152/japplphysiol.00662.2003 (2005)

80 Friedlander, A. L., Casazza, G. A., Horning, M. A., Buddinger, T. F. & Brooks, G. A. Effects of exercise intensity and training on lipid metabolism in young women. Am J Physiol 275, E853–863, doi:10.1152/ajpendo.1998.275.5.E853 (1998)

81 Roepstorff, C. et al. Gender differences in substrate utilization during submaximal exercise in endurance-trained subjects. American journal of physiology. Endocrinology and metabolism 282, E435–447, doi:10.1152/ajpendo.00266.2001 (2002)

82 Fritzen, A. M., Lundsgaard, A. M. & Kiens, B. Tuning fatty acid oxidation in skeletal muscle with dietary fat and exercise. Nat Rev Endocrinol 16, 683–696, doi:10.1038/s41574-020-0405-1 (2020)

83 Hendlinger, M. et al. Exercise training increases skeletal muscle sphingomyelinases and affects mitochondrial quality control in men with type 2 diabetes. Metabolism 172, 156361, doi:10.1016/j.metabol.2025.156361 (2025)

84 Sato, A. Y. et al. Glucocorticoids Induce Bone and Muscle Atrophy by Tissue-Specific Mechanisms Upstream of E3 Ubiquitin Ligases. Endocrinology 158, 664–677, doi:10.1210/en.2016-1779 (2017).PMC5460781

85 Kuo, T., Harris, C. A. & Wang, J. C. Metabolic functions of glucocorticoid receptor in skeletal muscle. Mol Cell Endocrinol 380, 79–88, doi:10.1016/j.mce.2013.03.003 (2013).PMC4893778

86 Namkung, J., Kim, H. & Park, S. Peripheral Serotonin: a New Player in Systemic Energy Homeostasis. Mol Cells 38, 1023–1028, doi:10.14348/molcells.2015.0258 (2015).PMC4696992

87 Falabregue, M. et al. Lack of Skeletal Muscle Serotonin Impairs Physical Performance. Int J Tryptophan Res 14, 11786469211003109, doi:10.1177/11786469211003109 (2021).PMC7989111

88 Park, S. et al. Inhibition of serotonin-Htr2b signaling in skeletal muscle mitigates obesity-induced insulin resistance. Exp Mol Med 57, 1177–1188, doi:10.1038/s12276-025-01460-x (2025).PMC12227683

89 Fefeu, M. et al. Serotonin reuptake inhibitors improve muscle stem cell function and muscle regeneration in male mice. Nature communications 15, 6457, doi:10.1038/s41467-024-50220-4 (2024).PMC11291725 Janssen, Lundbeck, Novartis, Roche, SOBI, Takeda. He has served as a consultant and/or a speaker for Astra Zeneca, Boehringer-Ingelheim, Pierre Fabre, Eli Lilly, Lundbeck, LVMH, MAPREG, Novartis, Otsuka, Pileje, SANOFI, Servier and received compensation, and he has received research support from Servier. He is a board member of the Regstem company. O.M. has received consultancy fees from Astra-Zeneca, Blueprint Medicines, Boehringer-Ingelheim, Bristol-Myers Squibb, Eli Lilly, Ipsen, Merck Sharpe & Dohme, Pfizer, Roche, Servier, and Vifor Pharma. O.M. is an employee and shareholder of Amgen since Feb 1st, 2022. FCh is a board member of the Regstem company. The remaining authors have nothing to disclose.

90 Dudley, A. C. & Griffioen, A. W. The modes of angiogenesis: an updated perspective. Angiogenesis 26, 477–480, doi:10.1007/s10456-023-09895-4 (2023).PMC10777330

91 Miniou, P. et al. Gene targeting restricted to mouse striated muscle lineage. Nucleic Acids Res 27, e27, doi:gnc027 [pii] (1999).148637

92 Medina, D. L. et al. Transcriptional activation of lysosomal exocytosis promotes cellular clearance. Developmental cell 21, 421–430, doi:10.1016/j.devcel.2011.07.016 (2011).3173716

93 Wei, W. et al. Organism-wide, cell-type-specific secretome mapping of exercise training in mice. Cell Metab 35, 1261–1279 e1211, doi:10.1016/j.cmet.2023.04.011 (2023).PMC10524249

94 Whitham, M. et al. Extracellular Vesicles Provide a Means for Tissue Crosstalk during Exercise. Cell Metab 27, 237–251 e234, doi:10.1016/j.cmet.2017.12.001 (2018)

95 McGill, C. J., Ewald, C. Y. & Benayoun, B. A. Sex-dimorphic expression of extracellular matrix genes in mouse bone marrow neutrophils. PLoS One 18, e0294859, doi:10.1371/journal.pone.0294859 (2023).PMC10688658

96 Simon, L. R., Scott, A. J., Figueroa Rios, L., Zembles, J. & Masters, K. S. Cellular-scale sex differences in extracellular matrix remodeling by valvular interstitial cells. Heart Vessels 38, 122–130, doi:10.1007/s00380-022-02164-2 (2023).PMC10120251

97 Li, Y. et al. Cell sex affects extracellular matrix protein expression and proliferation of smooth muscle progenitor cells derived from human pluripotent stem cells. Stem Cell Res Ther 8, 156, doi:10.1186/s13287-017-0606-2 (2017).PMC5496346

98 Schweitzer, A. M., Koehle, M. S., Fliss, M. D. & Mitchell, C. J. Collagen remodeling increases after acute resistance exercise in healthy skeletal muscle irrespective of age. American journal of physiology. Cell physiology 329, C68–C81, doi:10.1152/ajpcell.00992.2024 (2025)

99 Janowski, A. J. et al. The influence of sex on activity in voluntary wheel running, forced treadmill running, and open field testing in mice. Physiol Rep 13, e70246, doi:10.14814/phy2.70246 (2025).PMC11845322

100 Bartling, B. et al. Sex-related differences in the wheel-running activity of mice decline with increasing age. Exp Gerontol 87, 139–147, doi:10.1016/j.exger.2016.04.011 (2017)

101 Manzanares, G., Brito-da-Silva, G. & Gandra, P. G. Voluntary wheel running: patterns and physiological effects in mice. Braz J Med Biol Res 52, e7830, doi:10.1590/1414-431X20187830 (2018).PMC6301263

102 Cabelka, C. A. et al. Effects of ovarian hormones and estrogen receptor alpha on physical activity and skeletal muscle fatigue in female mice. Exp Gerontol 115, 155–164, doi:10.1016/j.exger.2018.11.003 (2019).PMC6331238

103 Rhodes, J. S., Gammie, S. C. & Garland, T., Jr. Neurobiology of Mice Selected for High Voluntary Wheel-running Activity. Integr Comp Biol 45, 438–455, doi:10.1093/icb/45.3.438 (2005)

104 Waters, R. P. et al. Selection for increased voluntary wheel-running affects behavior and brain monoamines in mice. Brain Res 1508, 9–22, doi:10.1016/j.brainres.2013.01.033 (2013).PMC3660142

105 Lightfoot, J. T. Sex hormones’ regulation of rodent physical activity: a review. Int J Biol Sci 4, 126–132, doi:10.7150/ijbs.4.126 (2008).PMC2359866

106 Jones, E. F., Howton, T. C., Flanary, V. L., Clark, A. D. & Lasseigne, B. N. Long-read RNA sequencing identifies region- and sex-specific C57BL/6J mouse brain mRNA isoform expression and usage. Mol Brain 17, 40, doi:10.1186/s13041-024-01112-7 (2024).PMC11188239

107 Fisher, J. L., Clark, A. D., Jones, E. F. & Lasseigne, B. N. Sex-biased gene expression and gene-regulatory networks of sex-biased adverse event drug targets and drug metabolism genes. BMC Pharmacol Toxicol 25, 5, doi:10.1186/s40360-023-00727-1 (2024).PMC10763002

108 Gilda, J. E. et al. A semiautomated measurement of muscle fiber size using the Imaris software. American journal of physiology. Cell physiology 321, C615–C631, doi:10.1152/ajpcell.00206.2021 (2021)

109 Bray, N. L., Pimentel, H., Melsted, P. & Pachter, L. Near-optimal probabilistic RNA-seq quantification. Nat Biotechnol 34, 525–527, doi:10.1038/nbt.3519 (2016)

110 Yang, S. et al. Decontamination of ambient RNA in single-cell RNA-seq with DecontX. Genome Biol 21, 57, doi:10.1186/s13059-020-1950-6 (2020).PMC7059395

111 Kowalczyk, M. S. et al. Single-cell RNA-seq reveals changes in cell cycle and differentiation programs upon aging of hematopoietic stem cells. Genome Res 25, 1860–1872, doi:10.1101/gr.192237.115 (2015).PMC4665007

112 McGinnis, C. S., Murrow, L. M. & Gartner, Z. J. DoubletFinder: Doublet Detection in Single-Cell RNA Sequencing Data Using Artificial Nearest Neighbors. Cell Syst 8, 329–337 e324, doi:10.1016/j.cels.2019.03.003 (2019).PMC6853612

113 Bais, A. S. & Kostka, D. scds: computational annotation of doublets in single-cell RNA sequencing data. Bioinformatics 36, 1150–1158, doi:10.1093/bioinformatics/btz698 (2020).PMC7703774

114 Guo, H. & Li, J. scSorter: assigning cells to known cell types according to marker genes. Genome Biol 22, 69, doi:10.1186/s13059-021-02281-7 (2021).PMC7898451

115 Zhang, Y., Zhang, F., Wang, Z., Wu, S. & Tian, W. scMAGIC: accurately annotating single cells using two rounds of reference-based classification. Nucleic Acids Res 50, e43, doi:10.1093/nar/gkab1275 (2022).PMC9071478

116 McCarthy, D. J., Campbell, K. R., Lun, A. T. & Wills, Q. F. Scater: pre-processing, quality control, normalization and visualization of single-cell RNA-seq data in R. Bioinformatics 33, 1179–1186, doi:10.1093/bioinformatics/btw777 (2017).PMC5408845

117 Leek, J. T., Johnson, W. E., Parker, H. S., Jaffe, A. E. & Storey, J. D. The sva package for removing batch effects and other unwanted variation in high-throughput experiments. Bioinformatics 28, 882–883, doi:10.1093/bioinformatics/bts034 (2012).PMC3307112

118 Love, M. I., Huber, W. & Anders, S. Moderated estimation of fold change and dispersion for RNA-seq data with DESeq2. Genome Biol 15, 550, doi:10.1186/s13059-014-0550-8 (2014).PMC4302049

119 Liberzon, A. et al. Molecular signatures database (MSigDB) 3.0. Bioinformatics 27, 1739–1740, doi:10.1093/bioinformatics/btr260 (2011).PMC3106198

120 Wang, M. & Han, X. Multidimensional mass spectrometry-based shotgun lipidomics. Methods Mol Biol 1198, 203–220, doi:10.1007/978-1-4939-1258-2_13 (2014).PMC4261229

121 Han, X., Yang, K., Cheng, H., Fikes, K. N. & Gross, R. W. Shotgun lipidomics of phosphoethanolamine-containing lipids in biological samples after one-step in situ derivatization. J Lipid Res 46, 1548–1560, doi:10.1194/jlr.D500007-JLR200 (2005).PMC2141546

122 Cheng, H., Jiang, X. & Han, X. Alterations in lipid homeostasis of mouse dorsal root ganglia induced by apolipoprotein E deficiency: a shotgun lipidomics study. J Neurochem 101, 57–76, doi:10.1111/j.1471-4159.2006.04342.x (2007).PMC2137162

123 Wang, M., Han, R. H. & Han, X. Fatty acidomics: global analysis of lipid species containing a carboxyl group with a charge-remote fragmentation-assisted approach. Anal Chem 85, 9312–9320, doi:10.1021/ac402078p (2013).PMC3826454

124 Wang, M., Fang, H. & Han, X. Shotgun lipidomics analysis of 4-hydroxyalkenal species directly from lipid extracts after one-step in situ derivatization. Anal Chem 84, 4580–4586, doi:10.1021/ac300695p (2012).PMC3352972

125 Han, X., Yang, K. & Gross, R. W. Microfluidics-based electrospray ionization enhances the intrasource separation of lipid classes and extends identification of individual molecular species through multi-dimensional mass spectrometry: development of an automated high-throughput platform for shotgun lipidomics. Rapid Commun Mass Spectrom 22, 2115–2124, doi:10.1002/rcm.3595 (2008).PMC2927983

126 Wang, M., Wang, C., Han, R. H. & Han, X. Novel advances in shotgun lipidomics for biology and medicine. Prog Lipid Res 61, 83–108, doi:10.1016/j.plipres.2015.12.002 (2016).PMC4733395

127 Yang, K., Cheng, H., Gross, R. W. & Han, X. Automated lipid identification and quantification by multidimensional mass spectrometry-based shotgun lipidomics. Anal Chem 81, 4356–4368, doi:10.1021/ac900241u (2009).PMC2728582

128 Sanford, J. A. et al. Molecular Transducers of Physical Activity Consortium (MoTrPAC): Mapping the Dynamic Responses to Exercise. Cell 181, 1464–1474, doi:10.1016/j.cell.2020.06.004 (2020)

