## Supplementary Figures and legends for "Single-nuclei RNA-sequencing uncovers sexually divergent exercise signatures partially mimicked by TFEB overexpression in mouse skeletal muscle"

### 2026 Supplementary Figure S1

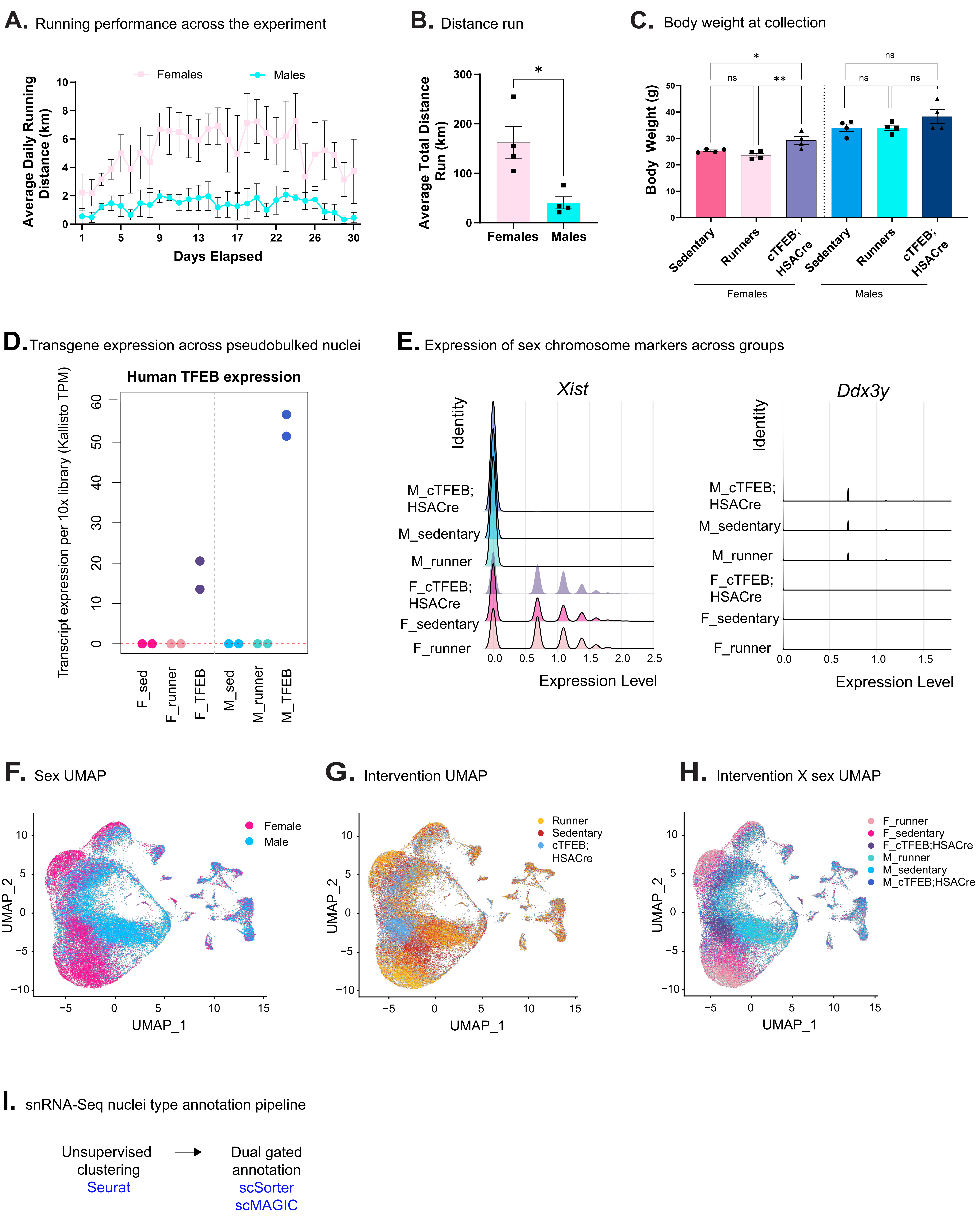

### Supplementary Figure S2

**A.** Representative images of TA muscle whole section fybertyping

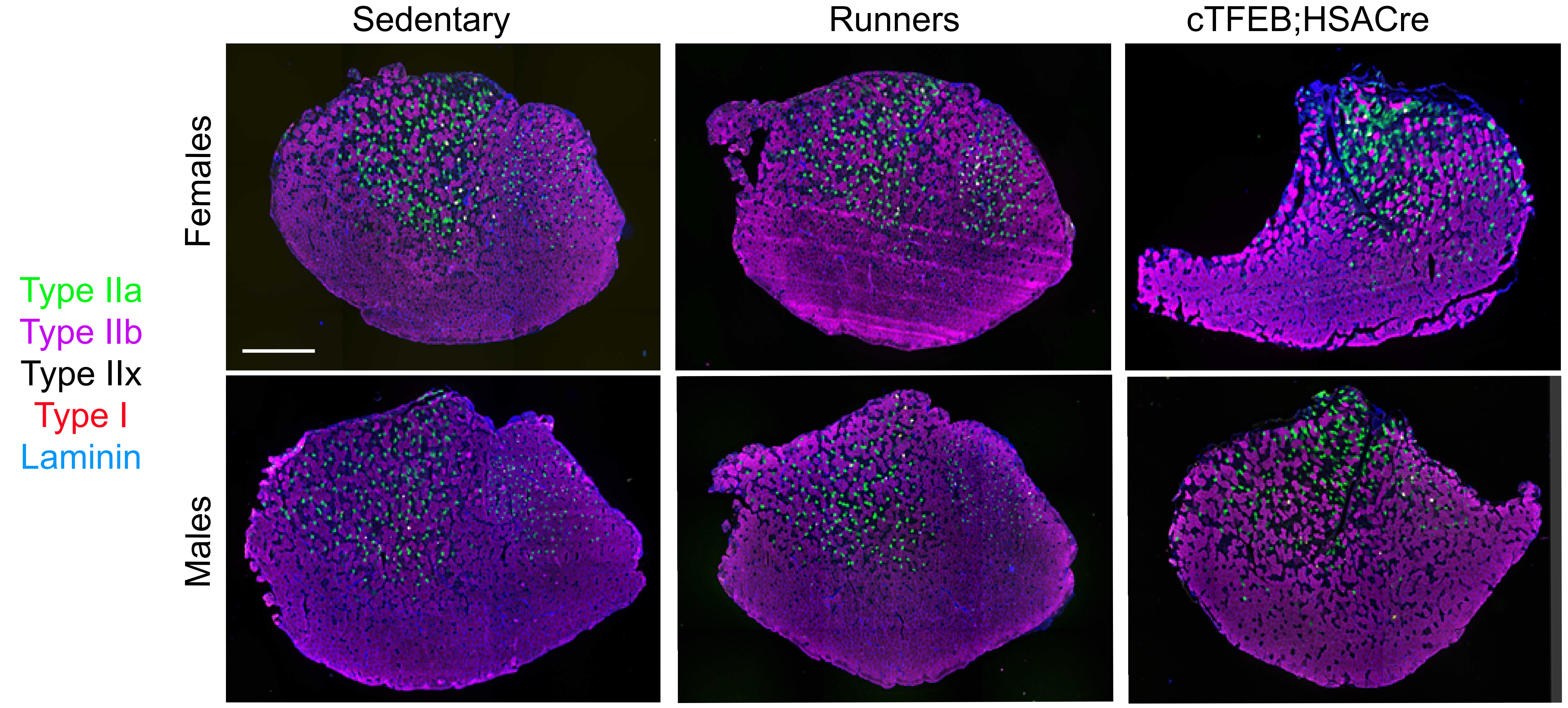


**B.** Cross-sectional area analyis per fiber type

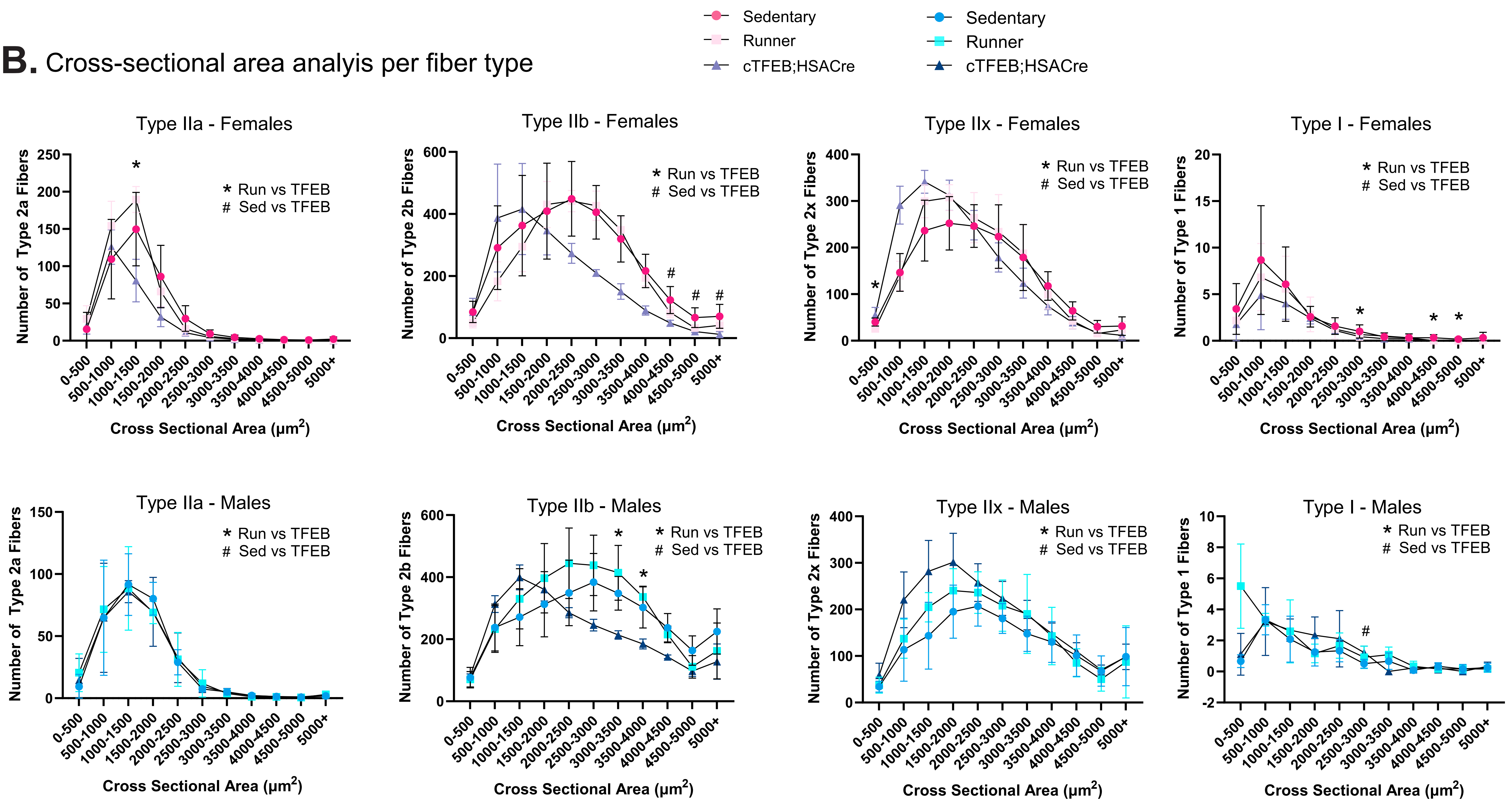

### 2026 Supplementary Figure S3

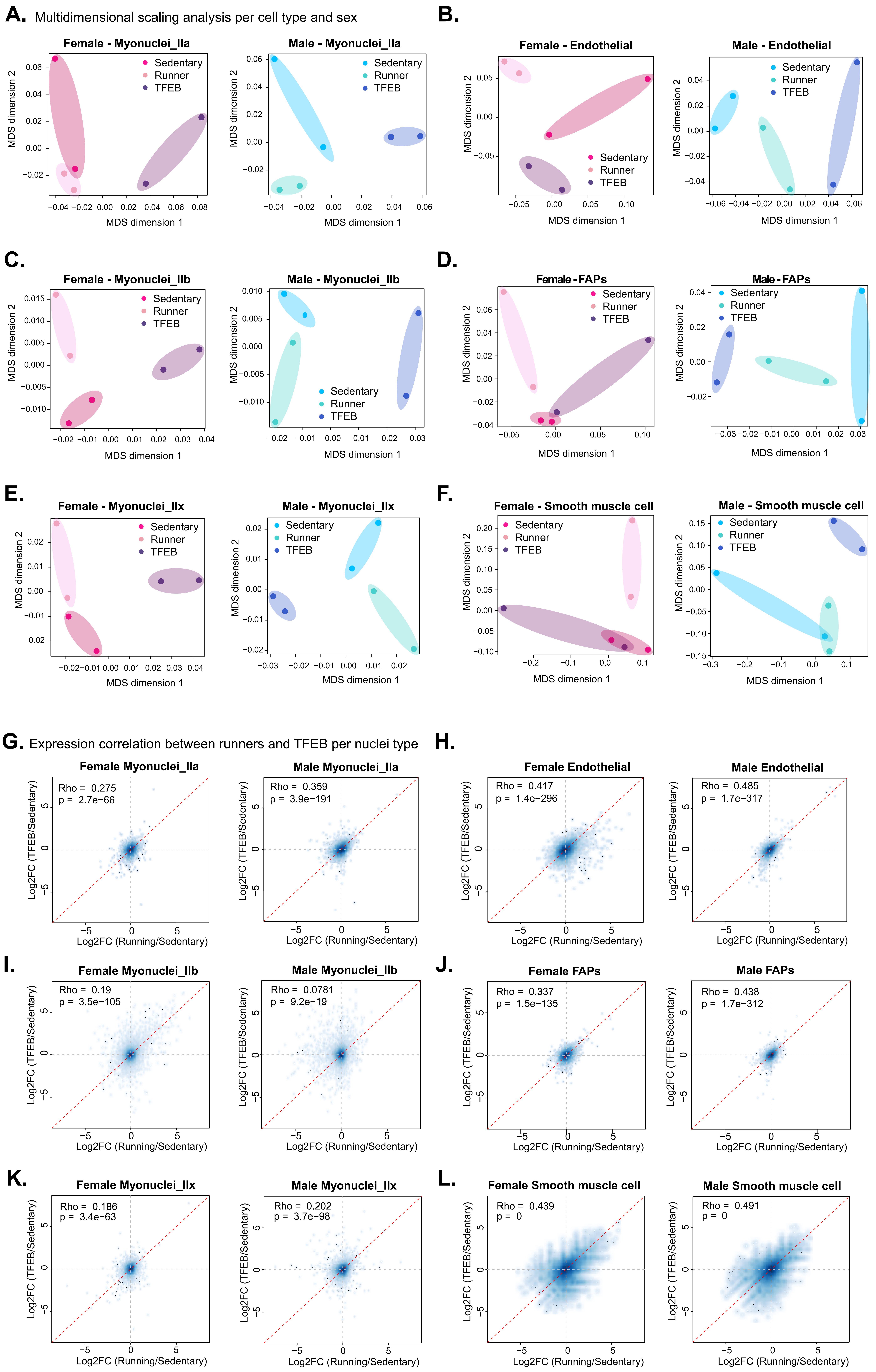

### 2026 Supplementary Figure S4

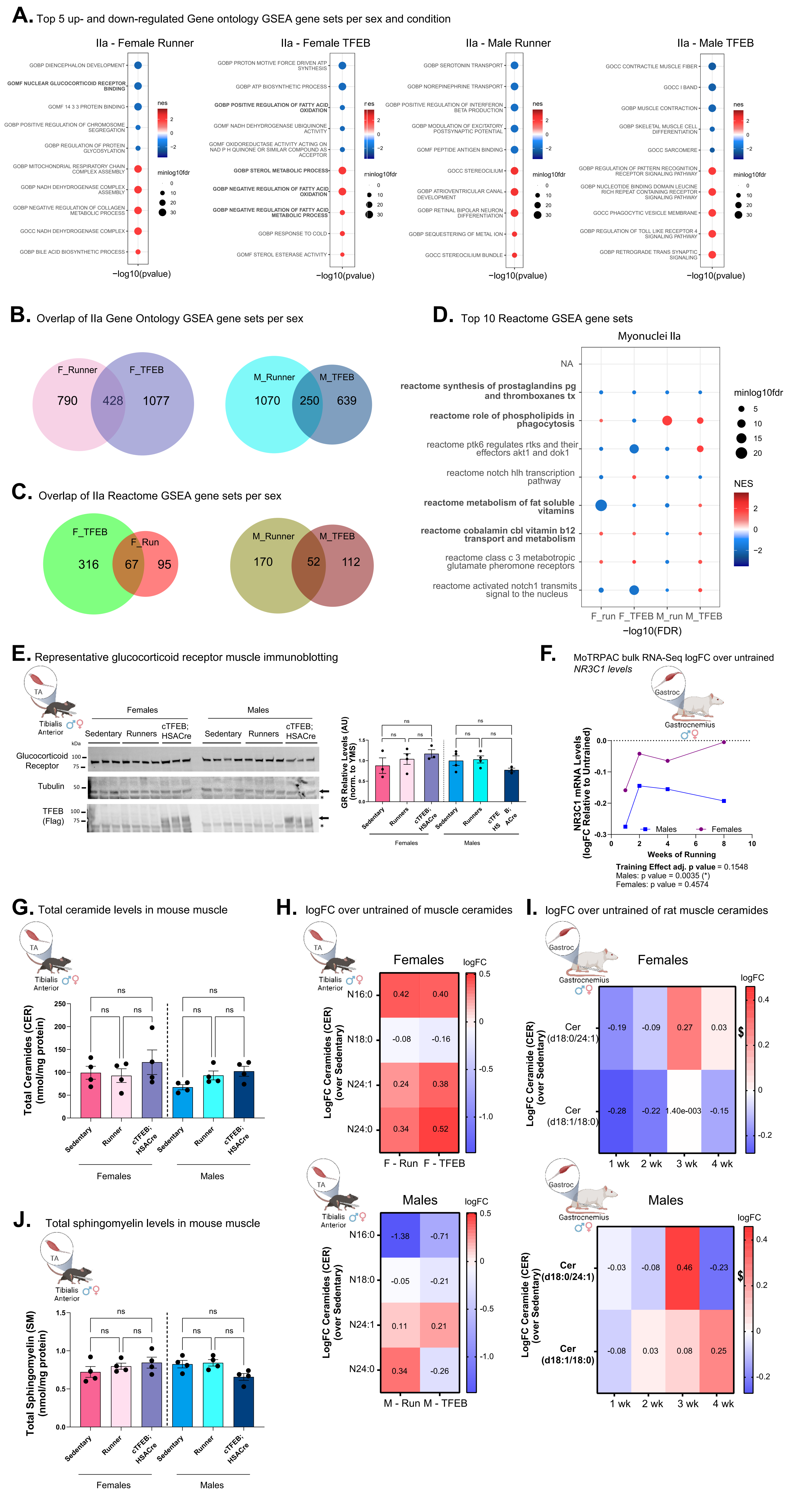

### Supplementary Figure S5

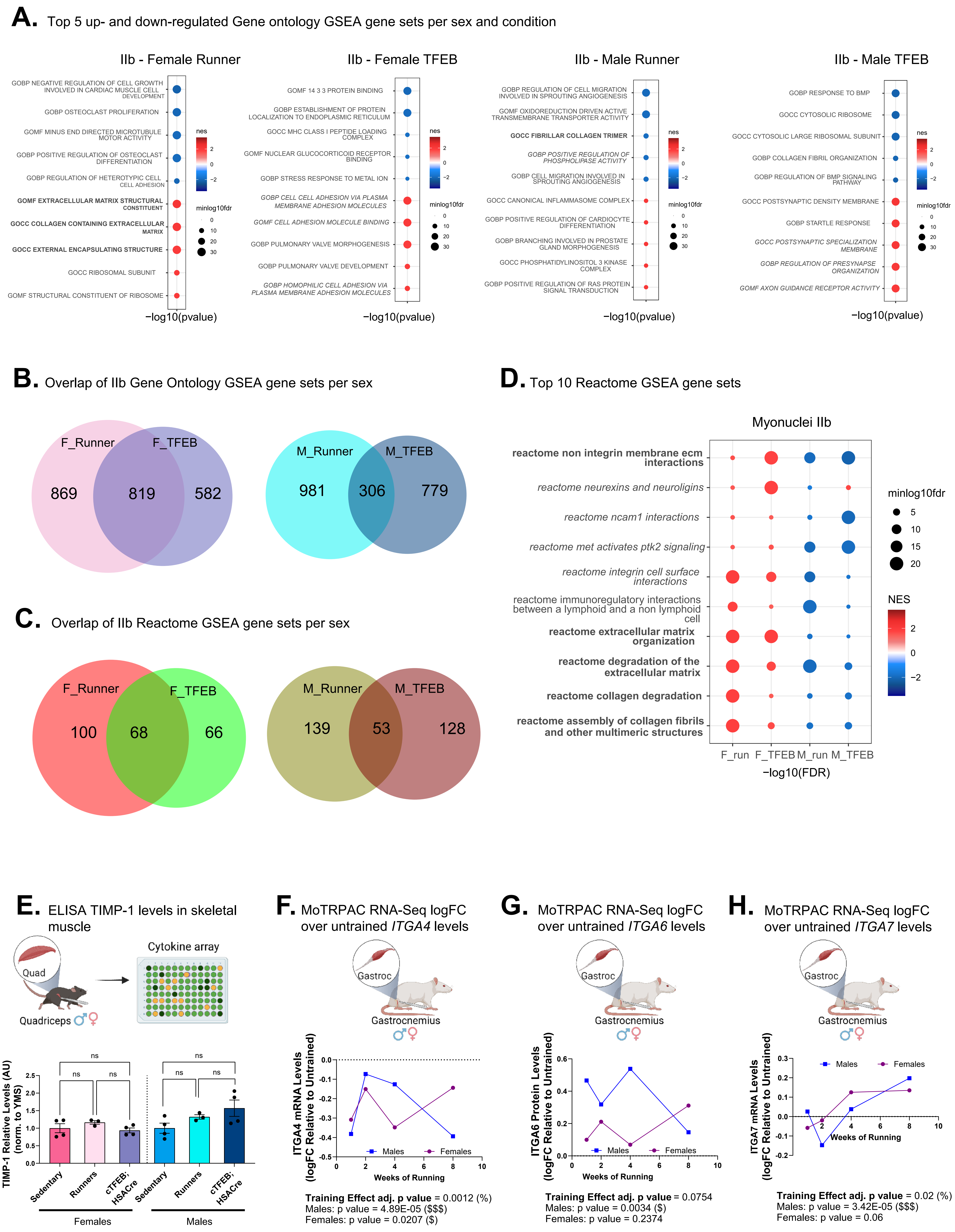

### Supplementary Figure S6

**A.** Top 5 up- and down-regulated Gene ontology GSEA gene sets per sex and condition

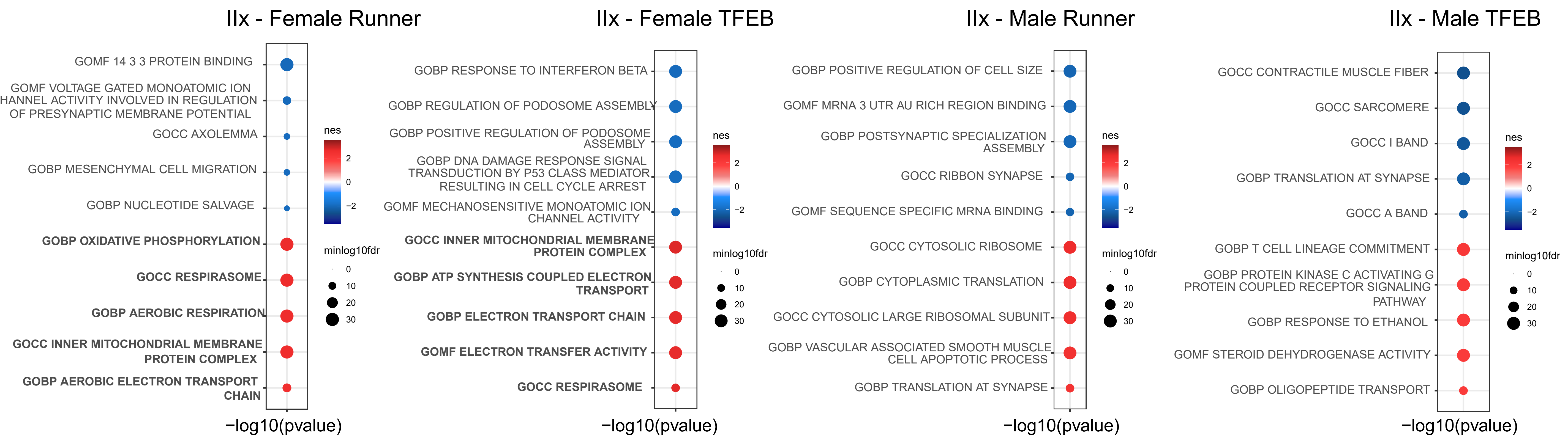


**B.** Overlap of IIx Gene Ontology GSEA gene sets per sex

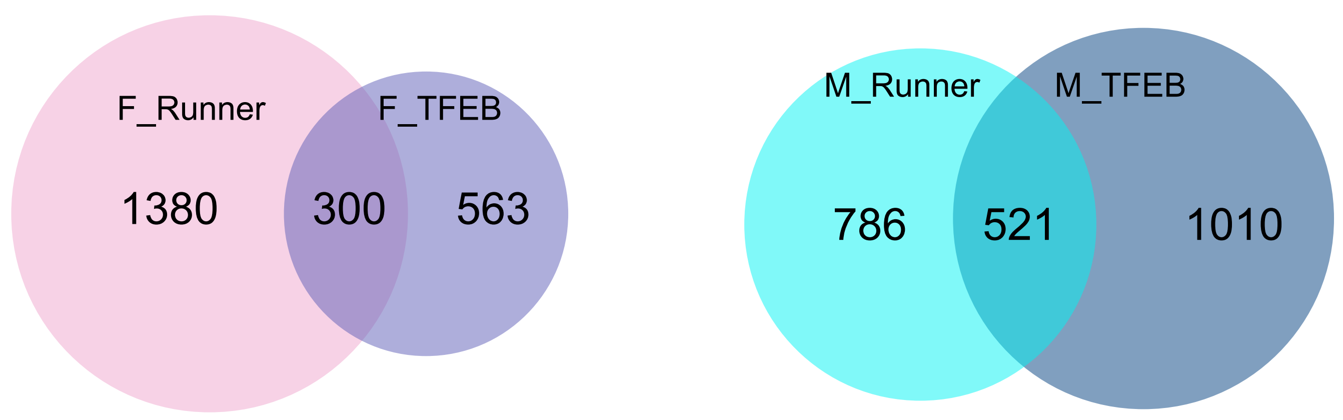


**D.** Top 10 Reactome GSEA gene sets

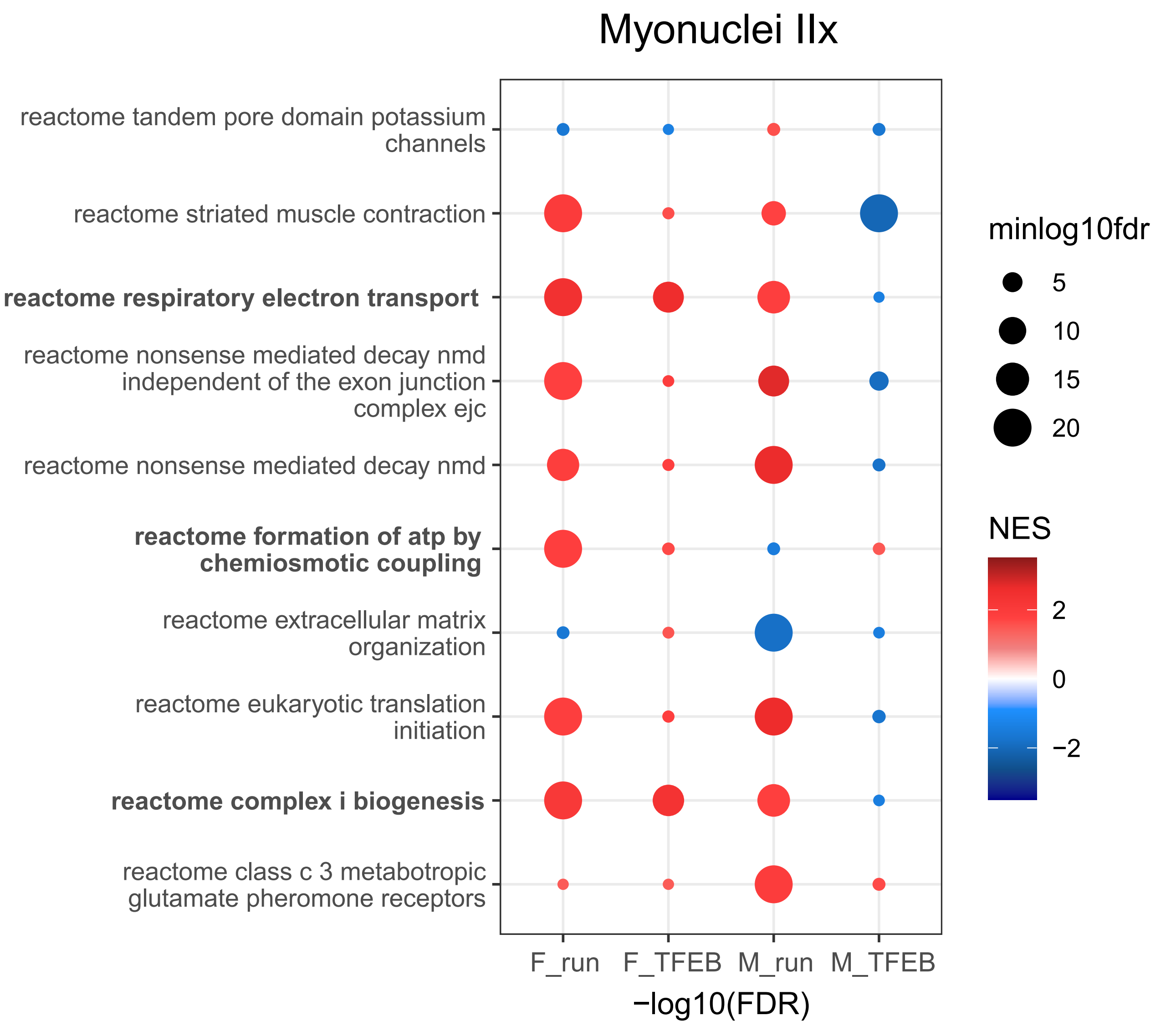


**C.** Overlap of IIx Reactome GSEA gene sets per sex

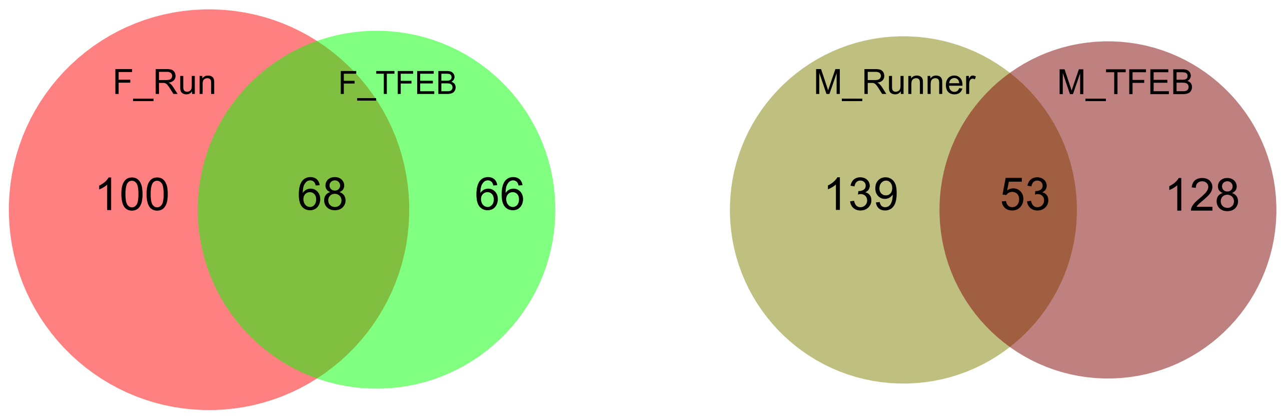


**E.** Total acyl-carnitine levels in mouse muscle

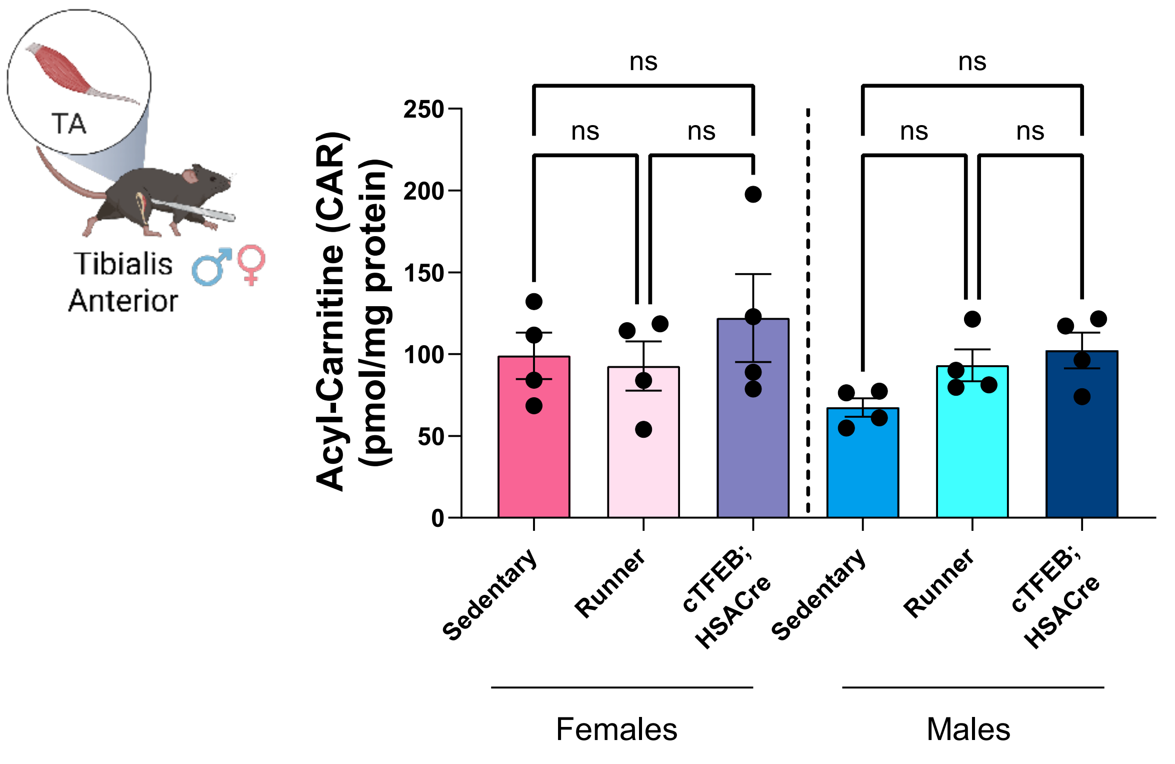


**F.** Total free fatty acid levels in mouse muscle

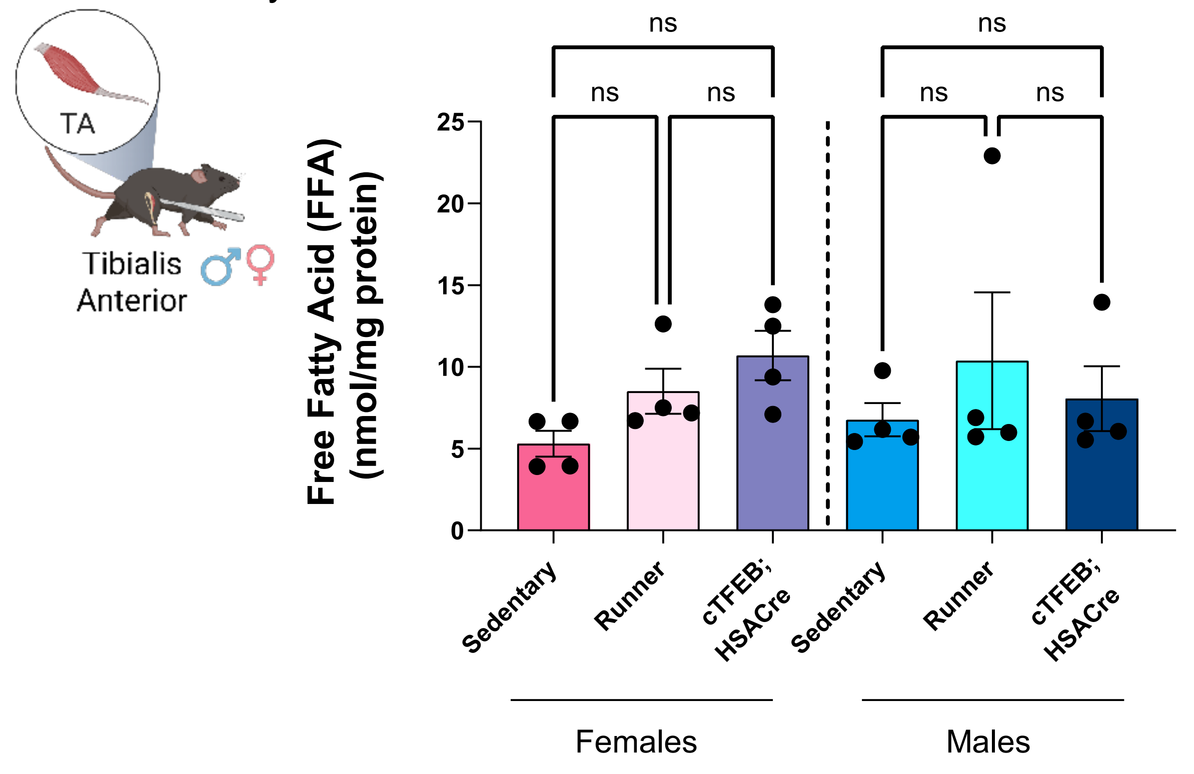


**G.** Total fatty acyl chains levels in mouse muscle

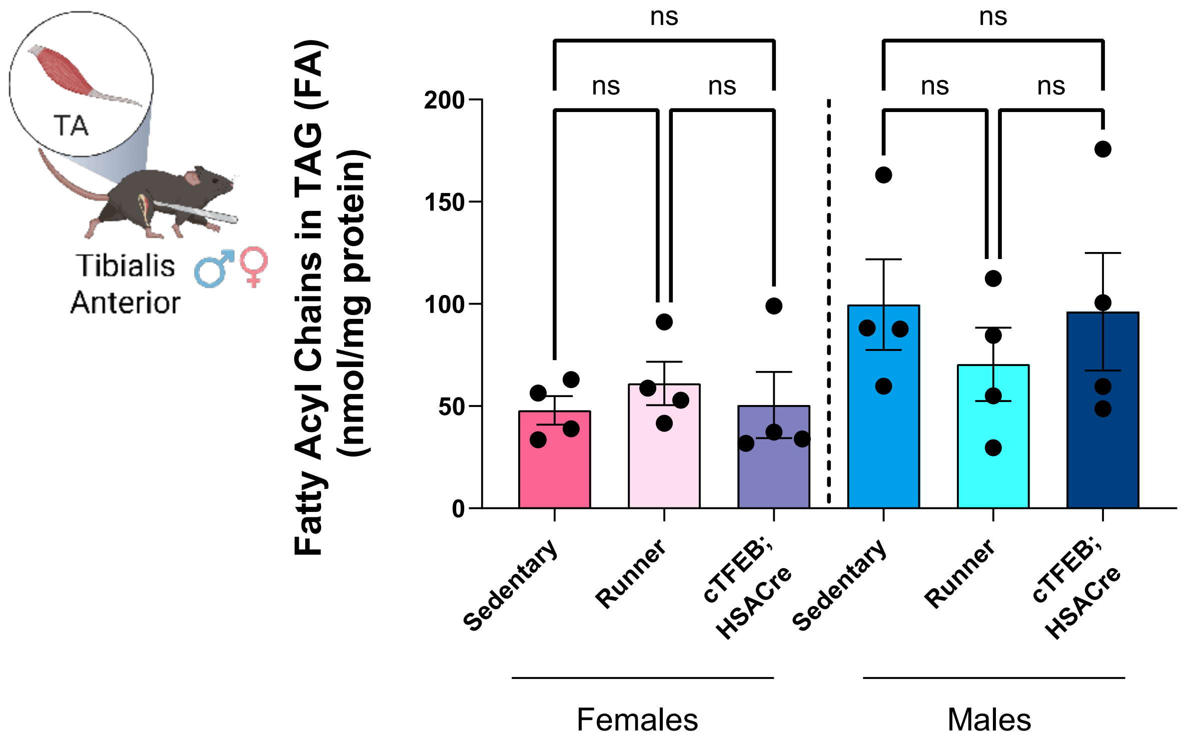


**H.** Total 4-hydroxynonenal levels in mouse muscle

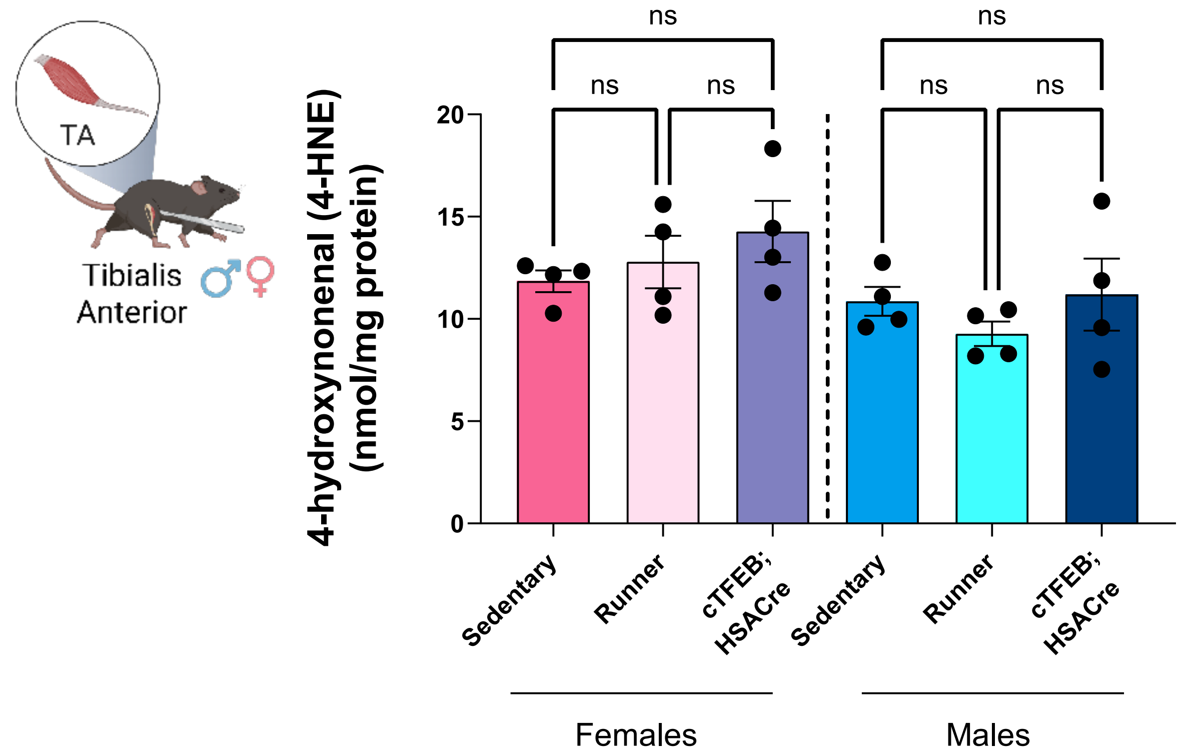

### Supplementary Figure S7

**A.** Top 5 up- and down-regulated Gene ontology GSEA gene sets per sex and condition

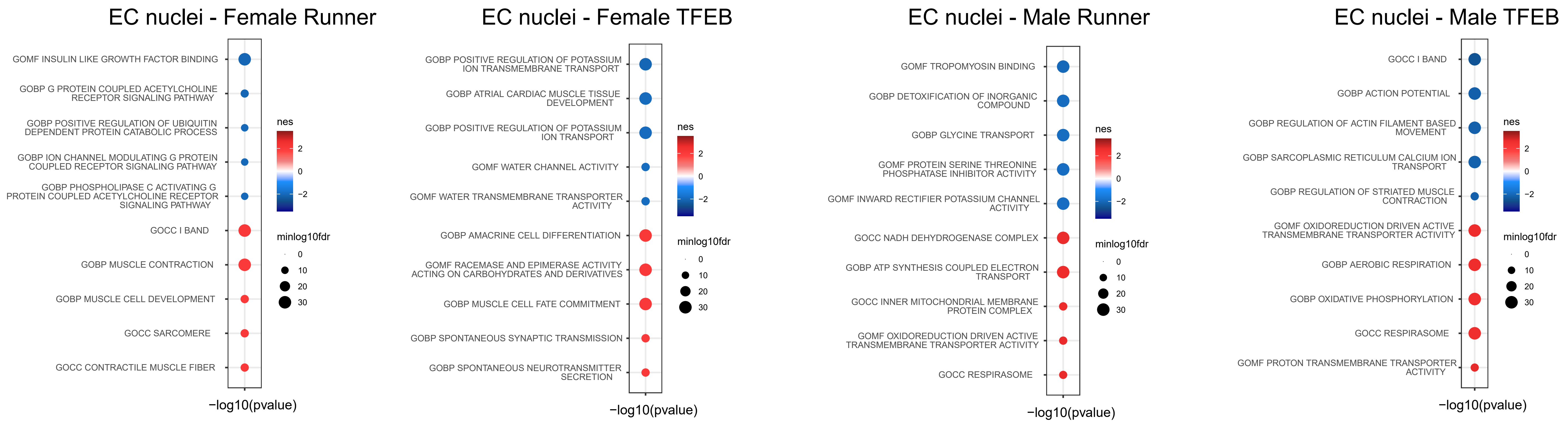


**B.** Overlap of EC nuclei Gene Ontology GSEA gene sets per sex

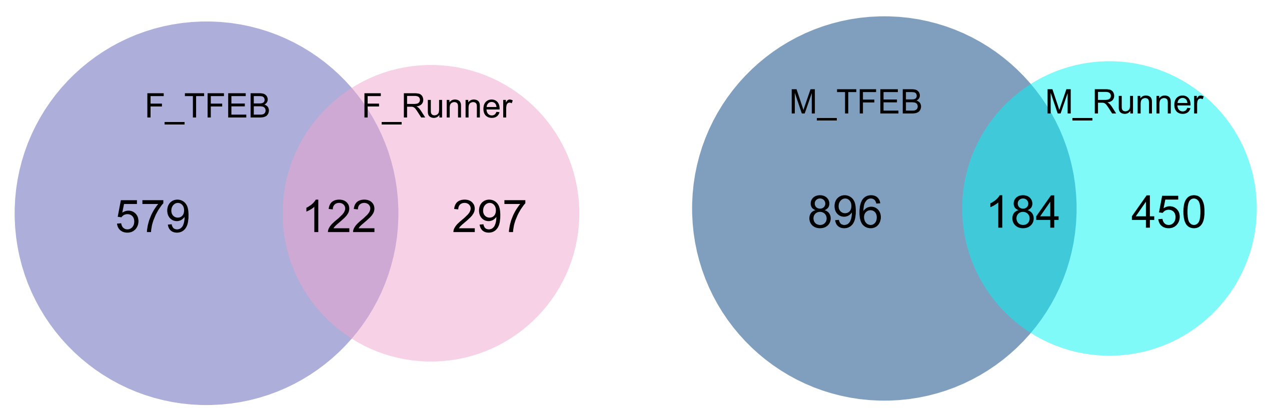


**C.** Overlap of EC nuclei Reactome GSEA gene sets per sex

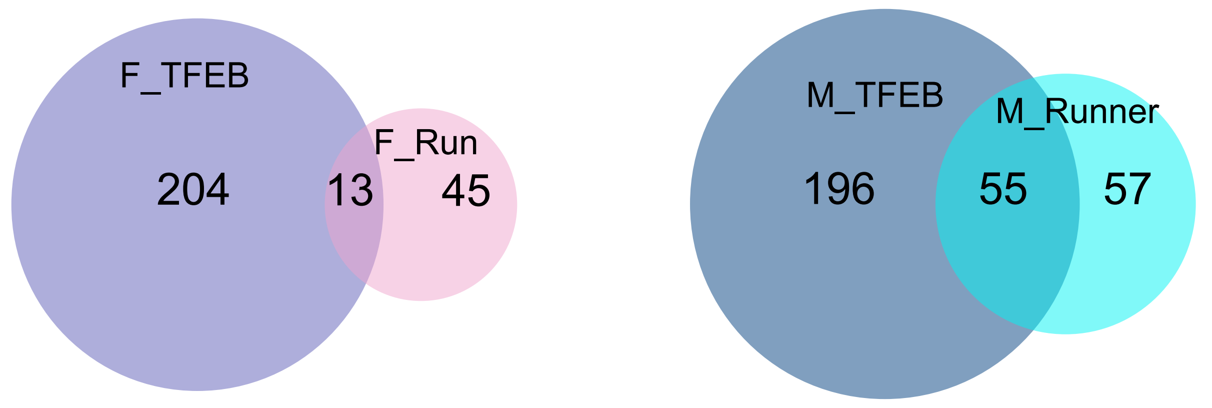


**D.** Top 10 Reactome GSEA gene sets

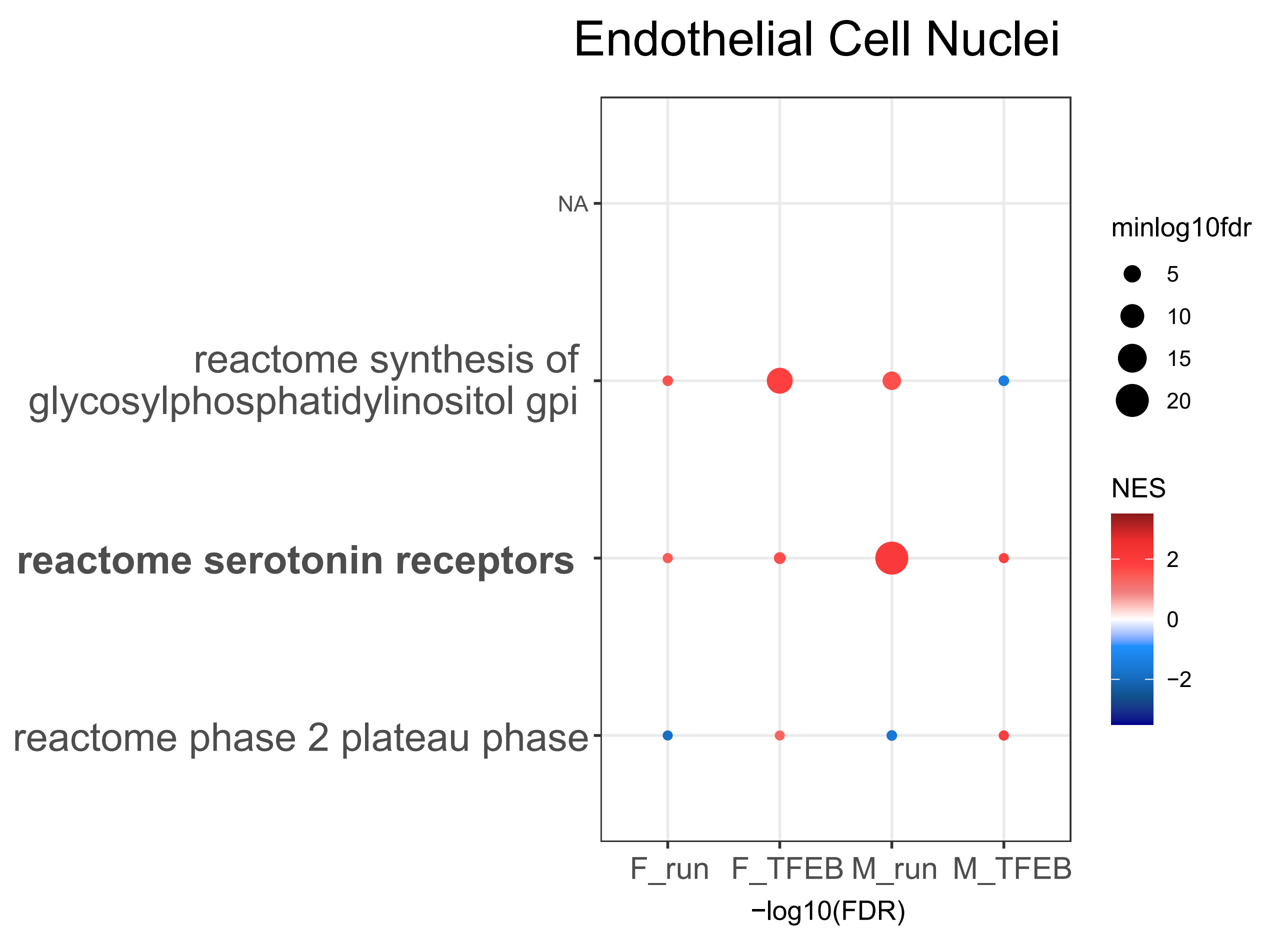


**E.** Representative images of CD31 staining

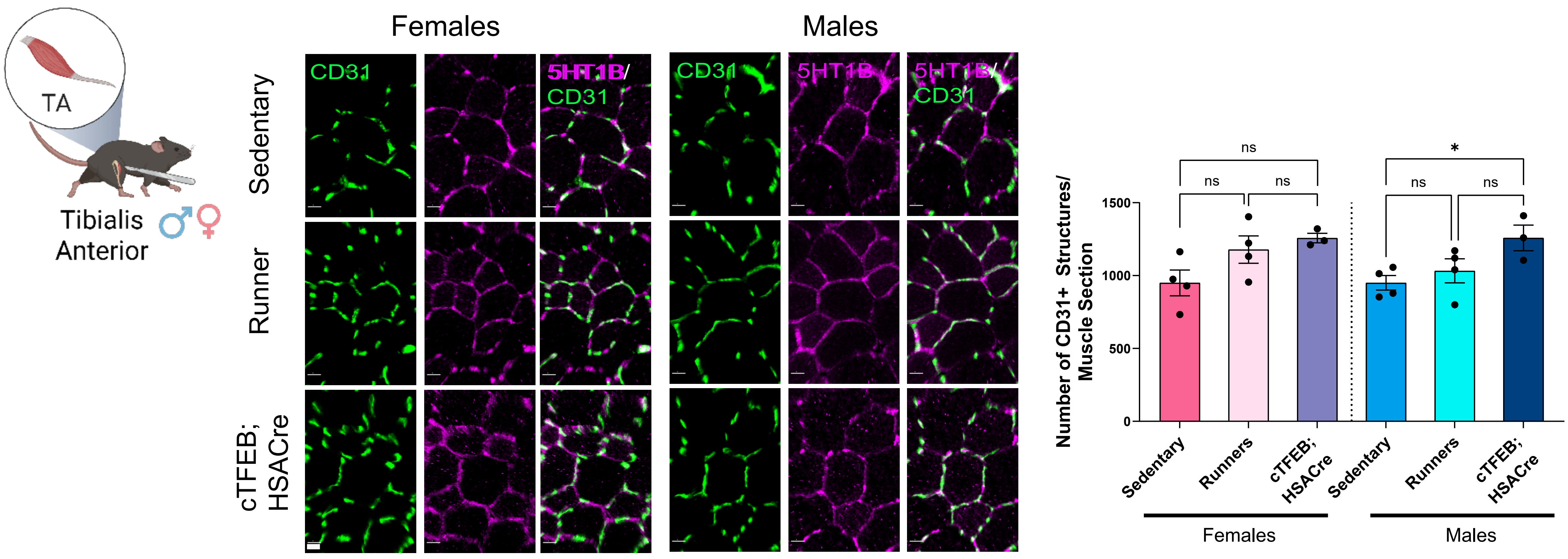

#### SUPPLEMENTARY FIGURE LEGENDS

**Supplementary Figure S1: Running performance and single-nuclei dataset characteristics. (A)** Total and **(B)** average distance run through the course of the 4 weeks of the intervention showed significant differences in female vs. male runners. T-test **(C)** Minor differences in total body weight at collection for female cTFEB;HSACre transgenic mice relative to sedentary controls. One-way ANOVA within sex with post hoc test. **(D)** Confirmation of TFEB transgene expression only in cTFEB;HSACre transgenic samples. **(E)** Confirmation of sex identity by expression of *Xist* (females) and *Ddx3y* (males) in corresponding groups. (**F-H**). UMAP of nuclei colored by sex **(F)**, intervention **(G),** and sex x group **(H)**. **(I)** nuclei annotation pipeline.

**Supplementary Figure S2: Extended fiber typing immunohistochemistry and cross-sectional area quantification across sexes and interventions. (A)** Whole muscle cross-sections of sedentary, runner and cTFEB;HSACre transgenic mice of both sexes stained for fiber typing.. MyHC1 (fiber type I) in red, MyHC2A (fiber type IIa) in green, MyHC2B (fiber type IIb in magenta, MyHC2X (fiber type IIx) unstained, and laminin in blue. Scale bars = 800 µm. **(B)** Fiber cross-sectional area distribution curves for each fiber type for each sex and group, as indicated. One-way ANOVA within sex with Kruskal-Wallis post-hoc test. * Statistical difference between sedentary and runner. # Statistical difference between sedentary and cTFEB;HSACre. Each data point represents the average of three separate images collected from three sections/individual. * $P \leq 0.05$, ** $P \leq 0.01$. Lack of annotation indicates comparisons were not significant. Data is represented as mean ± SD.

**Supplementary Figure S3: Global transcriptional convergence of voluntary running and TFEB activation by cell type and sex. (A-F)** MDS plot of single nuclei transcriptomes across myonuclei and endothelial cells for both sexes. Each point represents one combined sample, and they cluster strongly by intervention (sedentary, runner or TFEB-status). **(G-L)** Correlation Rho maps of global gene expression log2 fold change in running/sedentary vs. TFEB/sedentary nuclei for each cell type and sex. All nuclei types have significant transcriptional overlap between runner and TFEB-expressing samples.

**Supplementary Figure S4: Extended IIa myonuclei GO/Reactome pathway analysis and validation. (A)** Top GO terms enriched by running or TFEB-overexpression in IIa myonuclei. The top 5 GO terms that are significantly down-regulated (top, blue) and the top 5 terms that are significantly enriched across conditions (bottom, red) are shown and ordered by significance. **(B)** Unique and shared differentially expressed GO Biological Process pathways between runner and cTFEB;HSACre IIa myonuclei by sex. **(C)** Unique and shared differentially expressed Reactome pathways between runner and cTFEB;HSACre IIa myonuclei by sex. **(D)** Top eight Reactome terms enriched in at least one condition highlighting shared and divergent transcriptional signatures across sex and intervention. Note the sex-dimorphic enrichment for lipid-associated gene sets in runner and TFEB samples in bold. **(E)** Immunoblot analysis for glucocorticoid receptor (GR) in sedentary, runner, and cTFEB;HSACre tibialis anterior lysates from mice of both sexes (n=3-4/group, independent cohort). Marker densitometry quantification relative to tubulin (loading control) shows no changes in TA muscle from runner and cTFEB;HSACre mice of both sexes. One-way ANOVA within sex with post-hoc test. **(F) T**argeted bulk RNA-Seq analysis confirms reduced expression of NR3C1 (gene encoding for GR) in male (but not female) endurance-trained rat gastrocnemius muscle (MoTrPAC DataHub). **(G, J)** Total levels of ceramides **(G)** and sphingomyelin **(J)** in sedentary, runner, or cTFEB;HSACre tibialis anterior muscle (n=4/group, independent cohort). **(H)** Targeted lipidomics of mouse tibialis anterior muscle showing mild trends towards increased (both female groups), and reductions or no changes (male groups) runners) in detected ceramide (CER) species. One-way ANOVA within sex with Kruskal-Wallis post-hoc test. **(I) T**argeted analysis of ceramides confirms sex-specific CER adaptations to endurance training in rat gastrocnemius muscle of both sexes( MoTrPAC DataHub). Lipid levels reported as logFC over sex-matched sedentary controls. % statistical difference of endurance training as reported by MoTrPAC. N.s: non-significant, * P ≤ 0.05, ** P ≤ 0.01. Lack of annotation indicates comparisons were not significant. Data is represented as mean ± SEM unless otherwise noted.

**Supplementary Figure S5: Extended IIb myonuclei GO/Reactome pathway analysis and validation. (A)** Top GO terms enriched by running or TFEB-overexpression in IIb myonuclei. The top 5 GO terms that are significantly down-regulated (top, blue) and the top 5 terms that are significantly enriched across conditions (bottom, red) are shown and ordered by significance. **(B)** Unique and shared differentially expressed GO Biological Process pathways between runner and cTFEB;HSACre IIb myonuclei by sex. **(C)** Unique and shared differentially expressed Reactome pathways between runner and cTFEB;HSACre IIb myonuclei by sex. **(D)** Top eight Reactome terms enriched in at least one condition highlight shared and divergent transcriptional signatures across sex and intervention. Note the sex-dimorphic enrichment for extracellular membrane (in bold) and neuronal-associated pathways (in italics). **(E)** Cytokine array show clear trends towards sex-specific increases in Tissue Inhibitor of Metalloproteinases 1 (TIMP-1) protein levels in male (but not female) endurance-trained and TFEB-overexpressing quadriceps muscle lysates. One-way ANOVA within sex with post-hoc test. **(F-H)** MoTrPAC DataHub targeted analysis confirms transcriptional effects of endurance training on integrin isoforms *(ITGA4, ITGA6*, and *ITGA7*) on male (but not female) gastrocnemius muscle. All MoTrPAC results are reported as logFC over sex-matched sedentary controls and analyzed as described. % statistical difference of endurance training as reported by MoTrPAC. $ Statistical difference between trained and endurance groups within each sex as reported by MoTrPAC. Lack of annotation indicates comparisons were not significant. N.s: non-significant. Data is represented as mean ± SEM unless otherwise noted.

**Supplementary Figure S6: Extended IIx myonuclei GO/Reactome pathway analysis and validation. (A)** Top GO terms enriched by running or TFEB-overexpression in IIx myonuclei. The top 5 GO terms that are significantly down-regulated (top, blue) and the top 5 terms that are significantly enriched across conditions (bottom, red) are shown and ordered by significance. **(B)** Unique and shared differentially expressed GO Biological Process pathways between runner and cTFEB;HSACre IIx myonuclei by sex. **(C)** Unique and shared differentially expressed Reactome pathways between runner and cTFEB;HSACre IIx myonuclei by sex.

**(D)** Top eight Reactome terms enriched in at least one condition highlight shared and divergent transcriptional signatures across sex and intervention. Mitochondrial associated categories highlighted in bold. **(E)** Total levels of lipid species showing trends towards sex-specific effects in acyl-carnities (CAR) (trending higher in both male samples), **(F)** and free fatty acids (FFA) (trending higher in both female groups), or no changes in either sex fatty acyl chains in tryglycerides (FA) **(G)**, and 4-hydroxynonenal (4-HNE) **(H)** in sedentary, runner, or cTFEB;HSACre tibialis anterior muscle of both sexes. All MoTrPAC results are reported as logFC over sex-matched sedentary controls and analyzed as described. One-way ANOVA within sex with post-hoc test. N.s: non-significant. Data is represented as mean ± SEM unless otherwise noted.

**Supplementary Figure S7: Extended endothelial cell nuclei GO/Reactome pathway analysis and validation. (A)** Top GO terms enriched by running or TFEB-overexpression in endothelial cell (EC) nuclei. The top 5 GO terms that are significantly down-regulated (top, blue) and the top 5 terms that are significantly enriched across conditions (bottom, red) are shown and ordered by significance. **(B)** Unique and shared differentially expressed GO Biological Process pathways between runner and cTFEB;HSACre EC nuclei by sex. **(C)** Unique and shared differentially expressed Reactome pathways between runner and cTFEB;HSACre EC nuclei by sex. Serotonin-related gene sets are highlighted in bold. **(D)** Top 3 Reactome terms enriched in at least one condition highlight shared and divergent transcriptional signatures across sex and intervention. **(E)** Tibialis anterior cross-sections from sedentary, runner and cTFEB;HSACre mice of both sexes (n=3-4/group) stained for CD31 (green), 5HT1B (magenta). Quantification of total CD31+ and 5HT1B+ structures/section for both sexes (right). One-way ANOVA within sex with post-hoc test. N.s: non-significant. Data is represented as mean ± SEM. Scale bars = 15 μm.
